# Engineering DNA templated nonribosomal peptide synthesis

**DOI:** 10.1101/2020.07.27.223297

**Authors:** Hsin-Mei Huang, Philipp Stephan, Hajo Kries

## Abstract

Nanocontainers or macromolecular scaffolds for artificial biocatalytic cascades facilitate sequential enzyme reactions but diffusive escape of intermediates limits rate enhancement. Nonribosomal peptide synthetases (NRPS) naturally form gigantic assembly lines and prevent escape by covalently tethering intermediates. Here, we have built DNA-templated NRPS (DT-NRPS) by adding zinc finger tags to split NRPS modules. The zinc fingers direct the NRPS modules to 9-bp binding sites on a DNA strand, where they form a catalytically active enzyme cascade. DT-NRPS outperform previously reported DNA templated enzyme cascades in terms of DNA acceleration which demonstrates that covalent intermediate channeling is possible along the DNA template. Attachment of assembly line enzymes to a DNA scaffold is a promising catalytic strategy for the sequence-controlled biosynthesis of nonribosomal peptides and other polymers.

## Introduction

Enzymes work in complex metabolic networks that are often spatially subdivided into cascades of successively acting catalysts. In natural or engineered nanoreactors, confined space enhances the concentration of intermediates boosting catalytic performance.^1–4^ Similarly, fusion proteins and co-immobilization of enzyme cascades on DNA, protein or polymer scaffolds have been explored as means to streamline biocatalytic reaction sequences.^5–11^ However, confinement of intermediates rather than co-localization of active sites leads to rate enhancement when intermediates escape diffusively.^10,12^ Therefore, assembly line synthetases for natural products, such as nonribosomal peptide synthetases (NRPS) and polyketide synthases (PKS), harness covalent tethers for more efficient channeling.^13^

To orchestrate long reaction sequences, chemists have developed DNA-templated chemical synthesis.^14–16^ In addition to the rate enhancement caused by reactant proximity, codes written into DNA can control the order in which monomers are connected. For instance, ssDNA templates have been used to build peptide libraries by successively recruiting and connecting ssDNA-tagged monomeric building blocks through complementary annealing. While DNA templated chemical synthesis can utilize a large repertoire of building blocks, (bio)catalytic activation and condensation of unprotected monomers remain elusive.

Here, we engineer DNA templated NRPS (DT-NRPS) to achieve biocatalytic DNA-templated peptide assembly from unprotected building blocks. In natural NRPSs, one amino acid per module is selected in the binding pocket of an adenylation (A) domain, activated and tethered to a thiolation (T) domain. Condensation (C) domains catalyze peptide bond formation between neighboring C-A-T modules, which is a typical domain architecture for one-residue elongation (Figure 1). In addition to the large repertoire of A domains with ca. 500 different substrates reported,^17–19^ tailoring domains such as epimerization (E) domains and alternative product release mechanisms by thioesterase (TE) domains generate astounding structural complexity.^20,21^

**Figure 1.**
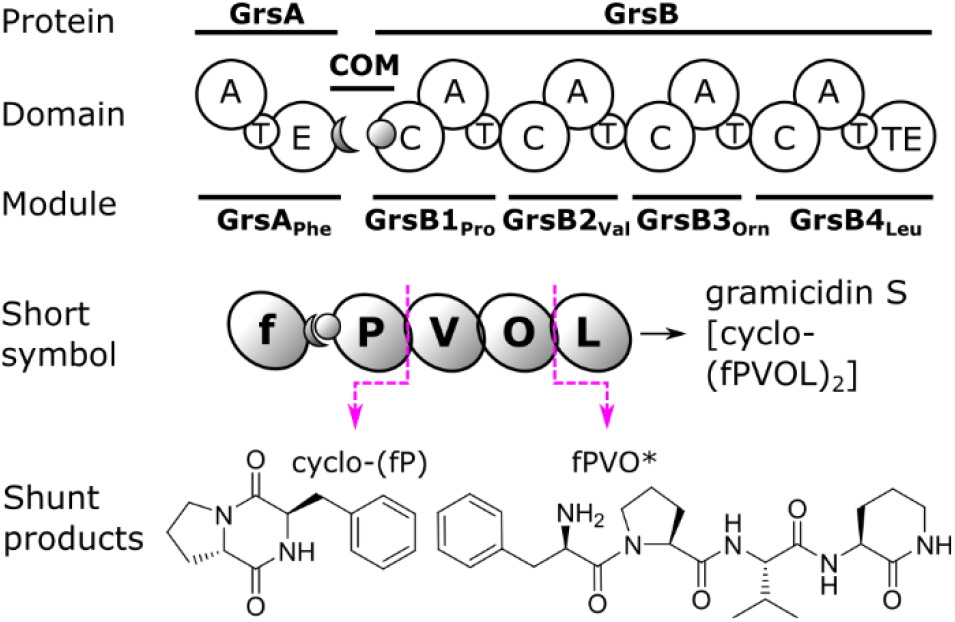
Nonribosomal peptide synthesis. Proteins GrsA and GrsB forming the gramicidin S synthetase interact via a pair of communication mediating domains (COM; grey crescent and circle) and synthesize peptides of different lengths. A, T, C, E, and TE domains (white circles) are grouped into modules (grey ovals) with corresponding substrates, in short symbols, shown in bold-faced single-letter-code.

## Results

Inspired by previous enzyme cascades on DNA scaffolds,^6^ we have built DT-NRPS in three steps (Figure 2). Zinc fingers (ZF) recognizing 9-bp DNA motifs are fused to free standing NRPS modules to confer DNA affinity. The engineering concept relies on a combination of tight, specific DNA binding and weak, unspecific intermodular interactions. We first added ZF tags to NRPS modules, second, engineered low-affinity docking domains for unspecific intermodular communication and third, optimized the intermodular spacer length within the template DNA. Modules for DT-NRPS were recruited by splitting the gramicidin S synthetase (Figure 1). Gramicidin S is a well-investigated cyclic decapeptide antibiotic with the sequence cyclo-(fPVOL)_2_.^22–24^ Mono-modular GrsA (f) and tetramodular GrsB (PVOL) concatenate two molecules of pentapeptide fPVOL into gramicidin S. Here, in addition to the standard abbreviation of amino acids, letters in small case designate D-amino acids, “O” ornithine, and “O*” cyclized ornithine. Interruption of the gramicidin S assembly process leads to shunt products, for instance cyclo-(fP)^25^ and fPVO* (Figure 1).^26^

**Figure 2.**
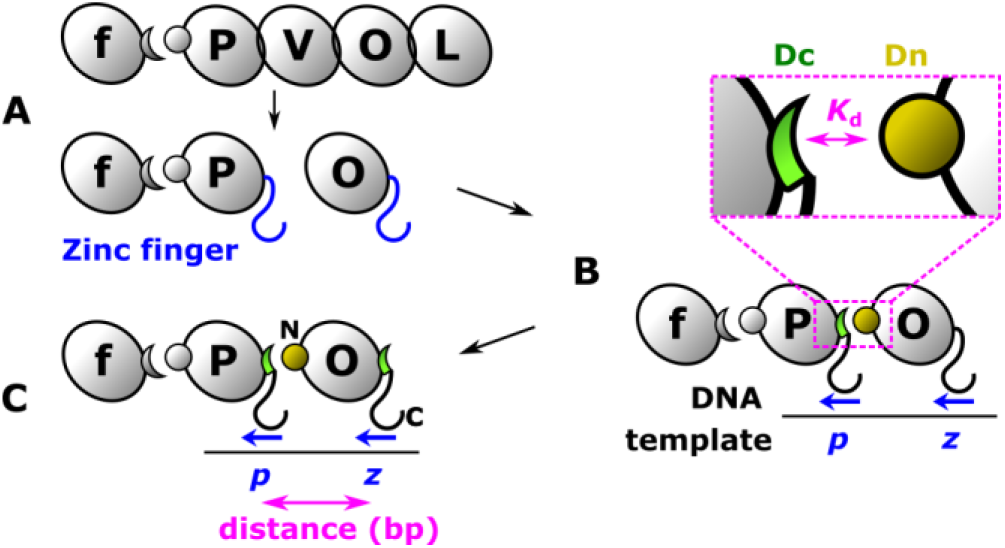
DNA templated NRPS (DT-NRPS) are established in three steps. (A) Zinc finger domains are fused to NRPS modules. (B) Interaction of docking domains (Dc and Dn) is tuned. (C) The DNA spacer between binding sites is optimized. The ZF binding sites (blue italic) are directional (blue arrow), with arrow heads pointing from C- to N-terminus of the ZF domain.

To connect all four modules of GrsB to DNA, they were split apart and equipped with ZF domains. The initiation module GrsA interacts with GrsB via a non-covalent COM domain contact which we kept unmodified. Instead of GrsB1, we used TycB1_Pro_^27^ as a second module – a functionally identical, better expressed homologue from tyrocidine synthesis. As ZF tags, we used PbsII (P) and Zif268 (Z) domains previously employed by Conrado et al. in enzyme cascades converting freely diffusing substrates, where up to 5-fold DNA enhancement was observed.^6,28–30^ Additionally, we employed the nuclear receptor element (N)^31^ and ZFB (B) engineered for the target site GGG-GCT-GCG with a design tool.^32^ With these ZF, we tagged NRPS modules TycB1-N, GrsB2-P, GrsB3-Z, Z-GrsB3 and B-GrsB4, where “N”, “P”, “Z”, and “B” designate ZF identity and position. For these proteins, DNA affinities were measured via fluorescence polarization assays employing fluorescein-dT labeled DNA templates (Table S4). The ZF domains convey specific binding of DNA recognition sequences to the NRPS modules with nanomolar dissociation constants (Figure 3).^33^ Since the ZF used here require three Zn^2+^ ions each for DNA binding and NRPS require Mg^2+^ for catalysis, we tested the effect of Zn^2+^ and Mg^2+^ on cyclo-(fP) formation of GrsA/TycB1 (Figure S1). A good compromise was found at 0.1 mM Mg^2+^ and 10 μM Zn^2+^, where Zn^2+^ has no inhibitory effect but will be able to saturate several ZF proteins added at micromolar concentration.

**Figure 3.**
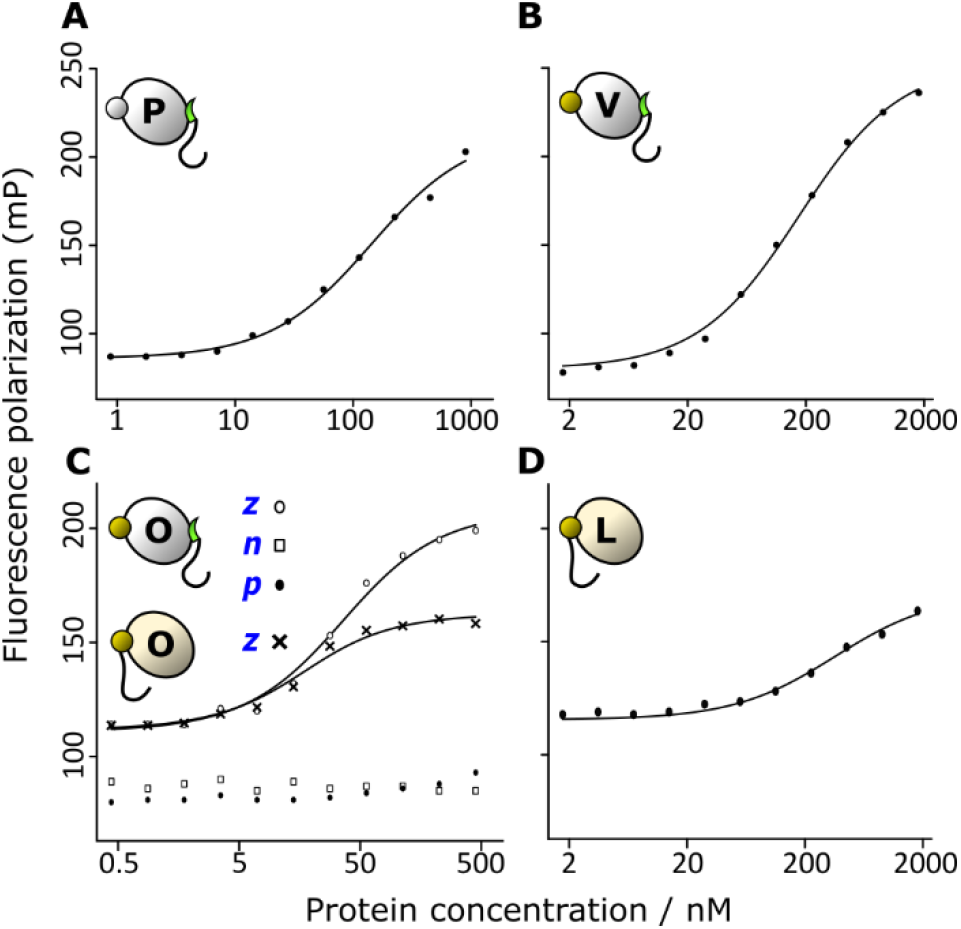
Affinity of ZF to DNA. Fluorescence polarization data were fitted to a bimolecular binding model to determine the dissociation constants (*K*_d_) between modules and ZF binding sites. (A) TycB1-N (140 ± 10 nM). (B) GrsB2-P (170 ± 10 nM), (C) GrsB3-Z (31 ± 4 nM) and Z-GrsB3 (12 ± 2 nM), (D) B-GrsB4 (350 ± 50 nM). A yellow shade indicates an N-terminal ZF. All modules were tested with their cognate ZF binding site, but module GrsB3-Z (bottom left) was also tested with noncognate binding sites *n* and *p*.

The first design target was a trimodular DT-NRPS with one DNA-independent and one DNA-dependent module connection that would produce the non-natural peptide fPO*. This peptide can be released from the synthetase via ornithine side-chain cyclisation (Figure 4a).^26^ GrsA and TycB1-P are connected via a native, non-covalent COM domain interaction. Between the DNA bound modules TycB1-P and GrsB3-Z, tuned intermodular affinities must ensure productive interaction while minimizing undesired DNA-independent activity. Artificial splitting of NRPS proteins into modules and reconnection with docking domains has yielded good amounts of protein and high peptide yields upon heterologous co-expression before.^34^ Therefore, we retrieved C-terminal (Dc) and N-terminal (Dn) docking domains mediating module interactions in xenortide biosynthesis from InxAB (InxA-Dc/Dn-InxB).^35,36^ These docking domains are fully portable to the gramicidin S NRPS modules. In a disconnected, trimodular system for fPO* formation (GrsA, TycB1, GrsB3) a Dc/Dn domain pair added to connect the second and third module increases activity more than 70-fold compared to modules without docking domains (Figure S2).

**Figure 4.**
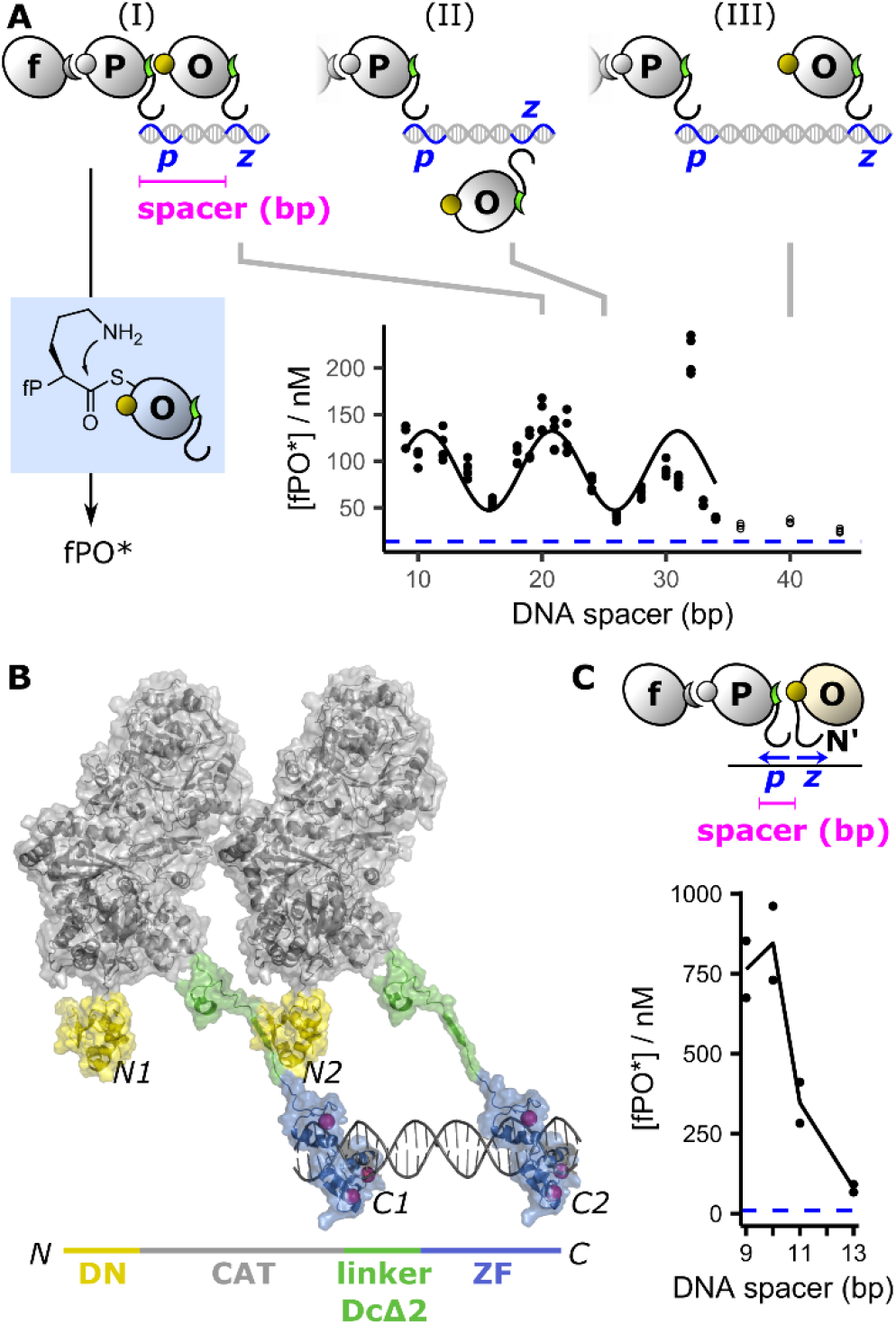
Adjustment of the DNA spacer in a trimodular NRPS. (A) One free and two DNA-bound modules synthesize tripeptide fPO* and the spacer between modules two and three is optimized. The spacer length includes the 9 bp ZF binding site and a random sequence of variable length (Table S4). Blue dashed line: no DNA. Reaction conditions: 0.4 μM GrsA, 0.1 μM TycB1-P, 0.5 μM GrsB3-Z, and 0.4 μM DNA template. At least three data points were collected per distance with two batches of protein. Data points up to 34 bp (closed circles) were nonlinearly fit in R to a sine wave with a wavelength of 10.1±0.3 bp. Three scenarios for the positioning of DT-NRPS modules along the DNA helix: (I) favourable helical angle, (II) unfavourable helical angle, and (III) favourable helical angle but unfavourable spacer length. (B) A two-modular DT-NRPS was homology modelled (see Supplementary Methods) on an NMR structure of the docking domain complex (Dc: green; Dn: yellow)^36^ and crystal structures of an NRPS module (grey)^37^ and a ZF (protein in blue, Zn atoms in pink, DNA in grey).^38^ (C) In the third module (yellow shade), the ZF is moved to the N-terminus and the DNA spacer is optimized between the C- and N-terminal ZF. Reaction conditions: 0.4 μM GrsA, 0.1 μM TycB1-P, 0.5 μM Z-GrsB3, and 0.4 μM DNA template with replicates from two batches of protein.

For tuning the docking domain (Dc-Dn) interaction to DT-NRPS requirements, modification of the C-terminal residues of the Dc domain seemed promising based on mutational and biophysical data.^36^ We prepared a series of truncated versions of Dc and tested activity in the presence and absence of DNA. The truncated docking domains were integrated between NRPS module and ZF (Tables S1 and S3). TycB1-P carrying a two-residue docking domain truncation (DcΔ2) showed a good compromise between DNA enhancement and overall activity (Figure S3). Hence, the DcΔ2 truncation was used in all ZF tagged modules. Measurements with an isothermal titration calorimeter (ITC) detect weak interaction between Dn-tagged GrsB3 and the C-terminal octapeptide of Dc (Figure S4). Additional C-terminal residues also present when a ZF follows the Dc domain weaken binding, suggesting beneficial electrostatic effects of a free C-terminus. No interaction is detectable by ITC after two-residue truncation, confirming the desired lowering of binding affinity.

The rigid double-stranded DNA template with predictable molecular dimensions served as a ruler to probe the structural requirements of the floppy NRPS modules attached to it. In crystal structures of the linear gramicidin synthetase A, module dimensions of 85-216 Å have been measured in highly variable conformations.^37^ To optimize intermodular spacing, we tested a range of spacers between the ZF recognition sites that would hold the NRPS modules at approximately these distances. Strikingly, with the trimodular system for fPO* formation, we observed a periodic pattern with activity maxima at 10, 20 and 32 bp distance (Figure 4a). We interpret the sinusoidal activity pattern as an effect of the DNA helical angle because it has a wavelength of 10.1±0.3 bp similar to the length of a turn in B-DNA (10.5 bp). The NRPS modules are either on the same (maxima) or on opposite sides of the helix (minima). The maximum at 32 bp (≙ 109 Å) is a positive outlier which might imply a higher fragility of the architecture at this length, arguing in favor of 20 bp (≙ 68 Å; scenario I) as a default spacer. A spacer of 20 bp yielded plausible geometries in a homology model (Figure 4b) and was also expected to prevent module skipping since twice this length would be too long for productive interaction (Figure 4a, scenario III). In agreement with a generally high flexibility of the DT-NRPS, inverting the direction of the ZF binding sites in the DNA template sequence only reduces the fPO* yield by 40 ± 2 % (Figure S5).

Then, we tested with construct Z-GrsB3 whether a ZF domain could also be fused to the N-terminus. With a pair of modules connected head-to-tail at C- and N-terminus, the distance optimum for the DNA ‘staple’ reinforcing the docking domain was observed at 10 bp (Figure 4c). More fPO* product is obtained with this architecture compared to the head-to-head construct (Figure 4a) and DNA accelerates the reaction by a factor of up to 90 ± 30 but application seems limited to terminal modules. Internal modules would have to be ZF-tagged on both ends, leading to additional complications.

We validated the optimized spacer length with two adjacent spacers by testing modules TycB1-N, GrsB2-P, and GrsB3-Z on templates with 9, 20, and 32-bp spacing (Figure 5a). To compare with the natively fused, first three modules of GrsB (GrsB123), TycB1-N which causes non-templated cyclo-(fP) formation was supplied at limiting concentration. Comparison with the cyclo-(fP) concentration corrects for the imperfect purity of the covalently linked control protein GrsB123 with a size of 358 kDa (Figure S6). Addition of a 20-bp spacer template enhances the [fPO*]/[fP] ratio by a factor of 40 ± 10. Hence, the DT-NRPS reaches one third of the productivity of natively fused GrsB123 and higher DNA enhancement than previously reported artificial enzyme cascades relying on freely diffusing intermediates.^6^ A 20-bp spacing reduces formation of byproduct fPO* compared to 9 bp and is almost twice as efficient as 32 bp, confirming that 20-bp spacing is preferable.

**Figure 5.**
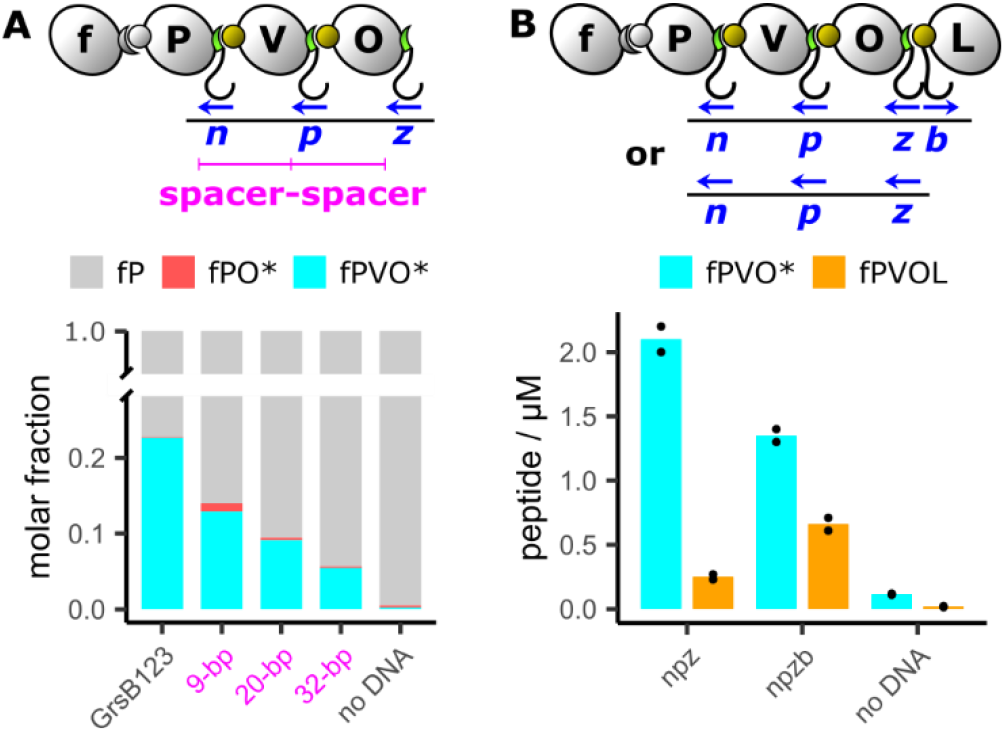
Multimodular templating. (A) Optimization of spacers between three DNA-bound modules and comparison to natively-fused, trimodular GrsB123. Production of the desired product fPVO* and the module-skipped product fPO* was quantified with biological duplicates in comparison with chemically synthesized standards (Table S7). Reaction conditions: 0.4 μM GrsA, 0.25 μM GrsB123, no DNA (GrsB123); 0.4 μM GrsA, 0.1 μM TycB1-N, 1 μM GrsB2-P, 1 μM GrsB3-Z, with or without 0.5 μM DNA template (other columns). (B) Effect of full (npzb) or partial (npz) template on tetra- and pentapeptide formation measured with biological duplicates. Reaction conditions: 1 μM of GrsA, TycB1-N, GrsB2-P, GrsB3-Z, and B-GrsB4 with 0.25 μM DNA template.

Addition of module B-GrsB4 completed the gramicidin S synthetase (Figure 5b). The expected fPVOL peptide was formed with the appropriate DNA template at a concentration of 0.7 ± 0.1 μM. However, fPVO* was formed more efficiently (1.4 ± 0.1 μM) and removing the ZF binding site for the last module from the DNA template only reduced fPVOL formation by a factor of 2.6 ± 0.4. Imperfect elongation in the last step might be due to relatively low DNA affinity of the B-GrsB4 construct (Figure 3). These results indicate potential for optimization of DT-NRPS through recruitment of ZF or other DNA binding domains with higher affinity.

To confirm the supramolecular structure, we measured apparent sizes of module-DNA complexes by analytical size exclusion chromatography (SEC; Figure 6). Successive addition of GrsB3-Z, GrsB2-P, and TycB1-N revealed accelerated elution, suggesting that multimodular assembly occurs mostly as intended.

**Figure 6.**
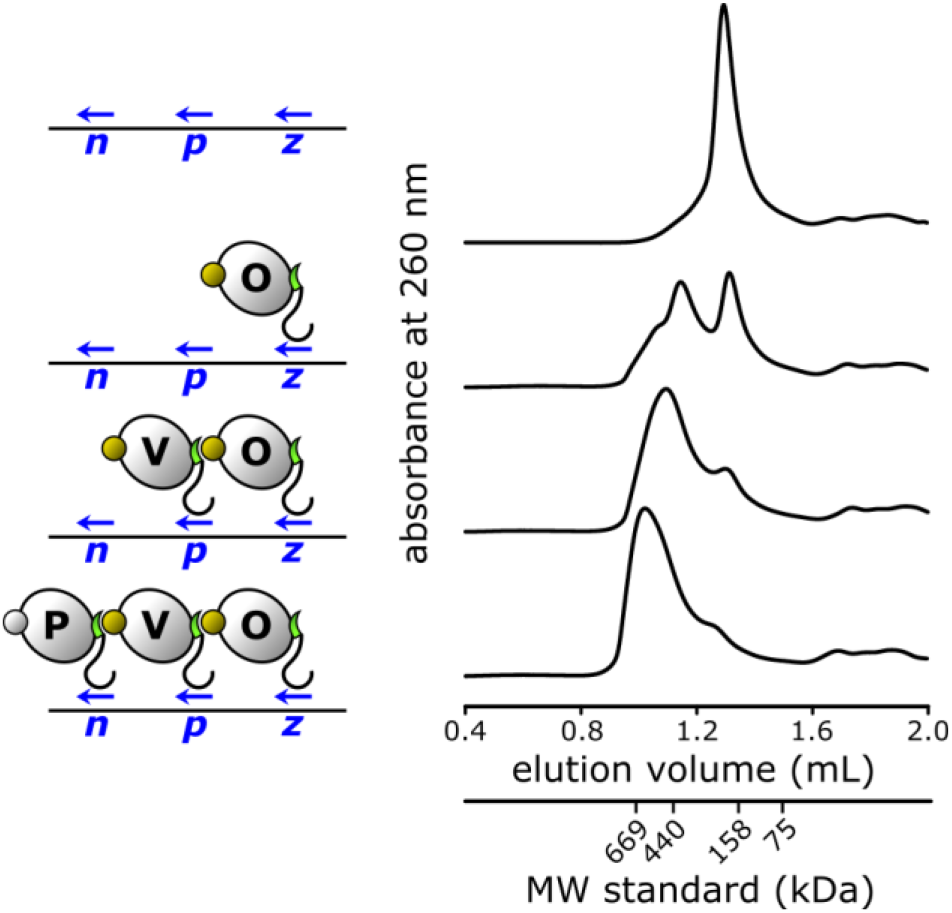
Investigation of the supramolecular assembly. Analytical size exclusion chromatography was performed at 2.5 μM of each protein and 2 μM template DNA. Molecular weight (MW) standards: 669 kDa thyroglobin, 440 kDa ferritin, 158 kDa aldolase, and 75 kDa conalbumin.

## Discussion

With the DNA-directed assembly of a pentamodular DT-NRPS, we are spearheading a novel catalytic concept for DNA-templated peptide synthesis relying on biocatalysts and unprotected amino acid building blocks. High DNA-enhancement factors demonstrate the advantage of covalent intermediate capture in biocatalytic cascade reactions. NRPS modules are mechanistically far more complex than single-domain enzymes previously assembled on DNA, for instance horseradish peroxidase and glucose oxidase, and intermediate tethering comes with the additional challenge to hand over the intermediate to the next module in the right orientation. Nonetheless, NRPS are not only active on DNA but also show a remarkable tolerance for different structural arrangements (Figures 4 and 5). The oscillating activity upon spacer elongation shows a strong influence of the helical angle between the NRPS modules up to a distance cut-off, where they can no longer assume the compact conformation required for peptide formation.^37^ The generally high tolerance of the DT-NRPS for different spatial arrangements is consistent with the vast conformational flexibility of NRPS modules observed in crystallography. This flexibility augurs well for the integration of non-canonical modules performing formylation, methylation, cyclization, or oxidation, without re-optimization of linkers, spacers, and docking domains.

DT-NRPS open up new perspectives for combinatorial NRPS engineering which receives broad attention since nonribosomal peptides such as actinomycin D, vancomycin, or cyclosporin A are clinically used drugs.^20,21,39,40^ When megaenzymes are spliced or shuffled, every design requires time-consuming manipulation of large gene clusters, limiting combinatorial freedom. In contrast, DT-NRPS form the desired module sequence in situ directed by the DNA template. Instead of ~3 kb per module gene, DT-NRPS only require 20 bp “codons” containing a 9 bp ZF binding site and a spacer possibly useful for binding site recombination. The successful combination of modules TycB1 and GrsB3 gives a first indication that DNA can enforce non-native module interactions. Not all module interactions must rely on DNA. Here, we have left the GrsA/TycB1 interaction in the native state to generate a non-templated control peptide. Especially in the re-design of complex, non-linear architectures, parts of the enzyme can also remain non-templated.

DT-NRPS showcases a catalytic concept with implications beyond peptide synthesis. Next to NRPS, polyketide synthases are in nature’s repertoire of intermediate-tethering assembly line enzymes, which may be suited to join DNA-assembled machineries, too. In the future, enzyme design and engineering can integrate various modules into DNA templated cascades to achieve programmable, biocatalytic synthesis of multifarious polymers.

## Acknowledgments

We gratefully acknowledge donation of plasmids pSU18-grsA, pTrc99a-tycB1, pTrc99a_grsB_M3574L and *E. coli* strain HM0079 by Donald Hilvert (ETH Zurich) under a material transfer agreement. We thank Sarah E. O’Connor and Clemens Mayer for helpful discussions, Daniela Winkler for help with chemical synthesis and Yan Li for determining high resolution masses. HK gratefully acknowledges a fellowship of the Daimler und Benz Foundation. This work was financially supported by the Leibniz Society, the *Fonds der Chemischen Industrie* (FCI) and the BMBF (ZIK Septomics, FKZ 03Z22JI1).

## Supplementary Information

Complete experimental procedures, additional data, Figures S1-S6, Tables S1-S7 and NMR spectra (PDF)

## References

(1) Bernhardsgrütter, I.; Vögeli, B.; Wagner, T.; Peter, D. M.; Cortina, N. S.; Kahnt, J.; Bange, G.; Engilberge, S.; Girard, E.; Riobé, F.; et al. The Multicatalytic Compartment of Propionyl-CoA Synthase Sequesters a Toxic Metabolite. Nat. Chem. Biol. 2018, 14 (12), 1127–1132.

(2) Wheeldon, I.; Minteer, S. D.; Banta, S.; Barton, S. C.; Atanassov, P.; Sigman, M. Substrate Channelling as an Approach to Cascade Reactions. Nat. Chem. 2016, 8 (4), 299–309.

(3) Brasch, M.; Putri, R. M.; De Ruiter, M. V.; Luque, D.; Koay, M. S. T.; Castón, J. R.; Cornelissen, J. J. L. M. Assembling Enzymatic Cascade Pathways inside Virus-Based Nanocages Using Dual-Tasking Nucleic Acid Tags. J. Am. Chem. Soc. 2017, 139 (4), 1512–1519.

(4) Vázquez-González, M.; Wang, C.; Willner, I. Biocatalytic Cascades Operating on Macromolecular Scaffolds and in Confined Environments. Nat. Catal. 2020, 3 (3), 256–273.

(5) Fu, J.; Liu, M.; Liu, Y.; Woodbury, N. W.; Yan, H. Interenzyme Substrate Diffusion for an Enzyme Cascade Organized on Spatially Addressable DNA Nanostructures. J. Am. Chem. Soc. 2012, 134 (12), 5516–5519.

(6) Conrado, R. J.; Wu, G. C.; Boock, J. T.; Xu, H.; Chen, S. Y.; Lebar, T.; Turnek, J.; Tomšič, N.; Avbelj, M.; Gaber, R.; et al. DNA-Guided Assembly of Biosynthetic Pathways Promotes Improved Catalytic Efficiency. Nucleic Acids Res. 2012, 40 (4), 1879–1889.

(7) Kang, W.; Ma, T.; Liu, M.; Qu, J.; Liu, Z.; Zhang, H.; Shi, B.; Fu, S.; Ma, J.; Lai, L. T. F.; et al. Modular Enzyme Assembly for Enhanced Cascade Biocatalysis and Metabolic Flux. Nat. Commun. 2019, 10 (1), 4248.

(8) Bugada, L. F.; Smith, M. R.; Wen, F. Engineering Spatially Organized Multi-Enzyme Assemblies for Complex Chemical Transformation. ACS Catal. 2018, 8, 7898–7906.

(9) Aalbers, F. S.; Fraaije, M. W. Enzyme Fusions in Biocatalysis: Coupling Reactions by Pairing Enzymes. ChemBioChem 2019, 20 (1), 20–28.

(10) Zhang, Y.; Hess, H. Toward Rational Design of High-Efficiency Enzyme Cascades. ACS Catal. 2017, 7 (9), 6018–6027.

(11) Xu, X.; Tian, L.; Tang, S.; Xie, C.; Xu, J.; Jiang, L. Design and Tailoring of an Artificial DNA Scaffolding System for Efficient Lycopene Synthesis Using Zinc-Finger-Guided Assembly. J. Ind. Microbiol. Biotechnol. 2020, 47 (2), 209–222.

(12) Bauler, P.; Huber, G.; Leyh, T.; McCammon, J. A. Channeling by Proximity: The Catalytic Advantages of Active Site Colocalization Using Brownian Dynamics. J. Phys. Chem. Lett. 2010, 1 (9), 1332–1335.

(13) Fischbach, M. A.; Walsh, C. T. Assembly-Line Enzymology for Polyketide and Nonribosomal Peptide Antibiotics: Logic, Machinery, and Mechanisms. Chem. Rev. 2006, 106 (8), 3468–3496.

(14) Gartner, Z. J.; Tse, B. N.; Grubina, R.; Doyon, J. B.; Snyder, T. M.; Liu, D. R. DNA-Templated Organic Synthesis and Selection of a Library of Macrocycles. Science 2004, 305 (5690), 1601–1605.

(15) Meng, W.; Muscat, R. A.; McKee, M. L.; Milnes, P. J.; El-Sagheer, A. H.; Bath, J.; Davis, B. G.; Brown, T.; O’Reilly, R. K.; Turberfield, A. J. An Autonomous Molecular Assembler for Programmable Chemical Synthesis. Nat. Chem. 2016, 8 (6), 542–548.

(16) Goodnow, R. A.; Dumelin, C. E.; Keefe, A. D. DNA-Encoded Chemistry: Enabling the Deeper Sampling of Chemical Space. Nat. Rev. Drug Discov. 2017, 16 (2), 131–147.

(17) Walsh, C. T.; O’Brien, R. V; Khosla, C. Nonproteinogenic Amino Acid Building Blocks for Nonribosomal Peptide and Hybrid Polyketide Scaffolds. Angew. Chem. Int. Ed. 2013, 52 (28), 7098–7124.

(18) Flissi, A.; Ricart, E.; Campart, C.; Chevalier, M.; Dufresne, Y.; Michalik, J.; Jacques, P.; Flahaut, C.; Lisacek, F.; Leclère, V.; et al. Norine: Update of the Nonribosomal Peptide Resource. Nucleic Acids Res. 2020, 48 (D1), D465–D469.

(19) Stanišić, A.; Kries, H. Adenylation Domains in Nonribosomal Peptide Engineering. ChemBioChem 2019, 20 (11), 1347–1356.

(20) Süssmuth, R. D.; Mainz, A. Nonribosomal Peptide Synthesis-Principles and Prospects. Angew. Chem. Int. Ed. 2017, 56 (14), 3770–3821.

(21) Winn, M.; Fyans, J. K.; Zhuo, Y.; Micklefield, J. Recent Advances in Engineering Nonribosomal Peptide Assembly Lines. Nat. Prod. Rep. 2016, 33 (2), 317–347.

(22) Krätzschmar, J.; Krause, M.; Marahiel, M. A. Gramicidin S Biosynthesis Operon Containing the Structural Genes GrsA and GrsB Has an Open Reading Frame Encoding a Protein Homologous to Fatty Acid Thioesterases. J. Bacteriol. 1989, 171 (10), 5422–5429.

(23) Berditsch, M.; Afonin, S.; Reuster, J.; Lux, H.; Schkolin, K.; Babii, O.; Radchenko, D. S.; Abdullah, I.; William, N.; Middel, V.; et al. Supreme Activity of Gramicidin S against Resistant, Persistent and Biofilm Cells of Staphylococci and Enterococci. Sci. Rep. 2019, 9 (1), 17938.

(24) Guan, Q.; Huang, S.; Jin, Y.; Campagne, R.; Alezra, V.; Wan, Y. Recent Advances in the Exploration of Therapeutic Analogues of Gramicidin S, an Old but Still Potent Antimicrobial Peptide. J. Med. Chem. 2019, 62 (17), 7603–7617.

(25) Stachelhaus, T.; Mootz, H. D.; Bergendahl, V.; Marahiel, M. A. Peptide Bond Formation in Nonribosomal Peptide Biosynthesis. Catalytic Role of the Condensation Domain. J. Biol. Chem. 1998, 273 (35), 22773–22781.

(26) Mootz, H. D.; Schwarzer, D.; Marahiel, M. A. Construction of Hybrid Peptide Synthetases by Module and Domain Fusions. Proc. Natl. Acad. Sci. USA 2000, 97 (11), 5848–5853.

(27) Mootz, H. D.; Marahiel, M. A. The Tyrocidine Biosynthesis Operon of Bacillus Brevis: Complete Nucleotide Sequence and Biochemical Characterization of Functional Internal Adenylation Domains. J. Bacteriol. 1997, 179 (21), 6843–6850.

(28) Blancafort, P.; Segal, D. J.; Barbas, C. F. Designing Transcription Factor Architectures for Drug Discovery. Mol. Pharmacol. 2004, 66 (6), 1361–1371.

(29) Segal, D. J. The Use of Zinc Finger Peptides to Study the Role of Specific Factor Binding Sites in the Chromatin Environment. Methods 2002, 26 (1), 76–83.

(30) Lemaire, P.; Revelant, O.; Bravo, R.; Charnay, P. Two Mouse Genes Encoding Potential Transcription Factors with Identical DNA-Binding Domains Are Activated by Growth Factors in Cultured Cells. Proc. Natl. Acad. Sci. USA 1988, 85 (13), 4691–4695.

(31) Greisman, H. A.; Pabo, C. O. A General Strategy for Selecting High-Affinity Zinc Finger Proteins for Diverse DNA Target Sites. Science 1997, 275 (5300), 657–661.

(32) Mandell, J. G.; Barbas, C. F. Zinc Finger Tools: Custom DNA-Binding Domains for Transcription Factors and Nucleases. Nucleic Acids Res. 2006, 34 (Web Server), W516–W523.

(33) Jantz, D.; Berg, J. M. Probing the DNA-Binding Affinity and Specificity of Designed Zinc Finger Proteins. Biophys. J. 2010, 98 (5), 852–860.

(34) Kegler, C.; Bode, H. B. Artificial Splitting of a Non-Ribosomal Peptide Synthetase by Inserting Natural Docking Domains. Angew. Chem. Int. Ed. 2020, anie.201915989.

(35) Cai, X.; Nowak, S.; Wesche, F.; Bischoff, I.; Kaiser, M.; Fürst, R.; Bode, H. B. Entomopathogenic Bacteria Use Multiple Mechanisms for Bioactive Peptide Library Design. Nat. Chem. 2017, 9 (4), 379–386.

(36) Hacker, C.; Cai, X.; Kegler, C.; Zhao, L.; Weickhmann, A. K.; Wurm, J. P.; Bode, H. B.; Wöhnert, J. Structure-Based Redesign of Docking Domain Interactions Modulates the Product Spectrum of a Rhabdopeptide-Synthesizing NRPS. Nat. Commun. 2018, 9 (1), 4366.

(37) Reimer, J. M.; Eivaskhani, M.; Harb, I.; Guarné, A.; Weigt, M.; Schmeing, T. M. Structures of a Dimodular Nonribosomal Peptide Synthetase Reveal Conformational Flexibility. Science 2019, 366 (6466), eaaw4388.

(38) Pavletich, N. P.; Pabo, C. O. Zinc Finger-DNA Recognition: Crystal Structure of a Zif268-DNA Complex at 2.1 A. Science 1991, 252 (5007), 809–817.

(39) Kalkreuter, E.; Williams, G. J. Engineering Enzymatic Assembly Lines for the Production of New Antimicrobials. Curr. Opin. Microbiol. 2018, 45, 140–148.

(40) Alanjary, M.; Cano-Prieto, C.; Gross, H.; Medema, M. H. Computer-Aided Re-Engineering of Nonribosomal Peptide and Polyketide Biosynthetic Assembly Lines. Nat. Prod. Rep. 2019, 36 (9), 1249–1261.

