## Supplementary Information for "Engineering DNA templated nonribosomal peptide synthesis"

### Experimental Procedures

#### Plasmid construction

Plasmids are listed in Table S1 and representative sequences are provided in GenBank file format (Table S2). DNA polymerases, T5 exonuclease, and restriction enzymes were obtained from New England Biolabs. Synthetic oligos, gene synthesis, and Sanger sequencing services were provided by Eurofins Genomics (Ebersberg, Germany).

Nonribosomal peptide synthetase (NRPS) modules were cloned from plasmids pTrc99a-*tycB1*,[1] pSU18-*grsA*[2] and pTrc99a_*grsB*_M3574L.[3] Linkers between NRPS module and zinc finger domain are listed in Table S3. Zinc finger domains *zif268* (Z), *pbsII* (P), *nre* (N)[4], and *zfb* (B) were synthesized or cloned from synthetic fragments followed by a C-terminal $His_6$ tag unless otherwise specified. The gene for the N-terminal docking domain (Dn) was cloned from *inxB* of *Xenorhabdus innexi* DSM 16336. The cognate C-terminal docking domain (Dc) was synthesized according to the peptide sequence of *inxA* (GenBank: KR871226)[5] with a conservative mutation from Ile to Leu in order to embed an *Afl*II restriction site.

Recombinant genes were assembled into pSU18 vector ($Chl^R$) using the Hot Fusion cloning method.[6] Chemically competent cells were prepared from *E. coli* HM0079,[1] and transformed following the TSS protocol.[7]

#### Protein expression and purification

Proteins were expressed using *E. coli* HM0079 as host which genomically encodes a phosphopantetheinyl transferase (Sfp) and generates the holo-form of NRPS modules. Media were supplemented with 25 µg/mL chloramphenicol. Bacterial cultures were incubated at 37 °C shaking at 230 rpm with an orbit diameter of 5.1 cm unless otherwise specified.

For protein expression, a preculture in LB broth was inoculated from a single colony of HM0079 freshly transformed with the relevant plasmid and incubated overnight (14-16 h). As growth medium, we used 2YT medium containing 16 g tryptone, 10 g yeast extract, 5 g NaCl in 1 L water adjusted to pH 7.0. A 500 mL Erlenmeyer flask containing 100 mL 2YT media was inoculated with 100 µL preculture and supplemented with sterile 10 µM $ZnCl_2$. The resulting culture was grown to mid-log phase (ca. 4 h, $OD_{600}$ ~ 0.5) and protein expression induced with 0.25 mM IPTG. Incubation was continued at 20 °C for 15-16 h. Bacteria were pelleted by centrifugation at 10,000 x g and 7 °C for 3 min and stored at -20 °C.

All reagents for protein purification were precooled to 4 °C. Procedures were performed below 10 °C except for nickel affinity chromatography. Bacterial pellets were thawed and thoroughly resuspended in one tenth of the original culture volume with buffer A1 (500 mM NaCl, 20 mM imidazole, 2 mM tris(2-carboxyethyl)phosphine [TCEP], 50 mM Tris-HCl, pH 7.4). For cell lysis, up to 25 mL suspension were contained in a 50 mL conical tube cooled on ice. Immediately before cell lysis, 0.02% of the original culture volume of a protease inhibitor cocktail (Cat.No. P8849, Sigma) were added. Ultrasonic lysis was performed for a total of 10 min using an ultrasonic homogenizer UP200St (Hielscher) with sonotrode S26d7 (7 mm diameter) at 40% amplitude and periodic on/off cycles of 2 and 3 s. Cellular debris was removed by centrifugation at 19,000 x g for 30 min. To capture His-tagged protein, the supernatant was applied to 1 mL Rotigarose-His/Ni IDA-agarose beads suspension (Cat.No. 1308.1, Carl Roth) pre-equilibrated with buffer A1 in an Econo-Column (1.5 x 15 cm; Bio-Rad). The suspension was incubated for 12.5 min for resin to settle before draining the lysate. The beads were then washed three times with 6 mL buffer A1 followed by elution with 200 µL buffer B1 (500 mM NaCl, 300 mM imidazole, 2 mM TCEP, 50 mM Tris-HCl, pH 7.4) which were discarded and two times 0.5 mL buffer B1 which contained the protein of interest.

For further purification and to remove contaminating genomic DNA, the protein was subjected to anion exchange chromatography unless specified otherwise. The protein was buffer exchanged using Vivaspin 500 centrifugal filters (30 kDa cut-off, Sartorius) into low salt buffer (20 mM NaCl, 20 mM Tris-HCl, pH 8.0, filtered) and then injected onto a MonoQ 5x50 GL anion exchange column (GE Healthcare) connected to an NGC Quest 10 Plus Chromatography System (Bio-Rad). Elution was performed at a flow rate of 1 mL/min at 10 °C with a linear gradient of 20 to 650 mM NaCl in 20 mM Tris buffer (pH 8.0) in a total volume of 7.85 mL. Fractions containing target protein were pooled, concentrated, and exchanged into DKP-M buffer (100 mM NaCl, 1 mM TCEP, 50 mM HEPES, pH 8.0) using Vivaspin 500 centrifugal filters.

Protein concentrations were determined spectrophotometrically using calculated extinction coefficients. Of each protein, two 2.5 µL aliquots were loaded onto a Take3 Plate (Biotek) and absorbance at 280 nm determined on an Epoch2 microplate reader (Biotek) equipped with Gen5 software (Biotek). The purified protein was adjusted to a concentration of 50 µM in DKP-M buffer, supplemented with 10% glycerol, frozen in liquid nitrogen, and stored at -20 °C until further assays were performed. Protein purity was verified by SDS-PAGE (Figure S6).

### Preparation of template DNA for peptide synthesis

DNA templates for peptide production assays were prepared by annealing synthetic oligos (Eurofins Genomics; Table S4). The oligos were dissolved in $H_2O$ and annealing reactions were performed in buffer AN (100 mM NaCl, 10 mM Tris-HCl, pH 7.4) containing 10 µM forward and reverse oligos. The solution was heated to 95 °C in a water bath for 3 min to denature secondary structure; the water bath was then switched off to allow gradual cooling over the course of 3 hours. The annealed dsDNA templates were stored at -20 °C.

### Peptide formation assay

The assay buffer (100 mM NaCl, 50 mM HEPES, 1 mM of each amino acid substrate, 0.1 mM $MgCl_2$, 10 µM $ZnCl_2$, 2% glycerol, 0.5 mM TCEP, 5 mM adenosine-5'-triphosphate [ATP]) was formulated to enable zinc-finger binding while minimizing zinc toxicity on NRPS enzyme activity (Figure S1). NaCl and HEPES were supplied from 20x DKP-M buffer (2 M NaCl, 1 M HEPES, pH 8.0). ATP disodium salt (Cat. No. 34369-07-8, Sigma) was freshly dissolved in $H_2O$ before use. The components were mixed thoroughly before adding enzymes and DNA templates. In the final reaction mixture, a pH of 7.5 ± 0.5 was determined with pH-Fix 2.0 - 9.0 test strips (Macherey-Nagel). Unless indicated otherwise, reactions were run at 37°C for 1 hour in a water bath.

### Chemical synthesis

To verify the identities of the nonribosomal peptide products, cyclo-(FP) was purchased from Bachem, fPVOL from Cambridge Research Biochemicals and further synthetic standards were synthesized in-house. Cyclic ornithine (*S*-3-aminopiperidin-2-one) is abbreviated as cOrn or O*.

#### *Analytics*

**NMR spectra** were recorded in deuterated solvents (Carl Roth, Germany) on a Bruker AVANCE II 300 or Bruker AVANCE III 600 MHz spectrometer, equipped with a Bruker Cryoplatform. The chemical shifts (δ) are reported in parts per million (ppm) relative to the solvent residual peak of DMSO-$d_6$ ($^{1}H$: 2.50 ppm, quintet; $^{13}C$: 39.5 ppm, heptet) or $CDCl_3$ ($^{1}H$: 7.24, singlet; $^{13}C$: 77.2, triplet). Some compounds show two sets of peaks due to conformational isomerism in which case only the major isomer was interpreted. **High resolution MS measurements** were performed on a Vanquish Horizon UHPLC system coupled with a Thermo QExactive HF-X Quadrupole-Orbitrap mass spectrometer (Thermo Scientific) equipped with a Acquity BEH C8 column (1.7 µm, 2.1 × 100 mm). Masses were detected by ESI as $M+H^+$ adducts in positive mode.

All reagents used were reagent grade and used as supplied (purchased from Sigma-Aldrich, Bachem, Fluorochem or Carl Roth). Reactions were performed at ambient temperature under argon atmosphere in anhydrous solvents (Acros Organics) unless otherwise stated. **Analytical thin-layer chromatography** was performed on silica 60 $F_{254}$ plates (0.25 mm, Merck). Compounds were visualized by dipping the plates in a ninhydrin/acetic acid solution followed by heating.

#### *HPLC purification of peptides*

Semi-preparative HPLC of peptide products was performed using a Phenomenex Luna C18(2) (5 µm, 250 x 10 mm) column connected to a Shimadzu Nexera LC-20AR system. Hydrochloride salts of peptides were dissolved in 5% MeCN and purified by semi-preparative HPLC using the following gradient: 0 - 5 min, 5% B; 5 - 25 min, 5% - 35% B; 25-30 min, 80% B; 30-38 min, 5% B (A: $H_2O$ with 0.1% (*v*/*v*) TFA, B: MeCN with 0.1% (*v*/*v*) TFA) with a flow rate of 8 mL/min. Fractions were collected from 5 min to 30 min. After purification, fractions containing the desired compounds ($H_2N$-D-Phe-L-Pro-L-cOrn: $R_t$ = 14.1 min, $H_2N$-D-Phe-L-Pro-L-Val-L-cOrn: $R_t$ = 17.4 min) were freeze dried, samples submitted to NMR measurements and pure compounds stored at -20 °C as TFA salts.

#### *Method A: Solution phase peptide coupling*

One equivalent of the carboxylic acid was dissolved in dry dichloromethane (DCM, 0.2 M solution) and cooled to 0 °C. *N*,*N*-Diisopropylethylamine (3 equivalents) was added drop-wise and the reaction was allowed to warm to room temperature. The acid was pre-activated through addition of hexafluorophosphate azabenzotriazole tetramethyl uronium (HATU, 1.2 equivalents) for 5 min at room temperature. Afterwards the amine (1.1 equivalents) was added and the reaction stirred at room temperature until completion as indicated by TLC (1 to 3 h). The reaction was quenched by pouring it onto 10% citric acid. Phases were separated and the aqueous phase extracted with DCM three times. The combined organic phases were washed with saturated $NaHCO_3$ solution and brine (one time each) and dried over $Na_2SO_4$. Volatiles were removed under reduced pressure. Purification by column chromatography (silica 60, DCM/MeOH 95:5) yielded the peptide product.

*Method B: Removal of Boc-protecting groups*

To the Boc-protected amine was added 4 M HCl in 1,4-dioxane (0.1 M final solution of the amine) at 0 °C (ice). After stirring for 5 min on ice the solution was allowed to warm to room temperature and stirred for further 3 h. Afterwards the volatile components were removed under vacuum, the residue dissolved in $H_2O$ and freeze dried to yield the respective amine hydrochloride salt.

*Synthesis of cyclo-L-ornithine (**2**)*

L-Orn (**1**, 3.0 g, 17.8 mmol) was dissolved in dry MeOH (40 mL) under argon atmosphere. Trimethylsilyl chloride (10 mL, 79.3 mmol) was added portion wise and the reaction stirred for 17 h at room temperature. Afterwards, the reaction was cooled to 0 °C (ice) and a 5 M solution of NaOMe in MeOH (27 mL, 135 mmol) added. After stirring at 0 °C for 10 min, the reaction was allowed to warm to room temperature and stirred for 2.5 h. Then, the reaction was acidified to pH 6 using concentrated HCl. The solvent was removed under reduced pressure and the residue re-dissolved in iPrOH (100 mL) and filtered. The filtrate was concentrated again under reduced pressure. Purification by column chromatography (silica 60, DCM/MeOH 7:3) yielded cyclo-L-ornithine **2** (1.74 g, 15.3 mmol, 86% yield) as a white-red foam.

**$^1$H NMR (600 MHz, DMSO):** δ = 8.43 (s, 3H), 8.04 (s, 1H), 3.69 (dd, J = 11.0, 6.1 Hz, 1H), 3.19 – 3.04 (m, 2H), 2.27 – 2.11 (m, 1H), 1.89 – 1.80 (m, 1H), 1.78 – 1.69 (m, 2H) ppm.
**$^{13}$C NMR (151 MHz, DMSO)**: δ = 167.3, 48.9, 40.8, 25.0, 20.2 ppm.
**UPLC-MS:** *m*/*z* = 114.98 $[M+H]^+$

*Synthesis of Boc-D-Phe-L-Pro-OMe (**5**)*

Method A with Boc-D-Phe-OH (**3**, 2.0 g, 7.7 mmol) as free acid and L-Pro-OMe (**4**, 1.1 g, 8.5 mmol) as amine yielded Boc-D-Phe-L-Pro-OMe **5** (2.2 g, 5.9 mmol, 77% yield) as an orange-white foam.

**$^1$H NMR (300 MHz, $CDCl_3$)** δ = 7.31 – 7.13 (m, 5H), 7.02 (d, *J* = 8.7 Hz, 1H), 4.43 (dd, *J* = 15.1, 8.5 Hz, 1H), 4.18 (dd, *J* = 8.4, 3.9 Hz, 1H), 3.58 (s, 3H), 3.28 – 3.12 (m, 1H), 2.92 – 2.67 (m, 3H), 2.06 – 1.93 (m, 1H), 1.90 – 1.59 (m, 3H) 1.32 (s, 9H).
**$^{13}$C NMR (75 MHz, DMSO):** δ = 172.2, 170.1, 154.9, 137.5, 129.3, 128.0, 126.3, 78.0, 58.6, 53.3, 51.6, 46.3, 37.3, 28.5, 28.1, 24.3 ppm.
**UPLC-MS:** *m*/*z* = 377.13 $[M+H]^+$

*Synthesis of Boc-D-Phe-L-Pro-OH (**6**)*

Compound **5** (530 mg, 1.4 mmol) was dissolved in THF (25 mL) and kept on ice (0 °C). 1 M LiOH (25 mL) was added, and the reaction stirred for 5 min at 0 °C. The reaction was allowed to warm to room temperature and stirred for further 2 h. THF was removed under vacuum, and the remaining aqueous solution acidified to pH 4 using 1 M HCl. The reaction mixture was extracted with DCM (20 mL) five times, the combined organic phases were washed with brine (100 mL) once, dried over $Na_2SO_4$ and concentrated under vacuum. Purification by column chromatography (silica 60, DCM/MeOH 95:5) yielded Boc-D-Phe-L-Pro-OH **6** (467 mg, 1.3 mmol, 93% yield) as a yellow-white solid.

**$^1$H NMR (300 MHz, DMSO)** δ = 12.54 (s, 1H), 7.35 – 7.10 (m, 5H), 6.95 (d, *J* = 8.8 Hz, 1H), 4.44 (dd, *J* = 15.0, 8.2, 1H), 4.11 (dd, *J* = 8.3, 3.5 Hz, 1H), 3.27 – 3.13 (m, 1H), 2.92 – 2.71 (m, 3H), 2.06 – 1.88 (m, 1H), 1.88 – 1.64 (m, 3H), 1.31 (s, 9H) ppm.
**$^{13}$C NMR (75 MHz, DMSO):** δ = 173.1, 169.8, 154.8, 137.5, 129.3, 128.0, 126.3, 77.9, 58.6, 53.3, 46.2, 37.5, 28.5, 28.1, 24.2 ppm.
**UPLC-MS:** *m*/*z* = 385.03 $[M+Na]^+$, *m*/*z* = 361.09 $[M-H]^-$

*Synthesis of $H_2N$-D-Phe-L-Pro- L-cOrn HCl (**8; fPO***)*

Method A with **6** (1.1 g, 2.9 mmol) as free acid and **2** (500 mg, 3.3 mmol) as amine yielded Boc-D-Phe-L-Pro-L-cOrn **7** (1.2 g, 2.6 mmol, 90% yield) as a yellow-white foam. Method B with **7** (1.2 g, 2.6 mmol) yielded $H_2N$-D-Phe-L-Pro-L-cOrn hydrochloride **8** (0.9 g, 2.7 mmol, quantitative yield) as a white solid.

**$^1$H NMR (600 MHZ, DMSO):** δ = 8.28 (s, 2H), 8.05 (d, *J* = 8.3 Hz, 1H), 7.57 (s, 1H), 7.37 – 7.20 (m, 5H), 4.36 (s, 1H), 4.21 (dd, *J* = 7.8, 2.8 Hz, 1H), 4.08 (ddd, *J* = 10.4, 8,1, 6.4 Hz, 1H), 3.58 – 3.46 (m, 1H), 3.14 – 3.09 (m, 2H), 3.07 (dd, *J* = 13.4, 6.1 Hz, 1H), 2.96 (dd, *J* = 13.3, 8.4 Hz, 1H), 2.68 (dd, *J* = 16.8, 7.1 Hz, 1H), 1.94 – 1.86 (m, 2H), 1.84 – 1.68 (m, 4H), 1.67 – 1.56 (m, 1H), 1.49 – 1.41 (m, 1H) ppm.
**$^{13}$C NMR (151 MHz, DMSO):** δ = 170.6, 169.6, 166.4, 134.5, 129.5, 128.5, 127.4, 59.7, 51.9, 48.8, 46.6, 41.0, 36.7, 29.2, 27.6, 23.7, 21.0 ppm.
**UPLC-MS:** *m*/*z* = 359.27 $[M+H]^+$

**HRMS:** calculated for $C_{19}H_{27}N_4O_3$ *m/z* = 359.2078 $[M+H]^+$; measured *m/z* = 359.2076 $[M+H]^+$
**HPLC:** $R_t$ = 14.1 min

*Synthesis of Boc-L-Val-L-cOrn (**10**)*

Method A with Boc-L-Val-OH (**9**, 1.7 g, 7.7 mmol) as free acid and **2** (0.8 g, 7 mmol) as amine yielded Boc-L-Val-L-cOrn **10** (1.4 g, 4.6 mmol, 66% yield) as a white solid.

**$^1$H NMR (300 MHz, CDCl3)**: δ = 7.28 (d, *J* = 4.3 Hz, 1H), 6.67 (s, 1H), 5.33 (d, *J* = 8.6 Hz, 1H), 4.34 – 4.18 (m, 1H), 4.10 – 3.96 (m, 1H), 3.40 – 3.23 (m, 2H), 2.58 – 2.39 (m, 1H), 2.22 – 2.01 (m, 1H), 1.96 – 1.82 (m, 2H), 1.71 – 1.50 (m, 1H), 1.42 (s, 9H), 0.95 (d, *J* = 6.8 Hz, 3H), 0.90 (d, *J* = 6.8 Hz, 3H) ppm.
**$^{13}$C NMR (75 MHz, CDCl3)**: δ = 171.9, 171.6, 155.8, 79.5, 59.6, 50.3, 41.6, 31.3, 28.3, 27.1, 20.9, 19.2, 17.6 ppm.
**UPLC-MS:** *m/z* = 314.14 $[M+H]^+$

*Synthesis of $H_2N$-L-Val-L-cOrn HCl (**11**)*

Method B with **10** (500 mg, 1.6 mmol) yielded $H_2N$-L-Val-L-cOrn hydrochloride **11** (400 mg, 1.6 mmol, quantitative yield).

**$^1$H NMR (300 MHz, DMSO)**: δ = 8.63 (d, *J* = 8.4 Hz, 1H), 8.27 (d, *J* = 3.5 Hz, 3H), 7.63 (s, 1H), 4.35 – 4.10 (m, 1H), 3.64 – 3.57 (m, 1H), 3.22 – 3.00 (m, 2H), 2.17 – 2.03 (m, 1H), 2.02 – 1.91 (m, 1H), 1.87 – 1.56 (m, 3H), 0.99 (d, *J* = 3.6 Hz, 3H), 0.96 (d, *J* = 3.6 Hz, 3H) ppm.
**$^{13}$C NMR (75 MHz, DMSO):** δ = 169.1, 167.4, 66.4, 57.3, 48.9, 41.0, 29.8, 27.5, 20.9, 18.2, 18.0 ppm.
**UPLC-MS:** *m/z* = 214.03 $[M+H]^+$

*Synthesis of $H_2N$-D-Phe-L-Pro-L-Val-L-cOrn HCl (**13; fPVO***)*

Method A with **6** as free acid and **11** (0.8 g, 7 mmol) as amine yielded Boc-D-Phe-L-Pro-L-Val-cOrn **12** (*m/z* = 558.38 $[M+H]^+$). Method B with **12** (176 mg, 0.3 mmol) yielded $H_2N$-D-Phe-L-Pro-L-Val-L-cOrn hydrochloride **13** (67.4 mg, 0.15 mmol, 50% yield) as a white solid.

**$^1$H NMR (600 MHz, DMSO)**: δ = 8.26 (s, 3H), 7.93 (d, *J* = 9.0 Hz, 1H), 7.87 (d, *J* = 7.6 Hz, 1H), 7.58 (s, 1H), 7.37 – 7.28 (m, 3H), 7.25 – 7.18 (m, 2H), 4.44 – 4.29 (m, 2H), 4.17 – 4.04 (m, 2H), 3.58 – 3.47 (m, 1H), 3.15 – 3.09 (m, 2H), 3.06 (dd, *J* = 13.3, 6.0 Hz, 1H), 2.95 (dd, *J* = 13.3, 8.3 Hz, 1H), 2.76 – 2.64 (m, 1H), 2.05 – 1.94 (m, 2H), 1.81 – 1.69 (m, 4H), 1.60 – 1.48 (m, 1H), 1.47 – 1.39 (m, 1H), 0.89 (d, *J* = 6.8 Hz, 3H), 0.87 (d, *J* = 6.8 Hz, 3H) ppm.
**$^{13}$C NMR (151 MHz, DMSO)**: δ = 170.9, 170.4, 169.6, 166.3, 134.5, 129.5, 128.5, 127.4, 59.4, 57.8, 51.9, 49.0, 46.7, 40.8, 36.8, 30.6, 29.3, 27.4, 23.8, 20.9, 19.3, 18.1 ppm.
**UPLC-MS:** *m/z* = 458.43 $[M+H]^+$
**HRMS:** calculated for $C_{24}H_{36}N_5O_4$ $[M+H]^+$ *m/z* = 458.2762 $[M+H]^+$; measured *m/z* = 458.2762 $[M+H]^+$
**HPLC:** $R_t$ = 17.4 min

**Peptide quantification by UPLC-MS/MS**

For quantification of peptides, a Xevo TQ-S micro tandem mass spectrometer (Waters) was used. Samples were diluted 20-fold with 50% ethanol containing 0.1% formic acid (FA) when analyzing cyclo-(fP), fPO*, and fPVO*. A 2 µL aliquot was injected onto a CORTECS UPLC C18 column (1.6 µm, 2.1 x 50 mm; Waters) equipped with a VanGuard pre-column (2.1 x 5 mm; Waters) connected to an H-class UPLC system (Waters). Elution was performed at 40 °C with acetonitrile (MeCN) in $H_2O$ containing 0.1% FA at a flow rate of 0.5 mL/min (2% MeCN at 0.00 - 0.30 min, linear gradient from 2 - 50% MeCN at 0.30 - 1.50 min, 95% MeCN at 1.51 - 1.80 min, and 2% MeCN at 1.81 - 2.80 min). Ionization was performed in an ESI Z-Spray probe operated in positive mode at a capillary voltage of 0.5 kV and a desolvation temperature of 600 °C. The cone voltage was set to 20 V and multiple reaction monitoring (MRM) was performed using argon as collision gas (Table S5).

#### Fluorescence polarization assay

Affinity between zinc finger domains and DNA was measured in a fluorescence polarization assay. The DNA probes (Table S4) were designed according to a previous report.[8] In short, a fluorescein-dT base was incorporated two bases upstream of the zinc finger binding site on the reverse oligo. Synthetic oligos (Eurofins) were dissolved in buffer NE (5 mM Tris-HCl, pH 8.5) at 100 µM concentration for long term storage. Double stranded DNA probes were generated by annealing of 10 µM forward and reverse oligos in buffer AN (100 mM NaCl, 10 mM Tris-HCl, pH 7.4). The DNA solutions were heated to 80 °C for 4 min to resolve secondary structure followed by gradual cooling to room temperature for 30 min. The resulting 10 µM probes were stored in the dark at -20 °C.

The fluorescence polarization assay buffer (100 mM NaCl, 50 mM HEPES, 0.1 mM $MgCl_2$, 9 µM $ZnCl_2$, 5% glycerol, 0.5 mM TCEP, 100 µg/mL bovine serum albumin [BSA; Roth Cat.No. 90604-29-8]) resembled the buffer for peptide formation as closely as possible. The buffer was freshly prepared, filtered through a cellulose acetate (CA) filter with a pore size of 0.22 µm (TH Geyer), and kept at room temperature for all following dilution procedures. DNA probes were diluted to 100 nM. Proteins were first diluted to 2 µM, centrifuged at 16,000 x g and 16 °C for 8 min to remove potential precipitates, and then subjected to 2-fold serial dilutions. In black 96-well plates (BRANDplates, Cat.No. 781668, BrandTech), 90 µL protein dilution were added to 10 µL of 100 nM DNA probe. After thoroughly mixing by pipetting, the sample was kept at room temperature for 10 min and measured on a Synergy H1 fluorescence microplate reader (Biotek) using a polarization filter set (485/528 nm) at 24.5 ± 1.0 °C. The polarization output (mP; Equation 1) was plotted against protein concentration (nM) and fitted to a bimolecular binding model (Equation 2) with R version 3.4.3 (Table S6).

Equation 1:

$$\text{polarization [mp]} = \frac{\text{Intensity}_{\parallel} - \text{Intensity}_{\perp}}{\text{Intensity}_{\parallel} + \text{Intensity}_{\perp}} \times 1000$$

where the incident light is subjected to a linear polarizer and
$\text{Intensity}_{\parallel}$: the intensity of emitted light measured using a parallel polarizing filter
$\text{Intensity}_{\perp}$: the intensity of emitted light measured using a perpendicular polarizing filter

Equation 2:

$$\mathrm{F} = \mathrm{F}_0 + \frac{\mathrm{F}_{\max} - \mathrm{F}_0}{\mathrm{A}_0} \times \frac{(\mathrm{A}_0 + \mathrm{P}_0 + K_\mathrm{d}) - \sqrt{(\mathrm{A}_0 + \mathrm{P}_0 + K_\mathrm{d})^2 - 4\mathrm{A}_0\mathrm{P}_0}}{2}$$

where these parameters are measured or controlled:
$\mathrm{F}$: polarization
$\mathrm{F}_0$: polarization in the absence of protein
$\mathrm{A}_0$: DNA probe concentration
$\mathrm{P}_0$: protein concentration
and the following parameters are predicted by the model:
$\mathrm{F}_{\max}$: polarization at saturating protein concentration
$K_\mathrm{d}$: dissociation constant

#### Isothermal titration calorimetry (ITC)

ITC measurements (Figure S4) were performed at 25 °C in 50 mM HEPES buffer pH 8, 150 mM NaCl and 5% (v/v) glycerol using a MicroCal PEAQ-ITC calorimeter (Malvern Instruments). A solution containing 50 µM DnGrsB3 was titrated with 1 mM of the respective Dc peptides. Peptides InxA-Dc8 (QALLKGDI), InxA-Dc8-GS (QALLKGDIGS) and InxA-Dc6-GS (QALLKGGS) were custom synthesized on a 20-24 mg scale at >95% HPLC purity by GenScript Biotech (Netherlands) as HCl salts. ITC measurements started with an initial delay of 60 s. The first injection of 0.4 µl was followed by 24 injections with 3 µl each for titration with peptides InxA-Dc8 and InxA-Dc8-GS or 12 injections with 3 µl each for titration with peptide InxA-Dc6-GS. Injections were performed in intervals of 150 s. The reference power was set to 10 µcal/s and the stir speed to 750 rpm. For each measurement, the “high feedback” mode was selected. The thermograms were analyzed using the MicroCal PEAQ-ITC Analysis Software v1.22 assuming a one site binding model. Signals in thermograms resulting from titration of the respective peptides into buffer were subtracted point-by-point as background correction. *c*-Values were calculated using equation 3:

Equation 3:

$$c = nK_a[M]_T$$

where $K_a$ is the binding constant, $[M]_T$ the total macromolecule concentration in the cell and $n$ the binding stoichiometry. With InxA-Dc8 and InxA-Dc8-GS, binding was detected but the c-value <1 indicated unreliable quantification of the binding constants. In both cases the apparent interaction was only qualitatively noted as weak binding. With InxA-Dc6-GS, no binding was observed.

#### Analytical size exclusion chromatography

To physically confirm module assembly on DNA template, 2.5 µM each of TycB1-N, GrsB2-P, and GrsB3-Z were added to 2 µM of SEC-DNA in SEC sample buffer (100 mM NaCl, 50 mM HEPES pH 8.0, 35 µM $ZnCl_2$, 2% glycerol, and 0.5 mM TCEP). The solution was centrifuged at 16,000 x g and 16 °C for 6 min and equilibrated at 10 °C for 20 min. A 20 µL aliquot was injected onto a Superdex 200 Increase 3.2/300 analytical size exclusion column (GE Healthcare) connected to an NGC Quest 10 Plus Chromatography System (Bio-Rad) while bypassing the column selection valve. Elution was performed at 10 °C in 2.4 mL buffer MT (100 mM NaCl, 50 mM HEPES pH 8.0) at a flow rate of 0.075 mL/min. Absorbance at 260 and 280 nm was recorded in a 5 mm flow cell.

Molecular weight standards were prepared by dissolving components of Gel Filtration Calibration Kit (GE Healthcare) separately in buffer T (150 mM NaCl, 50 mM Tris pH 7.5) at the following concentrations: 20 mg/mL thyroglobin (T), 2 mg/mL ferritin (F), 20 mg/mL aldolase (A), and 20 mg/mL conalbumin (C). A mixture containing equal volumes of each protein was stored at -80 °C. Prior to analysis, the mixture was thawed, diluted 2.5-fold with buffer MT, and centrifuged at 16,000 x g and 16 °C for 6 min. A 20 µL aliquot was analyzed via size exclusion as described above.

#### Homology modelling

A homology model of a two-modular DT-NRPS was built based on the sequence of GrsB3-Z and a DNA template with two corresponding binding sites and a 20 bp spacer. The DNA was modelled as ideal B-form DNA with the script "build_DNA.mcr" in YASARA (version 19.12.14).[9] Two models of Zif268 prepared on the Swiss-Model[10] server were aligned with the DNA in PyMOL.[11] After energy minimization in Yasara with default settings, a homology model of the Dn/Dc domain (Swiss-Model) was changed to the sequence from GrsB3-Z at the termini and appended to one of the ZF. Next, the homology model of GrsB3-CAT (Swiss-Model) was added. The linkers between the homology models were modelled in an arbitrary low energy conformation not based on experimental data that was not further optimized. The Dn-CAT-Dc-ZF2 was duplicated and overlayed on the ZF bound to the other binding site. Since there was a slight clash between both CAT modules, a dihedral in the Dn-CAT linker was adjusted in both modules until the clash disappeared.

#### Sinusoidal fit of spacer length dependence

The spacer length dependence observed with the trimodular DT-NRPS (Figure 4) was nonlinearly fit to a cosine curve in R version 3.6.3 using the following code:

```
sta <- list(a=40, b=10, c=100, d=-6)
m <- nls(y~a*cos(x*2*pi/b+d) + c, start = sta)
summary(m)
```

The optimized parameters of fit are a = 43 ± 6 nM (amplitude), b = 10.1 ± 0.3 bp (wavelength), c = 90 ± 4 nM (y-axis offset), and d = -6.6 ± 0.4 bp (x-axis offset) with a residual standard error of 35.5 nM for 70 degrees of freedom.

### Supplementary Figures and Tables

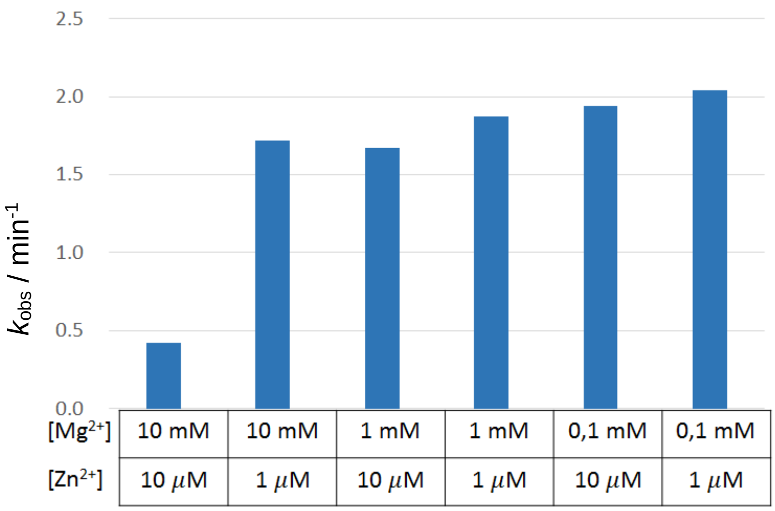

**Figure S1.**
Optimization of $Zn^{2+}$ and $Mg^{2+}$ concentration. Reactions were performed with 0.5 µM GrsA and TycB1 in buffer containing 100 mM NaCl, 50 mM HEPES pH 8.0, 2 % glycerol, and the indicated metal ions. Cyclo-(fP) was quantified by LC-MS/MS after 1 h at 37°C and divided by the enzyme concentration to obtain $k_{obs}$.

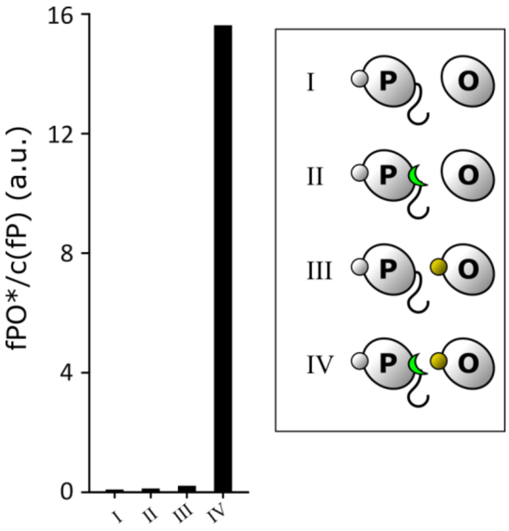

**Figure S2. Portability of the docking domain.**
GrsA was added to all reactions. In addition, proteins TycB1-linker o-P (I & III), TycB1-linker Dc-P (II & IV), GrsB3 (I & II), and Dn-GrsB3 (III & IV) were supplied (Table S1 and S3). The data were collected from a single batch of protein purified through nickel affinity chromatography. Reaction conditions: 1 µM of each protein, no DNA template or $ZnCl_2$; incubation at 37°C for 3 h. The ratio of fPO* to cyclo-(fP) determined by tandem UPLC-MS is a robust parameter to assess docking domain efficiency since the ratio is less sensitive to variations in enzyme purity than the overall yield.

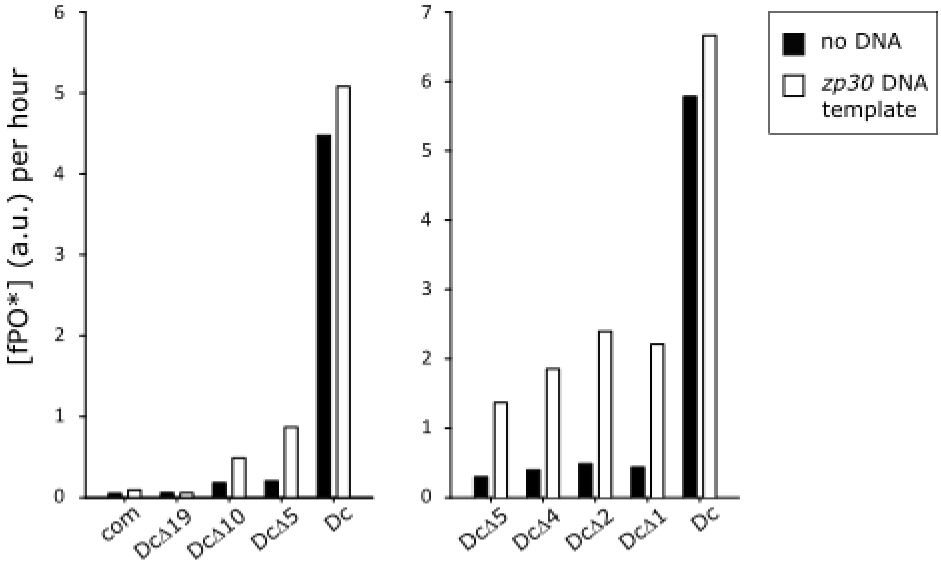

**Figure S3. Docking domain truncation tests.**
Reactions contained GrsA, TycB1-“X”-P, and Dn-GrsB3-linker com-Z. The linker “X” is specified on the x axes. Module assembly was tested on a DNA template with 30-bp spacer. Proteins were purified by nickel affinity chromatography only and the data was collected using a single batch of protein. Reaction conditions: 1 µM of each protein, and 0.5 µM DNA template zp30; incubation at 37°C for 2 h (left panel) or 1 h (right panel).

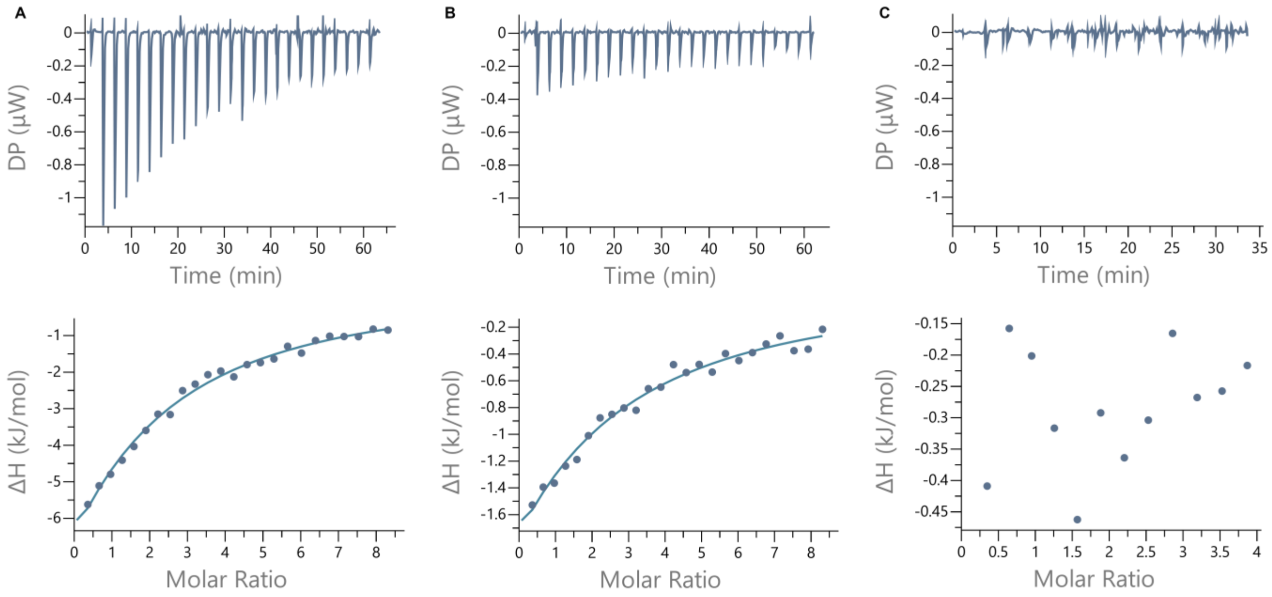

**Figure S4. Docking domain binding affinity.**
ITC thermograms (top) and derived binding curves (bottom) for titration of module Dn-GrsB3 with different variants of the InxA-Dc fragment responsible for interaction.[12] DP: power differential between reference and sample cell. (A) Titration with full length InxA-Dc8 ($K_D$: 280 ± 40 µM, ∆H: -41 ± 6 kJ/mol, c: 0.177). (B) Titration with peptide InxA-Dc8-GS containing additional C-terminal amino acids GS to mimic the linker region in ZF containing constructs ($K_D$: 350 ± 80 µM, ∆H: -13 ± 3 kJ/mol, c: 0.141). (C) Titration with peptide InxA-Dc6-GS truncated by two amino acids.

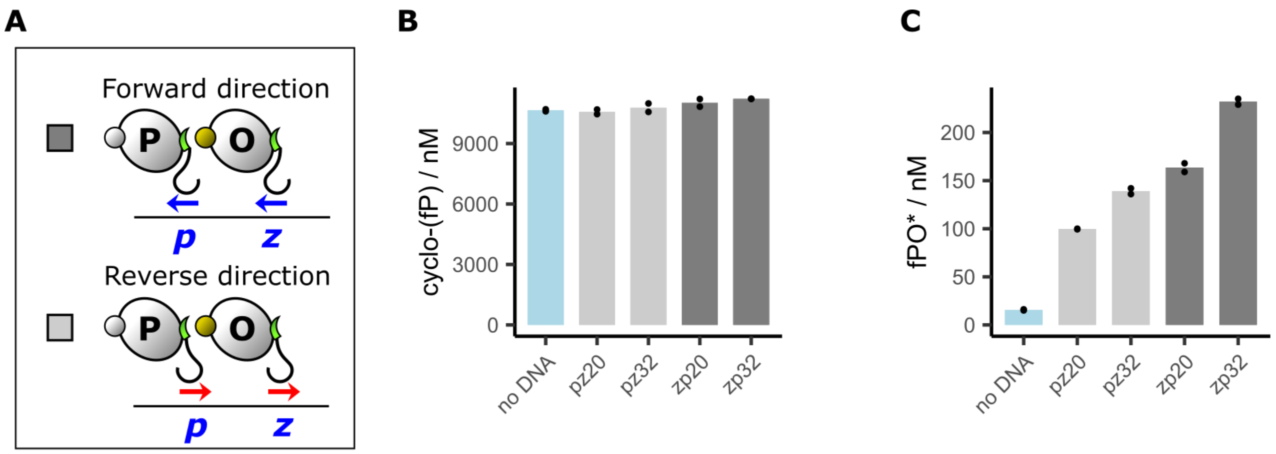

**Figure S5. Influence of binding site orientation on peptide formation efficiency.**
(A) Scheme of the ZF orientation. (B) DNA-independent formation of the control cyclo-(fP). (C) DNA-dependent formation of fPO* on templates with different ZF binding site orientations and linker lengths. Reaction conditions: 0.4 µM GrsA, 0.1 µM TycB1-P, 0.5 µM GrsB3-Z, and 0.4 µM DNA template. Data points are shown for a biological duplicate.

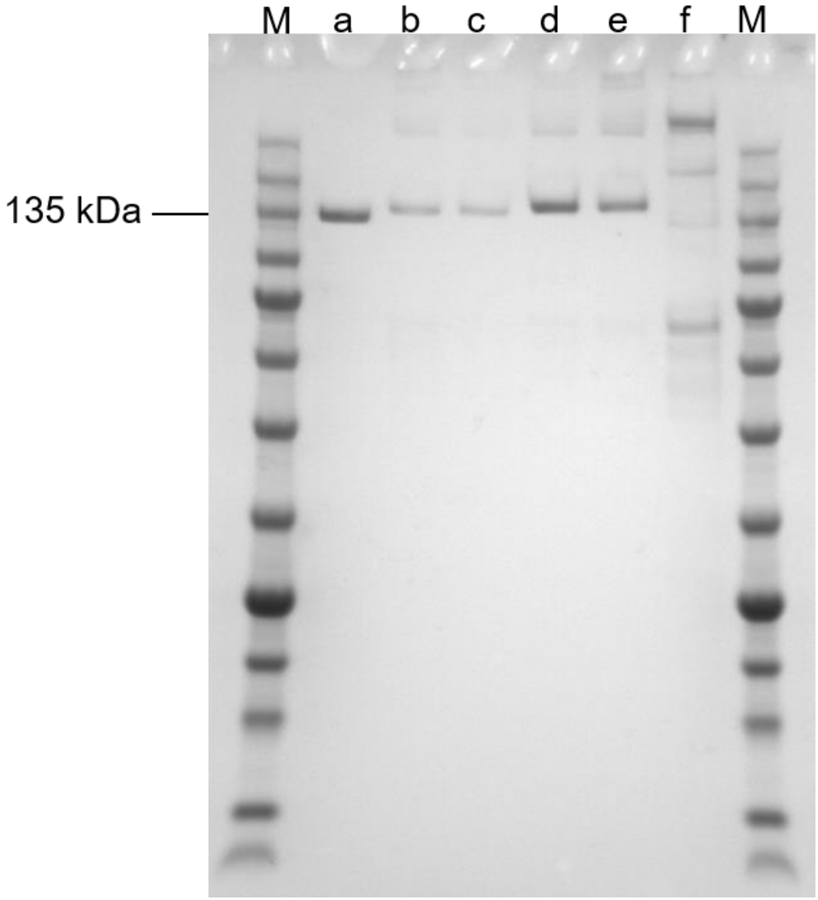

**Figure S6. SDS PAGE of purified proteins.**
Electrophoretic separation was performed on Bolt 4-12% Bis-Tris Plus Gels (Thermo Fisher). Per well, 500 ng protein were applied. M: Marker (SERVA Triple Color Protein Standard III, 5 µl used), a: GrsA (size 128 kDa), b: TycB1-P (size 131 kDa), c: TycB1-N (size 131 kDa), d: GrsB2-P (size 139 kDa), e: GrsB3-Z (size 141 kDa), f: GrsB123 (size 358 kDa). The gel was run at 200 V for 22 min in MES SDS buffer and stained with Quick Coomassie stain (Serva).

**Table S1. Plasmids and proteins used in this study.**

| Plasmid name | Genetic layout* | Encoded protein | Usage | Source |
|---|---|---|---|---|
| **pSU18-grsA** | pSU18 vector; Plac-grsA; $Chl^R$ | GrsA | Source of NRPS modules; Source of pSU18 vector; GrsA expression | [3] |
| **pTrc99a-tycB1** | Ptrc-tycB1; $Amp^R$ | not used | Source of NRPS genes | [1] |
| **pTrc99a_grsB_M3574L** | Ptrc-grsB; $Amp^R$ | not used | Source of NRPS genes | [3] |
| **pSU18F-tycB1-P (o)** | Plac-tycB1-linker o-pbsII; N' His6 tag | TycB1-linker o-P | Verify Dn & Dc portability | This study |
| **pSU18F-tycB1-P (Dc)** | Plac-tycB1-linker Dc-pbsII; N' His6 tag | TycB1-linker Dc-P | Verify Dn & Dc portability | This study |
| **pSU18-grsB3** | Plac-grsB3 | GrsB3 | Verify Dn & Dc portability | This study |
| **pSU18-Dn-grsB3** | Plac-Dn-grsB3 | Dn-GrsB3 | Verify Dn & Dc portability | This study |
| **pSU18-tycB1-P (com)** | Plac-tycB1-linker com-pbsII | TycB1-linker com-P | Dc truncation test | This study |
| **pSU18-tycB1-P (Dc)** | Plac-tycB1-linker Dc-pbsII | TycB1-linker Dc-P | Dc truncation test | This study |
| **pSU18-tycB1-P (DcΔ1)** | Plac-tycB1-linker DcΔ1-pbsII | TycB1-linker DcΔ1-P | Dc truncation test | This study |
| **pSU18-tycB1-P (DcΔ2)** | Plac-tycB1-linker DcΔ2-pbsII | TycB1-linker DcΔ2-P (TycB1-P) | Dc truncation test; DNA spacer length test | This study |
| **pSU18-tycB1-P (DcΔ4)** | Plac-tycB1-linker DcΔ4-pbsII | TycB1-linker DcΔ4-P | Dc truncation test | This study |
| **pSU18-tycB1-P (DcΔ5)** | Plac-tycB1-linker DcΔ5-pbsII | TycB1-linker DcΔ5-P | Dc truncation test | This study |
| **pSU18-tycB1-P (DcΔ10)** | Plac-tycB1-linker DcΔ10-pbsII | TycB1-linker DcΔ10-P | Dc truncation test | This study |
| **pSU18-tycB1-P (DcΔ19)** | Plac-tycB1-linker DcΔ19-pbsII | TycB1-linker DcΔ19-P | Dc truncation test | This study |
| **pSU18-Dn-grsB3-Z (com)** | Plac-Dn-grsB3-linker com-zif268 | Dn-GrsB3-linker com-Z | Dc truncation test | This study |
| **pSU18-grsB123** | Plac-grsB123 | GrsB123 | Positive control (fused multimodule) | This study |
| **pSU18-tycB1-N (DcΔ2)** | Plac-tycB1-linker DcΔ2-nre | TycB1-N | Multimodule assembly | This study |
| **pSU18-Dn-grsB2-P (DcΔ2)** | Plac-Dn-grsB2-linker DcΔ2-pbsII | GrsB2-P | Multimodule assembly | This study |
| **pSU18-Dn-grsB3-Z (DcΔ2)** | Plac-Dn-grsB3-linker DcΔ2-zif268 | GrsB3-Z | DNA spacer length test; Multimodule assembly | This study |
| **pSU18-Z-Dn-grsB3** | Plac-zif268-Dn-grsB3 | Z-GrsB3 | DNA spacer length test | This study |
| **pSU18-B-Dn-grsB4** | Plac-zfb-Dn-grsB4 | B-GrsB4 | Multimodule assembly | This study |

*For protein sequences encoded by the linkers, see Table S3. Genes of NRPS modules are highlighted in yellow, genes of linkers in green and genes of zinc fingers in cyan.

**Table S2. Coding sequences of recombinant NRPS proteins used in multimodule assembly.**
Sequences along with annotations are shown in GenBank file format, each starting with “LOCUS” and ending with “//”. The text can be saved with “.gb” file extension using text editors and visualized by molecular biology software.

```
LOCUS       tycB1-N          3522 bp ds-DNA     linear       06-FEB-2019
DEFINITION  .
FEATURES             Location/Qualifiers
     CDS             1..3522
                     /label="tycB1-N"
     misc_feature    1..3126
                     /label="NRPS module tycB1"
     misc_feature    3127..3219
                     /label="linker Dc_2"
     misc_feature    3127..3204
                     /label="docking domain Dc_2"
     misc_feature    3220..3480
                     /label="zinc finger domain nre"
ORIGIN
        1 ATGGGTGTAT TTAGCAAAGA ACAAGTTCAG GATATGTATG CGTTGACCCC GATGCAAGAG
       61 GGGATGCTGT TTCACGCCTT GCTCGACCAA GAGCACAACT CGCATCTGGT ACAGATGTCG
      121 ATTTCGTTGC AGGGCGATCT TGACGTTGGG CTATTTACGG ATAGCCTGCA TGTGCTGGTA
      181 GAGAGATACG ATGTATTCCG CACGTTGTTT CTCTATGAAA AGCTGAAGCA GCCTTTGCAA
      241 GTTGTCTTGA AGCAACGGCC TATTCCGATC GAATTTTACG GCTTGTCTGC CTGCGACGAG
      301 TCCGAGAAAC AACTTCGCTA TACGCAATAC AAAAGGGCGG ATCAGGAGCG GACGTTTCAT
      361 CTGGCAAAAG ACCCGTTGAT GCGGGTCGCC CTTTTCCAAA TGTCCCAGCA CGACTACCAG
      421 GTCATCTGGA GCTTTCATCA CATCCTCATG GACGGCTGGT GCTTCAGCAT TATTTTTGAT
      481 GACTTGCTTG CCATCTACTT GTCCTTGCAA AACAAGACGG CACTCTCCCT GGAGCCCGTA
      541 CAGCCATACA GTCGCTTTAT CAACTGGCTG GAAAAACAAA ATAAACAGGC CGCTCTCAAC
      601 TATTGGAGCG ACTATCTGGA AGCCTATGAA CAAAAGACTA CCTTGCCGAA GAAGGAAGCT
      661 GCCTTCGCCA AAGCATTTCA ACCAACCCAA TACCGCTTTT CGCTGAACCG CACCTTGACC
      721 AAGCAGCTCG GGACCATCGC CAGTCAAAAT CAAGTGACGC TATCGACGGT GATTCAAACG
      781 ATCTGGGGAG TTCTCCTGCA AAAATACAAT GCGGCCCATG ATGTGCTGTT CGGCTCTGTT
      841 GTATCCGGAC GCCCTACAGA CATCGTCGGA ATCGACAAAA TGGTTGGCTT GTTTATCAAT
      901 ACGATTCCAT TCCGGGTGCA AGCGAAAGCT GGTCAAACGT TTTCCGAGCT GTTGCAAGCT
      961 GTGCACAAAA GAACTTTGCA ATCACAGCCG TATGAGCACG TGCCTTTGTA CGACATTCAA
     1021 ACTCAGTCCG TCTTGAAGCA GGAGCTGATT GACCACCTGC TGGTCATCGA AAATTACCCG
     1081 CTGGTAGAGG CTTTGCAGAA AAAAGCATTG AACCAGCAGA TCGGCTTCAC GATTACTGCT
     1141 GTGGAAATGT TCGAGCCGAC CAATTACGAC TTGACTGTCA TGGTGATGCC AAAAGAAGAG
     1201 CTTGCCTTCC GTTTTGACTA CAATGCGGCT CTGTTTGACG AACAGGTCGT GCAAAAACTG
     1261 GCGGGGCACC TCCAACAGAT CGCGGATTGC GTGGCAAACA ATTCGGGAGT CGAGCTTTGC
     1321 CAGATTCCGT TGCTGACAGA AGCAGAAACT AGCCAGCTGT TGGCAAAGCG TACGGAAACA
     1381 GCGGCTGACT ATCCTGCCGC AACCATGCAC GAGCTGTTTT CGCGGCAGGC AGAAAAAACG
     1441 CCTGAGCAAG TGGCGGTAGT CTTCGCGGAT CAGCACCTGA CGTATCGGGA GCTGGATGAA
     1501 AAATCCAATC AGCTCGCCCG CTTTTTGCGC AAAAAAGGCA TTGGCACGGG CAGTCTTGTC
     1561 GGCACGCTGC TGGATCGCTC GCTGGACATG ATCGTCGGAA TCCTCGGCGT CTTGAAGGCA
     1621 GGCGGCGCAT TTGTGCCGAT CGACCCGGAG TTGCCTGCCG AACGAATCGC TTACATGCTG
     1681 ACGCATAGCA GAGTTCCATT GGTCGTGACG CAAAATCATT TGCGGGCAAA AGTGACCACG
     1741 CCTACAGAAA CAATTGACAT CAACACAGCG GTGATCGGGG AAGAGAGCCG CGCCCCTATC
     1801 GAATCGCTCA ATCAGCCGCA TGACTTGTTT TACATCATCT ATACGTCCGG AACGACAGGG
     1861 CAACCGAAAG GCGTCATGCT GGAGCATCGC AACATGGCGA ACCTGATGCG TTTTACGTTT
     1921 GATCAGACGA ACATCGCTTT TCATGAAAAA GTGTTGCAGT ATACCACGTG CAGCTTTGAT
     1981 GTTTGCTACC AGGAAATTTT CTCCACGCTG CTATCCGGGG GCCAGCTCTA CCTGATCACG
     2041 AACGAGCTGA GACGGCATGT GGAAAAGCTG TTTGCTTTCA TCCAGGAAAA GCAGATCAGC
     2101 ATTTTGTCTC TCCCGGTGTC CTTCCTGAAA TTTATTTTTA ACGAACAAGA CTACGCGCAA
     2161 AGCTTCCCGC GTTGTGTCAA ACATATCATC ACGGCCGGGG AACAACTCGT CGTCACACAC
     2221 GAGCTGCAAA AGTATCTGCG CCAGCATCGC GTATTTTTGC ACAATCACTA CGGCCCGTCG
     2281 GAGACGCATG TGGTGACGAC ATGCACGATG GACCCGGGAC AGGCGATACC AGAGCTGCCG
     2341 CCCATCGGAA AGCCGATCAG CAACACAGGC ATTTACATTT TGGATGAAGG GCTGCAATTG
     2401 AAGCCGGAGG GGATCGTCGG GGAGTTGTAC ATTTCCGGCG CAAACGTAGG AAGAGGGTAT
```

```
2461 TTGCACCAGC CGGAGCTGAC CGCGGAGAAG TTTCTCGACA ATCCGTATCA GCCAGGCGAA
2521 AGAATGTACC GAACGGGTGA TCTGGCGCGT TGGTTGCCGG ATGGCCAGCT CGAATTTTTG
2581 GGCCGAATCG ACCATCAGGT AAAAATCAGG GGCCATCGCA TCGAGCTGGG AGAGATCGAA
2641 TCGCGCCTGC TCAACCATCC CGCCATCAAG GAAGCGGTGG TTATCGACCG AGCAGACGAG
2701 ACAGGCGGCA AGTTTTTGTG CGCCTATGTC GTCCTGCAAA AAGCGCTCAG CGACGAAGAG
2761 ATGCGGGCAT ACTTGGCGCA AGCGTTGCCG GAGTATATGA TCCCTTCCTT TTTCGTGACG
2821 CTGGAGCGGA TTCCAGTCAC GCCGAACGGA AAAACAGACA GGCGAGCTTT GCCGAAGCCG
2881 GAAGGAAGTG CCAAGACGAA AGCGGATTAC GTCGCCCCGA CGACTGAGCT GGAACAAAAG
2941 CTGGTCGCGA TTTGGGAGCA AATTCTTGGC GTGTCGCCGA TCGGCATTCA GGATCATTTT
3001 TTCACGCTGG GCGGCCATTC GTTAAAAGCG ATTCAGCTCA TTTCCCGCAT CCAAAAGGAA
3061 TGCCAGGCGG ATGTCCCGCT GCGCGTCCTG TTTGAGCAAC CGACGATTCA AGCGCTGGCA
3121 GCGTATGTAA CAGTACATCT TCTTAAGGAG AAACGTAAGC ATTTCCAAGC GGGTCAAGAA
3181 ACTGCACAAG CTCTGTTAAA AGGTGGATCC GCGGCAGCCC CTGGTGAAAA GCCTTATGCC
3241 TGCCCAGTGG AAAGCTGTGA CCGCCGTTTC TCTCAGAGCC ACGATTTGAC GAAACACATC
3301 CGCATCCACA CGGGACAGAA GCCCTTCCAA TGTCGTATTT GTATGCGCAA TTTCAGCGAT
3361 AGCAGCAAAC TGAGCCGCCA TATCCGCACG CACACCGGGG AAAAACCCTT TGCTTGTGAC
3421 ATCTGTGGTC GTAAATTTGC CCGCTTGGAT AATCGTACCG CTCACACCAA AATTCACACA
3481 ACCTGCTTTT GCTCGCTTAA ACATCACCAT CACCATCACT AA
//
```

```
LOCUS       grsB2-P       3672 bp ds-DNA     linear       06-FEB-2019
DEFINITION  .
FEATURES             Location/Qualifiers
     misc_feature    3376..3630
                     /label="zinc finger domain pbsII"
     misc_feature    3283..3375
                     /label="linker Dc_2"
     CDS             1..3672
                     /label="grsB2-P"
     misc_feature    232..3282
                     /label="NRPS module grsB2"
     misc_feature    1..231
                     /label="docking domain Dn"
     misc_feature    3283..3360
                     /label="docking domain Dc_2"
ORIGIN
        1 ATGAAAGATG CAGCACAAAT TGTCAATGAG GCTTTAGATC AGGGAATAAC ACTGTTCGTG
       61 GCTGAGGATC GTTTGCAATA CGAAACCCGC CTCAGCAATA TTCCGGCAGA CTTAATCAGT
      121 GAATGGAAAC AGCATAAACA AGAGTTGATT GATTTTCTGA ACCAGCTCGA TTCATCAGAA
      181 GAGCAGGTGA GTACCCATCA ACTACAGGGT ATTCCACGTT ATGAACGGGC TGAGTACTAT
      241 CCTGTATCAT CAGTTCAAAA AAGAATGTTT ATTCTTAATG AATTTGATCG TTCAGGTACG
      301 GCCTATAATT TACCTGGTGT TATGTTTCTA GATGGAAAAT TGAACTACCG ACAATTGGAA
      361 GCAGCGGTAA AAAAATTAGT TGAGCGACAT GAAGCGCTGC GTACTTCCTT TCATTCAATT
      421 AATGGGGAAC CAGTTCAGCG GGTGCATCAA AATGTAGAAC TGCAGATTGC TTATTCAGAG
      481 TCAACGGAAG ATCAGGTGGA GCGAATTATT GCGGAATTTA TGCAACCATT TGCTCTTGAA
      541 GTTGCTCCGT TACTTCGTGT AGGTCTTGTT AAATTGGAGG CAGAACGTCA TCTATTTATA
      601 ATGGATATGC ATCATATCAT CTCGGATGGG GTATCCATGC AGATCATGAT TCAAGAAATT
      661 GCTGATTTGT ATAAAGAAAA GGAACTTCCT ACGTTAGGCA TTCAATATAA AGACTTTACT
      721 GTTTGGCATA ATCGCTTGCT TCAATCGGAT GTTATTGAAA AACAAGAAGC TTACTGGCTG
      781 AACGTATTTG CAGAAGAGAT TCCAGTATTG AATCTACCGA CCGATTACCC AAGACCAACC
      841 ATTCAAAGCT TTGATGGTAA AAGATTTACA TTCAGTACAG GAAAGCAGCT TATGGATGAT
      901 TTATACAAGG TGGCAACAGA AACAGGAACA ACACTATATA TGGTTTTACT TGCTGCGTAT
      961 AATGTTTTCT TATCGAAGTA TTCCGGGCAA GATGACATCG TTGTAGGAAC ACCGATTGCT
     1021 GGTAGGTCTC ATGCTGATGT GGAAAATATG CTGGGGATGT TTGTAAATAC ATTAGCAATA
     1081 AGAAGTCGTT TAAATAATGA GGATACTTTT AAAGATTTTT TAGCAAATGT AAAACAAACG
     1141 GCTTTGCATG CCTATGAAAA TCCAGATTAC CCATTTGATA CGCTTGTCGA AAAGTTGGGT
     1201 ATACAGAGAG ATTTAAGTAG AAATCCATTA TTTGATACGA TGTTTGTTTT GCAAAATACG
     1261 GATAGAAAGT CTTTTGAGGT TGAACAGATA ACGATTACAC CATATGTTCC AAATAGCAGA
     1321 CATTCTAAAT TTGATCTTAC ATTAGAGGTT AGCGAAGAAC AAAATGAGAT TTTATTATGC
     1381 CTAGAATATT GCACTAAATT ATTTACGGAT AAAACAGTTG AAAGAATGGC TGGTCATTTT
     1441 TTACAGATCT TGCATGCAAT TGTTGGGAAC CCAACGATTA TAATATCAGA AATCGAGATA
     1501 TTGTCTGAAG AAGAAAAACA ACATATTTTA TTCGAGTTCA ACGATACGAA AACCACATAT
     1561 CCACATATGC AAACAATTCA AGGATTATTT GAGGAACAGG TGGAGAAGAC GCCCGACCAT
     1621 GTTGCAGTTG GATGGAAAGA CCAAACATTA ACGTATCGGG AACTTAACGA AAGAGCGAAT
     1681 CAGGTCGCAA GAGTCTTACG GCAAAAAGGA GTCCAACCCG ATAATATCGT GGGATTGCTG
     1741 GTTGAGCGTT CACCTGAAAT GCTCGTGGGT ATCATGGGAA TTCTTAAAGC AGGGGGAGCT
     1801 TATTTACCTC TTGATCCGGA GTACCCAGCG GATAGAATTT CGTACATGAT ACAAGATTGT
     1861 GGTGTACGCA TTATGCTTAC CCAACAGCAT CTTTTATCTT TAGTACATGA TGAATTTGAT
     1921 TGTGTTATTT TGGATGAAGA CAGTTTGTAC AAGGGGGATT CTTCCAATTT GGCTCCGGTT
     1981 AACCAGGCCG GGGATTTAGC CTACATCATG TACACTTCTG GTTCTACAGG AAAGCCTAAA
     2041 GGTGTTATGG TAGAACATCG AAATGTGATT CGCCTTGTGA AAAATACAAA TTATGTTCAG
     2101 GTCCGCGAAG ACGATCGTAT AATACAGACC GGAGCAATTG GATTCGATGC ACTGACATTT
     2161 GAAGTTTTTG GCTCATTGCT GCATGGAGCT GAATTGTATC CTGTTACTAA AGACGTGCTA
     2221 TTAGATGCAG AGAAACTACA CAAATTTTTA CAAGCGAATC AAATTACGAT TATGTGGTTA
     2281 ACTTCTCCGT TATTTAACCA ATTGTCACAA GGAACCGAAG AGATGTTTGC TGGCCTTCGC
     2341 TCCCTAATTG TAGGTGGAGA TGCCTTGTCT CCGAAACACA TCAATAATGT AAAGCGAAAA
     2401 TGCCCTAATC TGACTATGTG GAACGGTTAC GGCCCAACAG AAAACACCAC TTTTTCTACA
     2461 TGCTTTCTTA TTGATAAAGA ATATGATGAC AATATTCCGA TAGGGAAGGC CATTAGTAAT
     2521 TCAACAGTGT ATATCATGGA CCGGTATGGC CAGCTTCAGC CGGTGGGTGT ACCAGGAGAA
     2581 TTATGTGTAG GAGGGGATGG GGTTGCCAGG GGATATATGA ATCAGCCTGC ATTAACAGAA
```

```
2641 GAGAAGTTTG TCCCAAATCC ATTCGCTCCT GGTGAGAGAA TGTATCGCAC GGGGGATTTG
2701 GCAAGATGGT TGCCTGATGG AACAATTGAG TATTTAGGTC GTATTGATCA GCAGGTGAAA
2761 ATCAGGGGCT ACCGTATTGA ACCGGGAGAG ATTGAAACGC TTCTTGTGAA GCACAAAAAA
2821 GTCAAAGAAT CGGTAATCAT GGTAGTAGAG GATAATAATG GACAAAAGGC TCTATGCGCT
2881 TATTACGTTC CGGAAGAAGA AGTAACGGTA TCTGAACTGA GGGAATATAT AGCTAAAGAG
2941 TTGCCTGTTT ACATGGTTCC AGCCTATTTT GTACAGATTG AACAAATGCC TCTTACACAG
3001 AACGGTAAAG TAAATCGAAG CGCGTTACCA AAACCAGATG GTGAATTTGG TACAGCAACC
3061 GAATATGTAG CGCCTAGCAG CGACATTGAA ATGAAGCTGG CAGAGATTTG GCATAATGTG
3121 TTAGGGGTAA ACAAAATCGG GGTACTGGAT AACTTCTTTG AATTAGGTGG TCATTCATTA
3181 AGAGCTATGA CAATGATTTC CCAGGTACAT AAAGAGTTCG ACGTTGAATT GCCATTAAAA
3241 GTGTTATTTG AAACACCAAC GATCTCTGCA TTAGCTCAAT ACGTAACAGT ACATCTTCTT
3301 AAGGAGAAAC GTAAGCATTT CCAAGCGGGT CAAGAAACTG CACAAGCTCT GTTAAAAGGT
3361 GGATCCGCGG CAGCCCCTGG TGAAAAGCCT TATGCCTGCC CAGAGTGCGG CAAATCTTTC
3421 TCGCAACGCG CTAACCTGCG TGCCCACCAG CGCACGCATA CAGGTGAAAA ACCATATAAG
3481 TGCCCGGAGT GCGGTAAGAG CTTCAGCCGC AGTGACCACT TAACTACACA CCAACGCACG
3541 CACACGGGCG AGAAGCCGTA CAAATGCCCA GAGTGCGGGA AATCTTTTAG CCGCTCAGAT
3601 GTGCTTGTTC GCCATCAACG TACACATACG ACCTGCTTTT GCTCGCTTAA ACATCACCAT
3661 CACCATCACT AA
//
```

```
LOCUS       grsB3-Z       3708 bp ds-DNA     linear       06-FEB-2019
DEFINITION  .
FEATURES             Location/Qualifiers
     CDS             1..3708
                     /label="grsB3-Z"
     misc_feature    1..231
                     /label="docking domain Dn"
     misc_feature    3313..3390
                     /label="docking domain Dc_2"
     misc_feature    3313..3405
                     /label="linker Dc_2"
     misc_feature    232..3312
                     /label="NRPS module grsB3"
     misc_feature    3406..3666
                     /label="zinc finger domain zif268"
ORIGIN
        1 ATGAAAGATG CAGCACAAAT TGTCAATGAG GCTTTAGATC AGGGAATAAC ACTGTTCGTG
       61 GCTGAGGATC GTTTGCAATA CGAAACCCGC CTCAGCAATA TTCCGGCAGA CTTAATCAGT
      121 GAATGGAAAC AGCATAAACA AGAGTTGATT GATTTTCTGA ACCAGCTCGA TTCATCAGAA
      181 GAGCAGGTGA GTACCCATCA ACTACAGGGT ATTCCACGTT ATGAACGGGC TGATTACTAT
      241 CCAGTATCAT CTGCGCAAAA GAGGATGTAC ATCCTTTATG AATTTGAAGG GGCTGGCATT
      301 ACCTATAATG TACCTAATGT AATGTTTATA GAAGGAAAGC TGGATTATCA GCGCTTTGAA
      361 TACGCTATAA AAAGTTTGGT AAATCGACAT GAGGCGCTTC GAACGTCTTT CTATTCGCTT
      421 AATGGAGAAC CAGTTCAGCG TGTACATCAA AATGTAGAGC TACAGATTGC TTATTCGGAG
      481 GCGAAAGAAG ATGAGATAGA GCAAATTGTA GAAAGCTTTG TTCAACCATT TGACCTTGAA
      541 ATAGCTCCGC TGCTTCGCGT AGGGCTTGTT AAATTGGCAT CGGATCGCTA TTTATTCCTA
      601 ATGGATATGC ATCATATTAT CTCAGATGGT GTATCAATGC AAATTATAAC AAAAGAAATT
      661 GCCGACTTAT ATAAAGGAAA AGAGCTTGCT GAACTGCATA TTCAGTATAA AGATTTTGCT
      721 GTATGGCAAA ACGAATGGTT TCAATCTGAC GCTCTTGAAA AACAGAAAAC GTATTGGTTG
      781 AACACCTTTG CAGAGGATAT TCCGGTTTTA AATTTGTCAA CTGATTATCC AAGACCGACA
      841 ATTCAAAGTT TTGAAGGAGA TATTGTCACG TTTAGTGCAG GGAAGCAACT TGCGGAAGAA
      901 TTGAAACGCC TGGCTGCAGA AACAGGGACG ACTTTGTATA TGCTTCTGTT AGCGGCGTAC
      961 AATGTACTTT TACACAAATA CTCGGGACAG GAAGAAATTG TAGTAGGAAC GCCTATTGCC
     1021 GGGCGATCTC ACGCAGATGT GGAAAATATT GTTGGGATGT TTGTCAATAC GCTTGCATTG
     1081 AAAAATACCC CTATAGCCGT ACGCACCTTC CACGAATTCC TGTTGGAAGT AAAACAAAAT
     1141 GCTTTAGAAG CTTTTGAAAA TCAAGACTAT CCATTTGAAA ATTTGATAGA GAAGCTGCAA
     1201 GTGCGTCGCG ACTTAAGTCG CAATCCATTA TTTGATACAA TGTTTAGCCT AAGCAATATT
     1261 GACGAACAAG TAGAGATAGG GATTGAGGGA TTGAACTTCA GCCCATATGA AATGCAGTAT
     1321 TGGATTGCAA AATTTGATAT TTCATTCGAT ATTTTAGAAA AGCAAGATGA CATTCAATTT
     1381 TATTTTAACT ATTGCACGAA TCTGTTTAAA AAAGAAACGA TAGAACGATT AGCGACACAC
     1441 TTTATGCATA TTTTACAGGA GATTGTTATT AATCCTGAGA TTAAGTTATG TGAAATTAAT
     1501 ATGCTGTCCG AAGAAGAACA GCAGCGTGTC CTGTATGACT TTAATGGCAC AGATGCAACC
     1561 TACGCTACGA ATAAAATATT CCATGAGTTA TTTGAAGAAC AGGTTGAAAA AACACCAGAT
     1621 CATATAGCGG TGATAGATGA AAGAGAAAAG CTTTCCTATC AGGAGCTTAA TGCGAAAGCG
     1681 AATCAGCTGG CACGAGTGCT GCGCCAAAAA GGAGTACAGC CTAATAGCAT GGTAGGTATT
     1741 ATGGTAGATC GCTCACTCGA CATGATTGTA GGAATGCTTG GGGTTTTAAA AGCAGGAGGA
     1801 GCATATGTGC CTATCGATAT AGACTATCCT CAGGAACGGA TTAGCTACAT GATGGAAGAT
     1861 AGTGGTGCAG CGCTCTTGTT AACACAACAA AAGTTGACAC AGCAAATTGC GTTTTCTGGT
     1921 GACATTTTGT ATCTTGACCA AGAAGAATGG CTTCATGAGG AAGCTTCAAA TTTAGAACCC
     1981 ATCGCTCGTC CGCAGGATAT AGCCTATATC ATTTACACTT CTGGTACAAC CGGAAAGCCA
     2041 AAAGGTGTGA TGATTGAGCA TCAAAGCTAT GTGAATGTAG CAATGGCATG GAAAGATGCC
     2101 TATCGGTTAG ATACATTCCC GGTCCGTTTG CTTCAGATGG CTAGCTTTGC CTTTGACGTA
     2161 TCTGCAGGTG ATTTTGCCAG AGCACTACTT ACAGGTGGGC AATTAATTGT ATGTCCAAAT
     2221 GAAGTAAAGA TGGACCCAGC TTCTTTATAT GCCATTATTA AGAAATATGA CATTACTATT
     2281 TTTGAAGCAA CGCCTGCTCT AGTGATTCCA TTGATGGAGT ATATTTATGA ACAGAAGCTG
     2341 GATATTAGCC AGTTACAGAT TCTGATTGTC GGATCGGACA GTTGTTCGAT GGAAGACTTT
     2401 AAAACCTTGG TTTCCCGTTT TGGTTCAACT ATACGTATTG TGAATAGCTA TGGAGTAACC
     2461 GAAGCGTGCA TTGATTCTAG CTATTATGAA CAACCGCTTT CTTCGTTACA TGTAACAGGA
     2521 ACTGTACCGA TTGGAAAACC GTACGCTAAC ATGAAAATGT ATATTATGAA TCAATATTTG
     2581 CAGATTCAGC CTGTAGGTGT AATTGGAGAA TTATGTATTG GAGGAGCCGG GGTTGCCCGT
```

```
2641 GGATATTTAA ATAGACCGGA CTTAACAGCA GAAAAGTTTG TCCCTAATCC TTTTGTTCCA
2701 GGTGAAAAGC TGTATCGAAC AGGCGACTTG GCAAGATGGA TGCCGGATGG GAATGTTGAG
2761 TTTCTTGGTC GAAATGACCA TCAGGTGAAA ATCAGAGGGA TTCGAATCGA GCTTGGAGAA
2821 ATCGAAGCAC AACTGCGTAA ACATGATAGC ATAAAAGAAG CAACTGTGAT CGCAAGAGAA
2881 GATCACATGA AAGAGAAATA TTTATGTGCG TATATGGTGA CCGAAGGAGA AGTAAATGTA
2941 GCTGAACTGC GTGCGTATCT AGCAAATGAT CTGCCTGCGG CAATGATTCC GTCATATTTT
3001 GTATCGCTCG AAGCAATGCC ACTTACTGCT AATGGAAAAA TTGATAAGCG ATCTTTACCA
3061 GAGCCCGATG GTTCCATATC GATAGGAACA GAATATGTAG CTCCGCGTAC CATGCTTGAG
3121 GGAAAACTAG AAGAGATATG GAAAGATGTA TTGGGTTTAC AGCGTGTTGG CATTCACGAT
3181 GACTTCTTTA CAATAGGTGG CCATTCATTG AAGGCTATGG CTGTTATTTC GCAAGTTCAT
3241 AAAGAATGCC AGACTGAAGT TCCTCTGCGT GTCTTATTTG AAACACCTAC CATTCAAGGA
3301 CTGGCTAAAT ATGTAACAGT ACATCTTCTT AAGGAGAAAC GTAAGCATTT CCAAGCGGGT
3361 CAAGAAACTG CACAAGCTCT GTTAAAAGGT GGATCCGCGG CAGCCCCTGG TGAAAAGCCT
3421 TATGCCTGCC CAGTGGAAAG CTGTGACCGC CGTTTCTCTC GTTCAGACGA ACTTACTCGT
3481 CACATCCGCA TCCACACGGG ACAGAAGCCC TTCCAATGTC GTATTTGTAT GCGCAATTTC
3541 AGCCGCTCAG ACCATCTGAC TACTCATATC CGCACGCACA CCGGGGAAAA ACCCTTTGCT
3601 TGTGACATCT GTGGTCGTAA ATTTGCCCGT AGTGACGAGC GTAAGCGCCA CACCAAAATT
3661 CACACAACCT GCTTTTGCTC GCTTAAACAT CACCATCACC ATCACTAA
//
```

```
LOCUS       B-grsB4             4467 bp ds-DNA     linear       06-FEB-2019
DEFINITION  .
FEATURES             Location/Qualifiers
     misc_feature    4..258
                     /label="zinc finger domain zfb"
     misc_feature    502..4434
                     /label="NRPS module grsB4"
     misc_feature    271..501
                     /label="docking domain Dn"
     CDS             1..4467
                     /label="B-grsB4"
ORIGIN
        1 ATGCCTGGTG AAAAGCCTTA TAAATGTCCC GAATGTGGGA AGAGCTTTAG CCGTAGCGAC
       61 AAACTGGTTC GGCACCAACG CACCCATACC GGTGAAAAGC CGTATAAATG CCCGGAATGT
      121 GGCAAATCCT TTTCGACCTC TGGCGAATTA GTCCGCCATC AGCGCACTCA CACGGGTGAG
      181 AAACCGTACA AATGCCCAGA ATGCGGCAAA TCGTTCAGTC GCTCAGATGA TCTGGTGCGT
      241 CATCAGCGTA CACACACCGG CGGTGGGAGC ATGAAAGATG CAGCACAAAT TGTCAATGAG
      301 GCTTTAGATC AGGGAATAAC ACTGTTCGTG GCTGAGGATC GTTTGCAATA CGAAACCCGC
      361 CTCAGCAATA TTCCGGCAGA CTTAATCAGT GAATGGAAAC AGCATAAACA AGAGTTGATT
      421 GATTTTCTGA ACCAGCTCGA TTCATCAGAA GAGCAGGTGA GTACCCATCA ACTACAGGGT
      481 ATTCCACGTT ATGAACGGGC TGACTATTAT CCAGTATCAT CAGCACAAAA GAGAATGTTT
      541 ATTGTTAATC AATTTGATGG AGTAGGAATT AGCTACAATA TGCCTTCCAT CATGCTGATT
      601 GAAGGAAAAC TTGAGCGAAC ACGCTTGGAA TCAGCATTTA AAAGATTGAT AGAACGACAT
      661 GAGAGCCTTC GAACATCTTT TGAAATAATA AATGGTAAGC CTGTACAGAA GATTCATGAG
      721 GAAGTTGATT TCAATATGTC CTATCAGGTG GCTTCTAATG AACAAGTAGA GAAGATGATC
      781 GATGAGTTCA TTCAGCCTTT CGATTTAAGT GTTGCACCGC TGCTTCGTGT GGAACTTTTA
      841 AAATTGGAAG AAGACCGTCA TGTGCTTATA TTTGATATGC ATCATATTAT CTCAGATGGT
      901 ATATCTTCCA ATATTTTGAT GAAAGAATTA GGAGAACTAT ATCAAGGTAA TGCTTTACCA
      961 GAACTTCGTA TTCAATACAA GGATTTCGCT GTATGGCAAA ATGAGTGGTT CCAGTCAGAA
     1021 GCCTTTAAAA AGCAAGAAGA ATACTGGGTA AATGTCTTCG CAGATGAACG CCCGATTCTG
     1081 GATATACCGA CGGATTATCC AAGGCCGATG CAACAAAGCT TTGATGGTGC TCAACTTACA
     1141 TTTGGAACCG GAAAGCAGCT TATGGATGGG TTATACAGGG TAGCAACGGA AACGGGAACA
     1201 ACGCTTTATA TGGTTTTGCT TGCGGCATAT AATGTTCTTC TTTCCAAATA TTCTGGTCAA
     1261 GAAGATATTA TTGTAGGGAC ACCGATTGTG GGTAGATCCC ATACTGACCT TGAGAATATT
     1321 GTCGGGATGT TTGTCAACAC GTTAGCAATG AGAAATAAAC CGGAAGGAGA AAAGACGTTC
     1381 AAAGCATTTG TATCAGAAAT AAAGCAGAAT GCACTAGCGG CTTTTGAGAA TCAGGATTAT
     1441 CCATTTGAAG AGCTTATCGA AAAACTAGAG ATACAAAGGG ACTTAAGCAG AAATCCATTA
     1501 TTTGATACGC TCTTTAGCCT TCAAAACATA GGTGAAGAAT CATTTGAACT AGCCGAATTA
     1561 ACATGCAAAC CTTTCGATTT GGTAAGCAAA TTAGAGCATG CCAAGTTTGA TCTGAGTCTT
     1621 GTGGCAGTAG AAAAAGAGGA AGAAATTGCA TTTGGGCTTC AATACTGCAC AAAACTGTAT
     1681 AAGGAAAAAA CAGTTGAACA ACTGGCTCAA CATTTTATTC AAATAGTAAA AGCAATTGTA
     1741 GAAAATCCAG ATGTCAAATT ATCTGATATT GATATGTTAT CTGAAGAAGA GAAGAAACAA
     1801 ATCTTACTTG AGTTCAATGA TACGAAAATA CAATATCCGC AGAATCAAAC AATACAGGAA
     1861 TTGTTTGAAG AGCAAGTGAA GAAAACACCT GAACATATAG CAATCGTATG GGAAGGGCAA
     1921 GCATTAACCT ATCATGAGCT AAATATAAAA GCTAATCAGT TAGCTCGTGT ATTACGAGAA
     1981 AAAGGGGTAA CCCCTAATCA TCCTGTAGCG ATTATGACGG AACGCTCATT AGAGATGATC
     2041 GTAGGTATCT TTAGTATTTT GAAAGCAGGA GGAGCATATG TTCCAATTGA TCCAGCCTAT
     2101 CCACAAGAAC GTATTCAATA CTTGCTTGAA GATAGCGGAG CGACGCTACT GCTTACTCAG
     2161 TCACATGTAT TAAATAAATT ACCGGTCGAT ATCGAATGGT TGGATCTTAC AGATGAACAA
     2221 AACTATGTAG AAGATGGTAC CAATCTTCCA TTTATGAATC AGTCAACAGA TCTTGCCTAT
     2281 ATTATTTATA CATCCGGTAC AACAGGCAAG CCTAAAGGGG TTATGATTGA ACATCAAAGC
     2341 ATCATCAACT GCCTGCAATG GCGGAAGGAA GAATACGAAT TTGGACCAGG GGATACGGCT
     2401 CTACAAGTGT TTTCCTTTGC TTTTGATGGA TTTGTAGCAA GTTTGTTTGC TCCGATTCTT
     2461 GCAGGTGCAA CGTCTGTTCT CCCTAAGGAG GAAGAAGCAA AAGATCCAGT TGCATTGAAA
     2521 AAACTGATCG CATCAGAAGA GATTACACAT TACTACGGTG TGCCTAGTTT GTTTAGTGCC
     2581 ATTCTTGATG TTTCTTCTAG TAAGGATTTG CAAAATTTAC GCTGCGTCAC TTTGGGAGGA
     2641 GAGAAATTAC CGGCTCAAAT TGTTAAAAAA ATCAAAGAAA AAAATAAAGA AATTGAAGTC
     2701 AACAACGAAT ATGGGCCTAC TGAAAATAGT GTAGTAACTA CTATTATGCG CGATATACAG
     2761 GTAGAACAAG AGATTACTAT TGGTCGCCCA TTATCTAACG TAGATGTATA TATTGTCAAT
     2821 TGTAATCATC AATTACAACC AGTAGGTGTA GTAGGGGAAT TATGTATTGG TGGACAGGGA
```

```
2881 CTTGCAAGAG GATATTTGAA TAAACCAGAG CTTACAGCAG ATAAATTTGT TGTAAATCCA
2941 TTCGTACCTG GTGAACGTAT GTACAAAACC GGTGACCTTG CAAAATGGCG CTCAGATGGA
3001 ATGATTGAAT ATGTGGGGCG TGTTGATGAA CAAGTAAAAG TAAGAGGATA TCGGATTGAG
3061 CTTGGTGAAA TTGAATCAGC TATCCTAGAA TACGAAAAAA TTAAGGAAGC GGTAGTTATG
3121 GTTTCGGAGC ATACTGCATC TGAACAGATG TTATGTGCTT ATATTGTAGG GGAAGAAGAT
3181 GTACTGACTC TGGACTTAAG AAGCTATCTA GCAAAATTAC TACCAAGTTA TATGATTCCA
3241 AACTATTTTA TCCAATTGGA TAGTATTCCG CTTACACCAA ACGGTAAAGT GGATCGTAAA
3301 GCATTGCCTG AACCTCAAAC CATTGGCTTA ATGGCAAGGG AGTATGTTGC ACCAAGGAAT
3361 GAAATCGAAG CACAGCTAGT ACTCATTTGG CAAGAGGTAT TAGGAATAGA ACTGATCGGT
3421 ATTACCGATA ATTTCTTTGA ATTAGGAGGG CATTCTTTAA AGGCAACGCT TTTAGTTGCA
3481 AAAATTTACG AGTACATGCA AATAGAGATG CCATTAAATG TTGTGTTTAA ACATTCAACT
3541 ATTATGAAAA TAGCGGAATA TATTACACAT CAAGAATCAG AAAATAATGT ACATCAGCCT
3601 ATTTTGGTAA ATGTAGAAGC AGATAGAGAG GCGCTATCTC TTAACGGCGA GAAGCAAAGA
3661 AAAAATATAG AGCTACCTAT TCTGCTAAAC GAAGAAACAG ATCGAAACGT ATTCTGCTTC
3721 GCGCCCATTG GTGCACAAGG TGTTTTTTAT AAAAAGCTTG CTGAACAAAT CCCTACTGCA
3781 TCCTTGTATG GCTTTGACTT CATTGAAGAT GATGATCGAA TTCAGCAATA TATTGAATCG
3841 ATGATTCAAA CTCAGTCAGA CGGACAATAT GTGCTAATTG GTTATTCTTC AGGAGGGAAC
3901 CTGGCTTTTG AAGTAGCAAA AGAAATGGAA AGGCAAGGAT ATAGTGTATC TGATTTGGTC
3961 TTGTTCGATG TTTACTGGAA GGGAAAAGTC TTCGAGCAAA CAAAAGAAGA AGAAGAAGAA
4021 AACATAAAAA TAATAATGGA AGAATTAAGG GAAAATCCAG GAATGTTCAA TATGACACGA
4081 GAGGATTTTG AACTGTATTT TGCGAATGAA TTTGTGAAAC AAAGTTTCAC ACGGAAAATG
4141 CGCAAATACA TGAGTTTTTA TACGCAGTTA GTTAATTATG GGGAAGTAGA AGCTACAATT
4201 CACCTTATAC AAGCAGAATT TGAGGAAGAA AAAATTGACG AAAACGAAAA AGCCGACGAA
4261 GAAGAAAAAA CATATCTAGA GGAAAAATGG AATGAAAAAG CATGGAACAA AGCAGCAAAA
4321 AGATTTGTAA AATATAACGG ATATGGCGCT CATTCTAACA TGCTAGGAGG TGATGGTTTA
4381 GAGAGAAATT CCTCTATCCT TAAACAGATA CTACAAGGGA CATTTGTAGT AAAAGGATCC
4441 AGATCTCATC ACCATCACCA TCACTAA
//
```

**Table S3. Linkers connecting NRPS module and zinc finger domain.**

| Linker | Peptide sequence | Description (underlined) |
|---|---|---|
| linker o | VEGGGESAYLAIPQAEGSAAA | TycB-derived, original linker following TycB1 |
| linker com | IKDRSELTPSDFSFGSRSSSTSAAA | GrsA-derived, designed in reference to pTrcHis-tycA::COM$^{D}$ΔE [13] |
| Dn | MKDAAQIVNEALDQGITLFVAEDRLQYETRLSNIPADLISE WKQHKQELIDFLNQLDSSEEQVSTHQLQGIPRYERA | N-terminus of InxB |
| linker Dc | VTVHLLKEKRKHFQAGQETAQALLKGDIGSAAA | C-terminus of InxA |
| linker Dc_1 | VTVHLLKEKRKHFQAGQETAQALLKGDGSAAA | Truncated C-terminus of InxA |
| linker Dc_2 | VTVHLLKEKRKHFQAGQETAQALLKGGSAAA | Truncated C-terminus of InxA |
| linker Dc_4 | VTVHLLKEKRKHFQAGQETAQALLGSAAA | Truncated C-terminus of InxA |
| linker Dc_5 | VTVHLLKEKRKHFQAGQETAQALGSAAA | Truncated C-terminus of InxA |
| linker Dc_10 | VTVHLLKEKRKHFQAGQEGSAAA | Truncated C-terminus of InxA |
| linker Dc_19 | VTVHLLKEKGSAAA | Truncated C-terminus of InxA |

**Table S4. DNA templates and probes.**[#]

| Oligos | Sequence (5' to 3') |
|---|---|
| | **Fluorescence polarization assay** |
| **z_F** | tacaGCGTGGGCGtaat |
| **z_R (FdT)** | at[FdT]aCGCCCACGCtgta |
| **p_F** | ggacGTGTGGAAAaacg |
| **p_R (FdT)** | cg[FdT]tTTTCCACACgtcc |
| **n_F** | ggacAAGGGTTCAgatg |
| **n_R (FdT)** | ca[FdT]cTGAACCCTTgtcc |
| **b_F** | tacaGCGGCTGGGtaat |
| **b_R (FdT)** | at[FdT]aCCCAGCCGCtgta |
| | **DNA spacer test** |
| **zp9_F** | ggtaccGCGTGGGCGGTGTGGAAAtattc |
| **zp9_R** | gaataTTTCCACACCGCCCACGCggtacc |
| **zp10_F** | ggtaccGCGTGGGCGtGTGTGGAAAtattc |
| **zp10_R** | gaataTTTCCACACaCGCCCACGCggtacc |
| **zp12_F** | ggtaccGCGTGGGCGtgaGTGTGGAAAtattc |
| **zp12_R** | gaataTTTCCACACtcaCGCCCACGCggtacc |
| **zp14_F** | ggtaccGCGTGGGCGtgaaaGTGTGGAAAtattc |
| **zp14_R** | gaataTTTCCACACtttcaCGCCCACGCggtacc |
| **zp16_F** | ggtaccGCGTGGGCGtgattaaGTGTGGAAAtattc |
| **zp16_R** | gaataTTTCCACACttaatcaCGCCCACGCggtacc |
| **zp18_F** | ggtaccGCGTGGGCGtgatcataaGTGTGGAAAtattc |
| **zp18_R** | gaataTTTCCACACttatgatcaCGCCCACGCggtacc |
| **zp19_F** | ggtaccGCGTGGGCGtgatccataaGTGTGGAAAtattc |
| **zp19_R** | gaataTTTCCACACttatggatcaCGCCCACGCggtacc |
| **zp20_F** | ggtaccGCGTGGGCGtgatccaataaGTGTGGAAAtattc |
| **zp20_R** | gaataTTTCCACACttattggatcaCGCCCACGCggtacc |
| **zp21_F** | ggtaccGCGTGGGCGtgatcctaataaGTGTGGAAAtattc |
| **zp21_R** | gaataTTTCCACACttattaggatcaCGCCCACGCggtacc |
| **zp22_F** | ggtaccGCGTGGGCGtgatccttaataaGTGTGGAAAtattc |
| **zp22_R** | gaataTTTCCACACttattaaggatcaCGCCCACGCggtacc |
| **zp24_F** | ggtaccGCGTGGGCGtgatcctgataataaGTGTGGAAAtattc |
| **zp24_R** | gaataTTTCCACACttattatcaggatcaCGCCCACGCggtacc |
| **zp26_F** | ggtaccGCGTGGGCGtgatcctgccataataaGTGTGGAAAtattc |
| **zp26_R** | gaataTTTCCACACttattatggcaggatcaCGCCCACGCggtacc |
| **zp28_F** | ggtaccGCGTGGGCGtgatcctgccccataataaGTGTGGAAAtattc |
| **zp28_R** | gaataTTTCCACACttattatggggcaggatcaCGCCCACGCggtacc |
| **zp30_F** | ggtaccGCGTGGGCGtgatcctgcccgccataataaGTGTGGAAAtattc |
| **zp30_R** | gaataTTTCCACACttattatggcgggcaggatcaCGCCCACGCggtacc |
| **zp31_F** | ggtaccGCGTGGGCGtgatcctgcccagccataataaGTGTGGAAAtattc |

| | |
|---|---|
| **zp31_R** | gaataTTTCCACACttattatggctgggcaggatcaCGCCCACGCggtacc |
| **zp32_F** | ggtaccGCGTGGGCGtgatcctgcccacgccataataaGTGTGGAAAtattc |
| **zp32_R** | gaataTTTCCACACttattatggcgtgggcaggatcaCGCCCACGCggtacc |
| **zp33_F** | ggtaccGCGTGGGCGtgatcctgcccatcgccataataaGTGTGGAAAtattc |
| **zp33_R** | gaataTTTCCACACttattatggcgatgggcaggatcaCGCCCACGCggtacc |
| **zp34_F** | ggtaccGCGTGGGCGtgatcctgcccatccgccataataaGTGTGGAAAtattc |
| **zp34_R** | gaataTTTCCACACttattatggcggatgggcaggatcaCGCCCACGCggtacc |
| **zp36_F** | ggtaccGCGTGGGCGtgatcctgcccatgtccgccataataaGTGTGGAAAtattc |
| **zp36_R** | gaataTTTCCACACttattatggcggacatgggcaggatcaCGCCCACGCggtacc |
| **zp40_F** | ggtaccGCGTGGGCGtgatcctgcccatgcgtctccgccataataaGTGTGGAAAtattc |
| **zp40_R** | gaataTTTCCACACttattatggcggagacgcatgggcaggatcaCGCCCACGCggtacc |
| **zp44_F** | ggtaccGCGTGGGCGtgatcctgcccatgcgcacgtctccgccataataaGTGTGGAAAtattc |
| **zp44_R** | gaataTTTCCACACttattatggcggagacgtgcgcatgggcaggatcaCGCCCACGCggtacc |
| | **Multimodular templating** |
| **zpn9_F** | gtacaGCGTGGGCGGTGTGGAAAAAGGGTTCAgctgg |
| **zpn9_R** | ccagcTGAACCCTTTTTCCACACCGCCCACGCtgtac |
| **zpn20_F** | gtacaGCGTGGGCGtgatcctgcccGTGTGGAAAactcccaggacAAGGGTTCAgctgg |
| **zpn20_R** | ccagcTGAACCCTTgtcctgggagtTTTCCACACgggcaggatcaCGCCCACGCtgtac |
| **zpn32_F** | gtacaGCGTGGGCGtgatcctgcccgccataatagccGTGTGGAAAaacatgcgcacgtctcccaggacAAGGGTTCA gctgg |
| **zpn32_R** | ccagcTGAACCCTTgtcctgggagacgtgcgcatgttTTTCCACACggctattatggcgggcaggatcaCGCCCACGC tgtac |
| **b'zpn_F** | gtacaCCCAGCCGCtGCGTGGGCGtgatcctgcccGTGTGGAAAactcccaggacAAGGGTTCAgctgg |
| **b'zpn_R** | ccagcTGAACCCTTgtcctgggagtTTTCCACACgggcaggatcaCGCCCACGCaGCGGCTGGGtgtac |
| | **DNA spacer test and direction test** |
| **p'z9_F** | gaataCGCCCACGCGTGTGGAAAagtacc |
| **p'z9_R** | ggtactTTTCCACACGCGTGGGCGtattc |
| **p'z10_F** | gaataCGCCCACGCtGTGTGGAAAagtacc |
| **p'z10_R** | ggtactTTTCCACACaGCGTGGGCGtattc |
| **p'z11_F** | ggtactTTTCCACACgaGCGTGGGCGtattc |
| **p'z11_R** | gaataCGCCCACGCtcGTGTGGAAAagtacc |
| **p'z13_F** | gaataCGCCCACGCtaccGTGTGGAAAagtacc |
| **p'z13_R** | ggtactTTTCCACACggtaGCGTGGGCGtattc |
| **pz20_F** | ggtaccGTGTGGAAAagatccaataaGCGTGGGCGtattc |
| **pz20_R** | gaataCGCCCACGCttattggatctTTTCCACACggtacc |
| **pz32_F** | ggtaccGTGTGGAAAagatcctgcccacgccataataaGCGTGGGCGtattc |
| **pz32_R** | gaataCGCCCACGCttattatggcgtgggcaggatctTTTCCACACggtacc |
| | **SEC*** |
| **SEC-DNA_F** | gtacaCCCAGCCGCtGCGTGGGCGtgatcctgcccGTGTGGAAAactcccaggacAAGGGTTCAgctgg |
| **SEC-DNA_R** | ccagcTGAACCCTTgtcctgggagtTTTCCACACgggcaggatcaCGCCCACGCaGCGGCTGGGtgtac |

#ZF domain binding sites are shown in capital letters. The linkers in between are random sequences (lower case). [FdT]: fluorescein-dT; _F: forward oligo; _R: reverse oligo. A prime after the ZF abbreviation indicates an inverted direction of the binding site. ***The** SEC-DNA contains a fourth zinc finger binding site not used in this work.

**Table S5. Tuning parameters for peptide quantification by tandem mass spectrometry.**

| Compound | Retention time (min) | Parent ion (m/z) | Daughter ion (m/z) | Collision energy (V) | Concentrations tested[#] |
|---|---|---|---|---|---|
| **cyclo-(fP)** | 1.78 | 245.1 | 70.1 | 18 | 10 nM – 1μM |
| **fPO*** | 1.59 | 359.2 | 212.1 | 18 | 20 pM – 200 nM |
| **fPVO*** | 1.69 | 458.3 | 245.1 | 21 | 200 pM – 200 nM |
| **fPVOL** | 1.73 | 295.2 | 120.1 | 18 | 172 pM – 172 nM |

[#]In this concentration range, the signal was linear but the full dynamic range may be larger. Only relevant concentrations were tested.

**Table S6. R script for modelling bimolecular binding.**

```
# Input data are organized in three columns in a csv (comma delimited) file
# 1st column starts with title "A0", the following rows specify DNA concentrations
# 2nd column starts with title "P0", the following rows specify protein concentration
# a data point without protein is required to calculate F0
# 3rd column starts with title "F", the following rows specify fluorescence polarization

library(dplyr)
library(stringr)

# Load input data
dat1 <- read.csv(file=file.choose())
F0 = dat1[dat1$P0==0,3]
print(dat1)

# Single-ligand model, requires preassigned variable F0
lig1 <- function(P0,A0,Ka,Fmax) {
  return((P0+A0+Ka-((P0+A0+Ka)^2-4*P0*A0)^0.5)/(2*A0)*(Fmax-F0)+F0)
}
# fit to model 1
m1<-nls(F~lig1(P0, A0, Ka, Fmax),data=dat1, start = list(Ka=20,Fmax = max(dat1$F)))
summary(m1)

# plot original data in log scale
plot.new()
x_label<- "P0 (nM)"; y_label<- "F (mP)"
plot(dat1[dat1$P0!=0,]$P0, dat1[dat1$P0!=0,]$F, log="x", xlab=x_label, ylab=y_label)

# draw regression line
Pseq=seq(log10(min(dat1[dat1$P0!=0,]$P0)),log10(max(dat1[dat1$P0!=0,]$P0)),len = 100)
Pseq=10^Pseq
lines(Pseq,predict(m1,list(P0= Pseq, A0=dat1$A0[1])),col="blue",lwd=2)
abline(h=F0, lty=3)
```

**Table S7. Quantification of peptides in tetramodular spacer adjustment (related to Figure 5).#**

| Experiment | Cyclo-(fP) (nM) | Std. dev. | fPO* (nM) | Std. dev. | fPVO* (nM) | Std. dev. | fPO*/ cyclo-[fP] (%) | Std. dev. | fPVO*/ cyclo-[fP] (%) | Std. dev. |
|---|---|---|---|---|---|---|---|---|---|---|
| **No DNA** | 8860 | 397 | 25 | 3 | 20 | 3 | 0.28 | 0.02 | 0.22 | 0.02 |
| **9-bp spacer** | 8386 | 629 | 106 | 19 | 1256 | 228 | 1.25 | 0.13 | 14.92 | 1.60 |
| **20-bp spacer** | 8635 | 651 | 34 | 5 | 868 | 161 | 0.40 | 0.03 | 10.00 | 1.11 |
| **32-bp spacer** | 8887 | 577 | 23 | 4 | 512 | 70 | 0.25 | 0.02 | 5.75 | 0.41 |
| **GrsB123** | 4564 | 348 | 8 | 3 | 1336 | 377 | 0.18 | 0.05 | 29.04 | 6.05 |

#Concentrations are the average of two independent measurements.

### NMR spectra

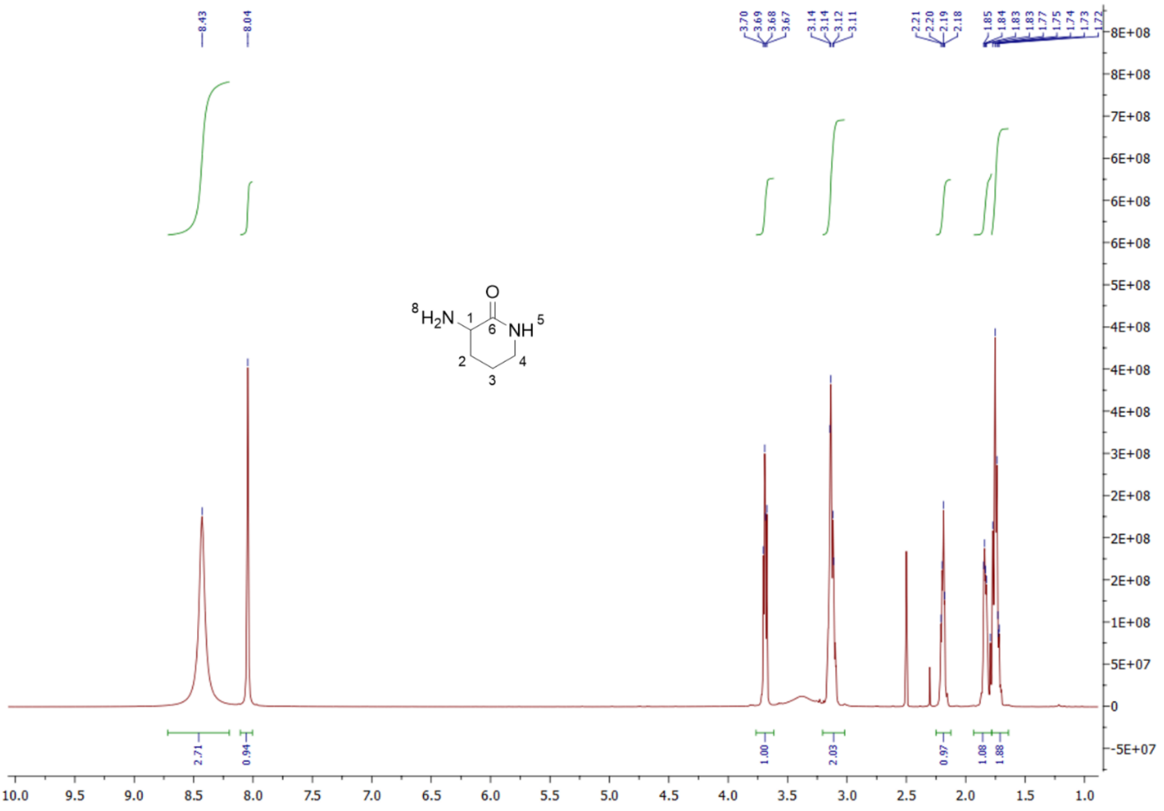

$^1$H NMR of compound **2** measured in DMSO-$d_6$

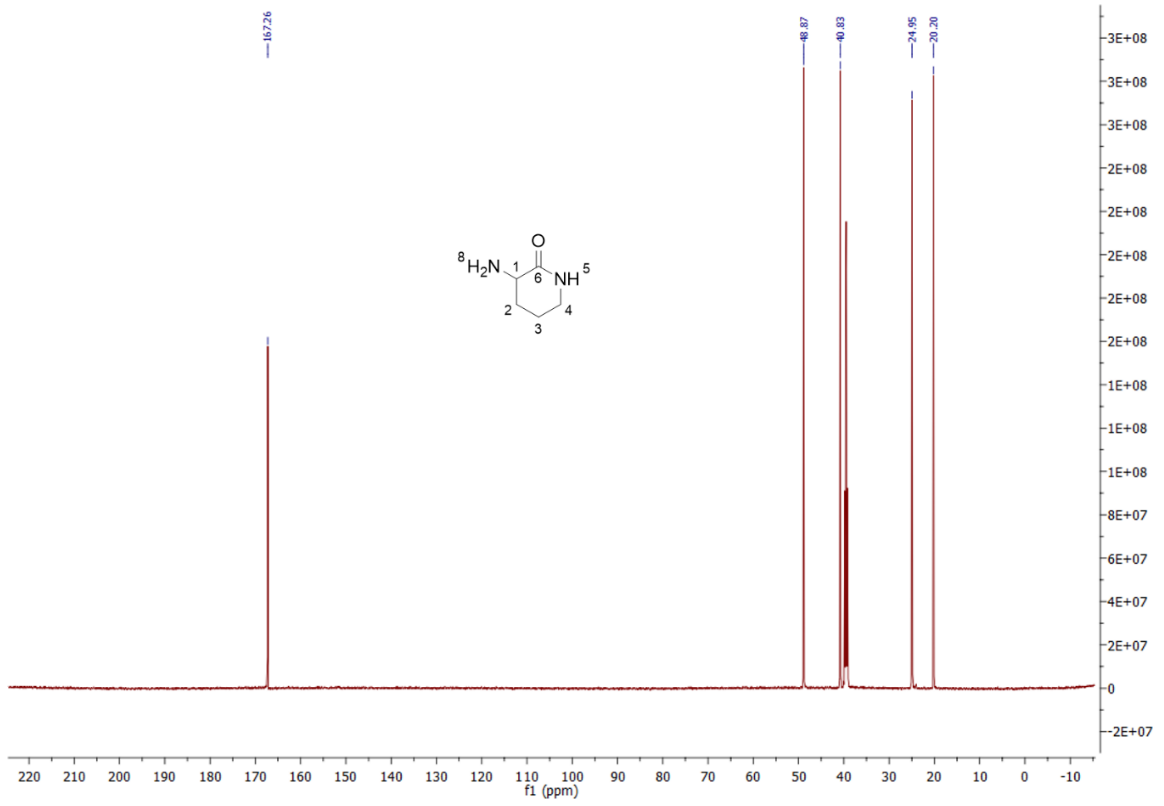

$^{13}$C NMR of compound **2** measured in DMSO-$d_6$

| Position | $\delta_C$ | $\delta_H$, mult. (*J* in Hz) |
|---|---|---|
| 1 | 48.9, CH | 3.69, dd (11.0, 6.1) |
| 2a | 25.0, $CH_2$ | 2.27 – 2.11, m |
| 2b | | 1.89 – 1.80, m |
| 3 | 20.2, $CH_2$ | 1.78 – 1.69, m |
| 4 | 40.8, $CH_2$ | 3.19 – 3.04, m |
| 5 | | 8.04, s |
| 6 | 167.3, qC | |
| 8 | | 8.43, broad s |

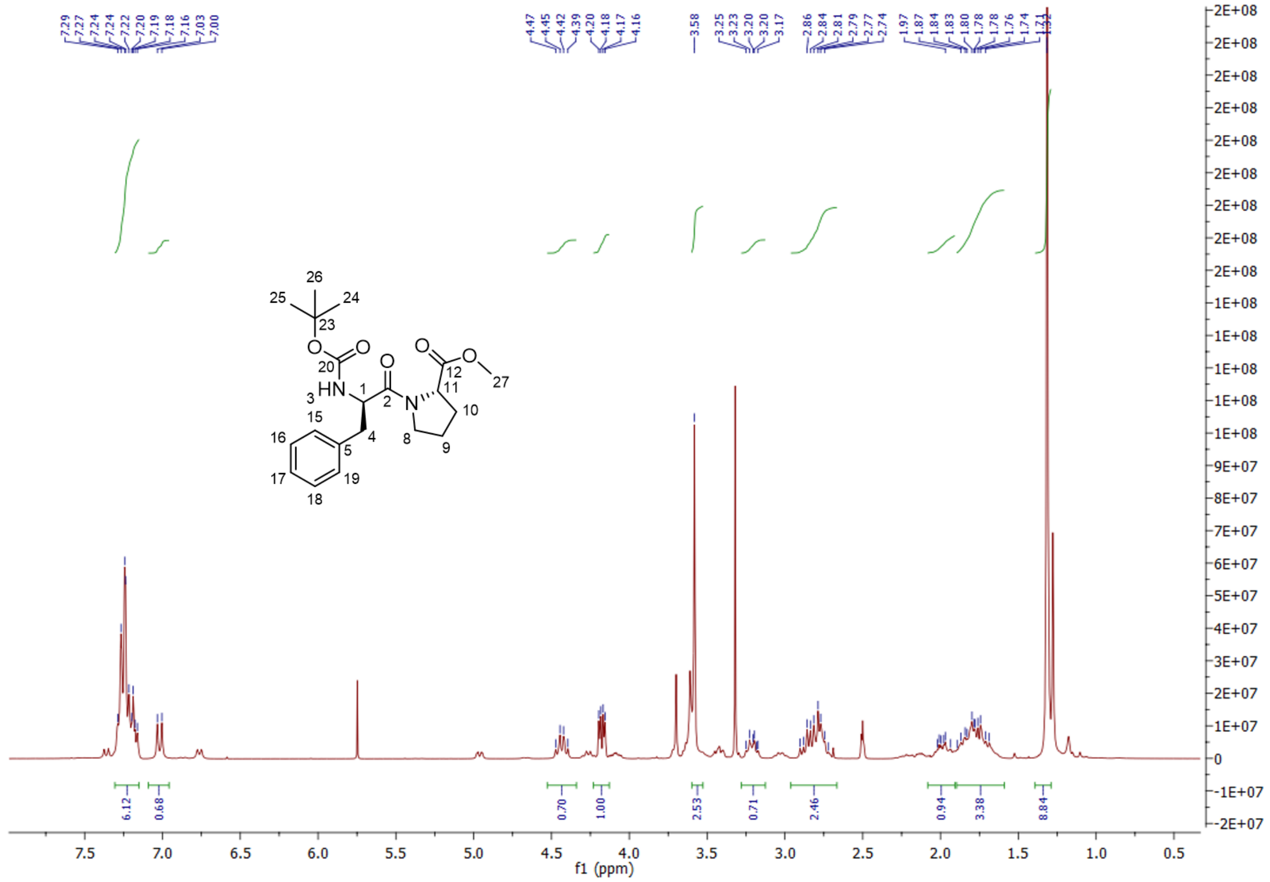

[1]H NMR of compound **5** measured in DMSO-$d_6$

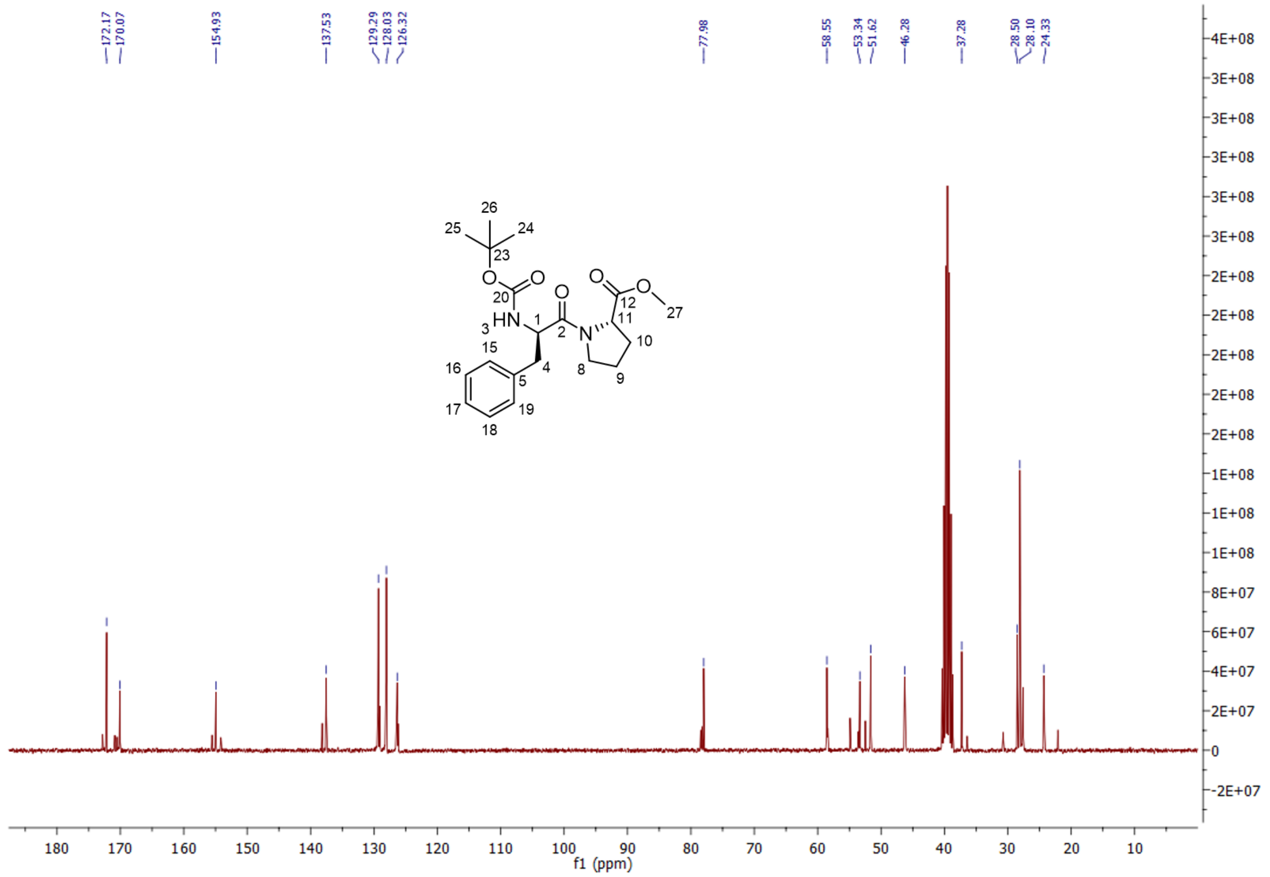

[13]C NMR of compound **5** measured in DMSO-$d_6$

| Position | $\delta_C$, | $\delta_H$, mult. (*J* in Hz) |
|---|---|---|
| 1 | 53.3, CH | 4.43, dd (15.1, 8.5) |
| 2 | 170.1, qC | |
| 3 | | 7.02, d (8.7) |
| 4 | 37.3, $CH_2$ | 2.92 – 2.67, m |
| 5 | 137.5, qC | |
| 8a<br>8b | 46.3, $CH_2$ | 3.28 – 3.12, m<br>2.92 – 2.67, m |
| 9 | 24.3, $CH_2$ | 1.90 – 1.59, m |
| 10a<br>10b | 28.5, $CH_2$ | 2.06 – 1.93, m<br>1.90 – 1.59, m |
| 11 | 58.6, CH | 4.18, dd (7.7, 4.3) |
| 12 | 172.2, qC | |
| 15 | 128.0, CH | 7.31 – 7.13, m |
| 16 | 129.3, CH | 7.31 – 7.13, m |
| 17 | 126.3, CH | 7.31 – 7.13, m |
| 18 | 129.3, CH | 7.31 – 7.13, m |
| 19 | 128.0, CH | 7.31 – 7.13, m |
| 20 | 154.9, qC | |
| 23 | 78.0, qC | |
| 24 | 28.1, $CH_3$ | 1.32, s |
| 25 | 28.1, $CH_3$ | 1.32, s |
| 26 | 28.1, $CH_3$ | 1.32, s |
| 27 | 51.6, $CH_3$ | 3.58, s |

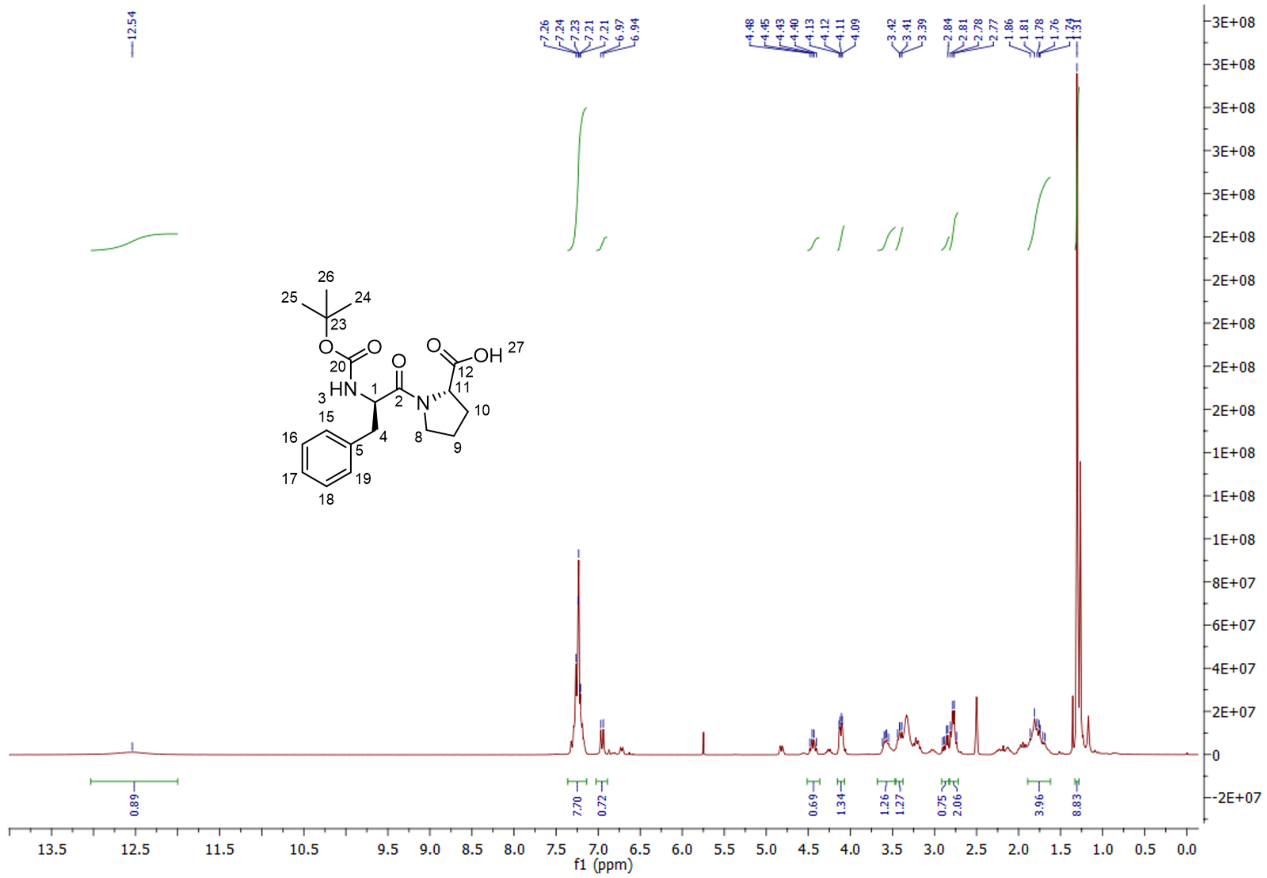

[1]H NMR of compound **6** measured in DMSO-$d_6$

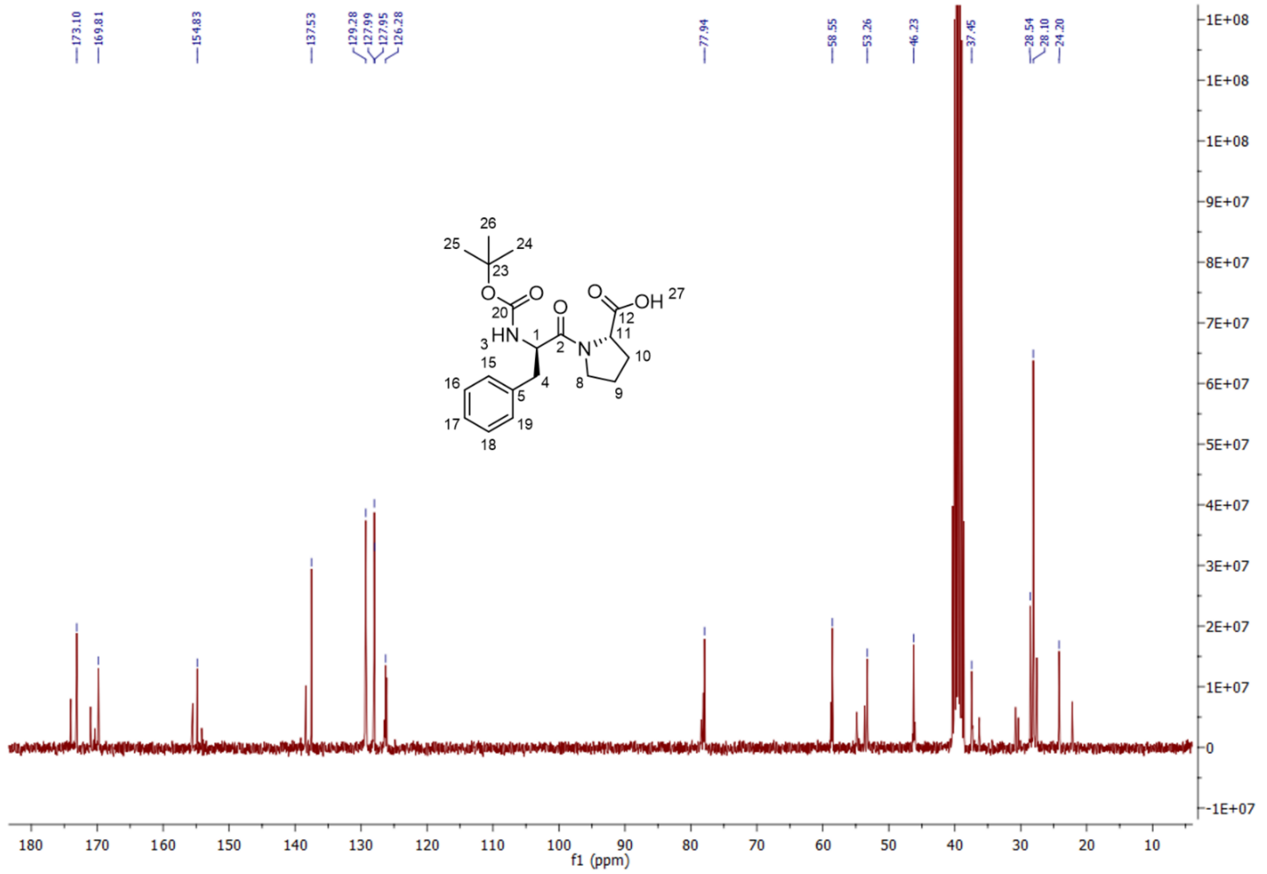

[13]C NMR of compound **6** measured in DMSO-$d_6$

| **Position** | **$\delta_C$** | **$\delta_H$, mult. (*J* in Hz)** |
|---|---|---|
| 1 | 53.3, CH | 4.44, dd (15.0, 8.3) |
| 2 | 169.8, qC | |
| 3 | | 6.95, d (8.8) |
| 4 | 37.5, $CH_2$ | 2.92 – 2.71, m |
| 5 | 137.5, qC | |
| 8a<br>8b | 46.2, $CH_2$ | 3.27 – 3.13, m<br>2.92 – 2.71, m |
| 9 | 24.2, $CH_2$ | 1.88 – 1.64, m |
| 10a<br>10b | 28.5, $CH_2$ | 2.06 – 1.88, m<br>1.88 – 1.64, m |
| 11 | 58.6, CH | 4.11, dd (8.3, 3.5) |
| 12 | 173.1, qC | |
| 15 | 128.0, CH | 7.35 – 7.10, m |
| 16 | 129.3, CH | 7.35 – 7.10, m |
| 17 | 126.3, CH | 7.35 – 7.10, m |
| 18 | 129.3, CH | 7.35 – 7.10, m |
| 19 | 128.0, CH | 7.35 – 7.10, m |
| 20 | 154.8, qC | |
| 23 | 77.9, qC | |
| 24 | 28.1, $CH_3$ | 1.31, s |
| 25 | 28.1, $CH_3$ | 1.31, s |
| 26 | 28.1, $CH_3$ | 1.31, s |
| 27 | | 12.54, broad s |

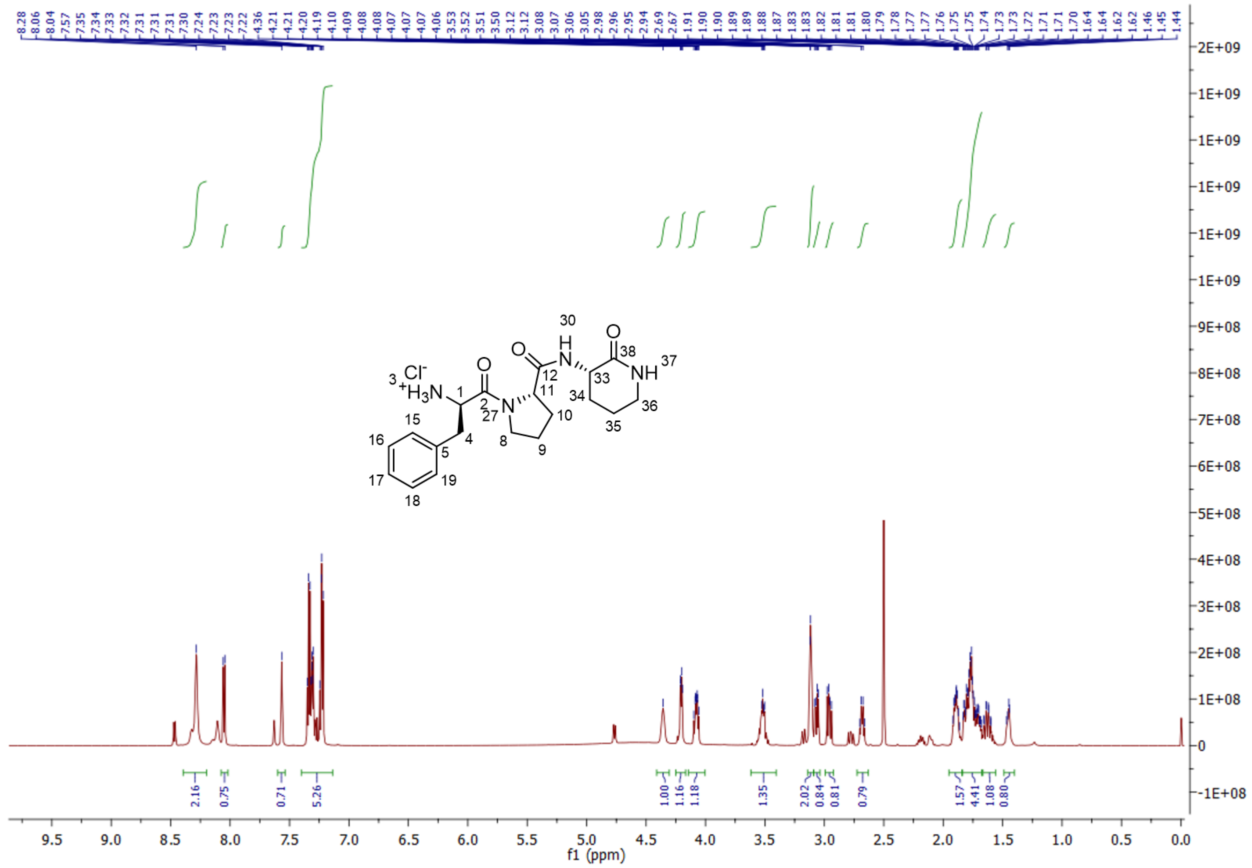

[1]H NMR of compound **8** measured in DMSO-d$_6$

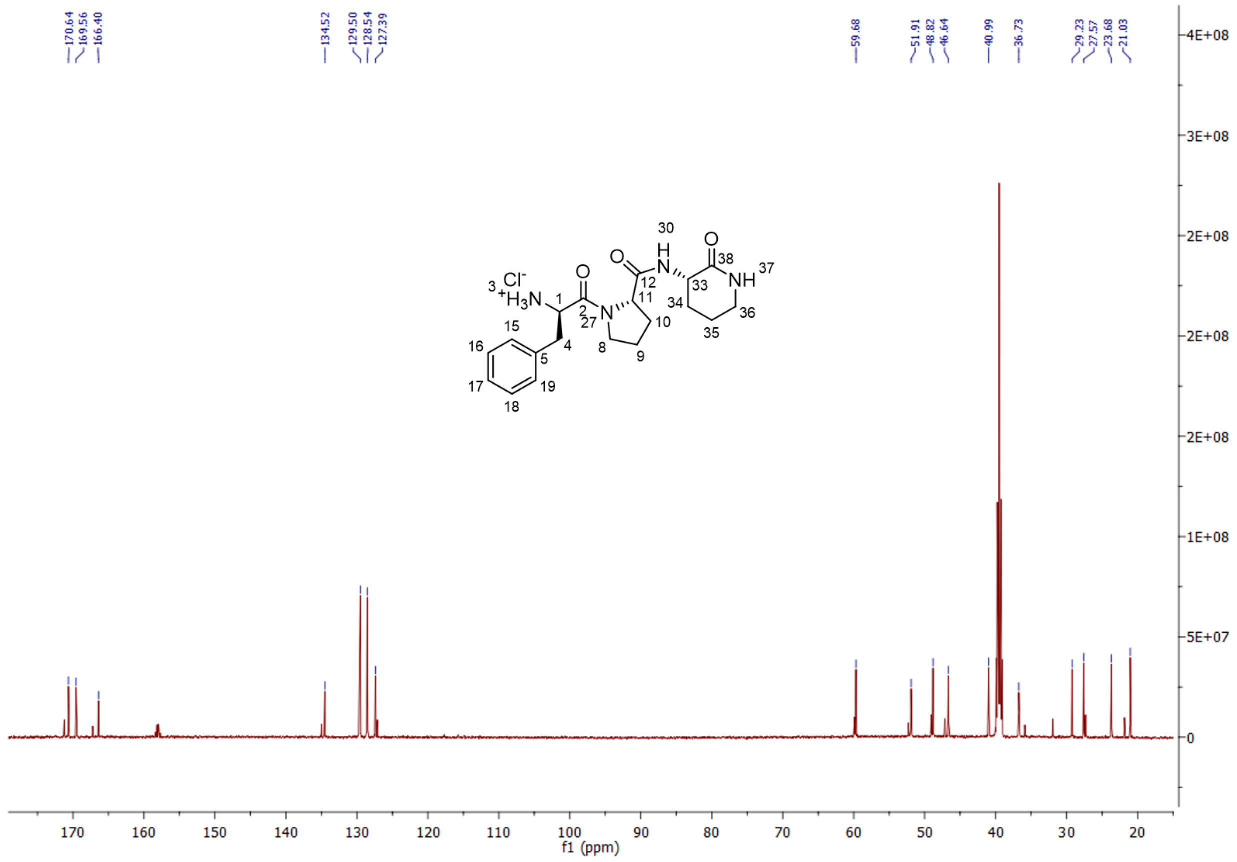

[13]C NMR of compound **8** measured in DMSO-d$_6$

| Position | $\delta_C$ | $\delta_H$, mult. (*J* in Hz) |
|---|---|---|
| 1 | 51.9, CH | 4.36, broad s |
| 2 | 170.6, qC | |
| 3 | | 8.28, broad s |
| 4a<br>4b | 36.7, $CH_2$ | 3.07, dd (13.4, 6.1)<br>2.96, dd (13.3, 8.4) |
| 5 | 134.5, qC | |
| 8a<br>8b | 46.6, $CH_2$ | 3.58 – 3.46, m<br>2.68, dd (16.8, 7.1) |
| 9a<br>9b | 23.7, $CH_2$ | 1.84 – 1.68, m<br>1.49 – 1.41, m |
| 10 | 29.2, $CH_2$ | 1.84 – 1.68, m |
| 11 | 59.7, CH | 4.21, dd (7.8, 2.8) |
| 12 | 166.4, qC | |
| 15 | 128.5, CH | 7.37 – 7.20, m |
| 16 | 129.5, CH | 7.37 – 7.20, m |
| 17 | 127.4, CH | 7.37 – 7.20, m |
| 18 | 129.5, CH | 7.37 – 7.20, m |
| 19 | 128.5, CH | 7.37 – 7.20, m |
| 30 | | 8.05, d (8.3) |
| 33 | 48.8, CH | 4.08, ddd (10.4, 8.1, 6.4) |
| 34a<br>34b | 27.6, $CH_2$ | 1.94 – 1.86, m<br>1.67 – 1.56, m |
| 35 | 21.0, $CH_2$ | 1.84 – 1.68, m |
| 36 | 41.0, $CH_2$ | 3.14 – 3.09, m |
| 37 | | 7.57, s |
| 38 | 169.6, qC | |

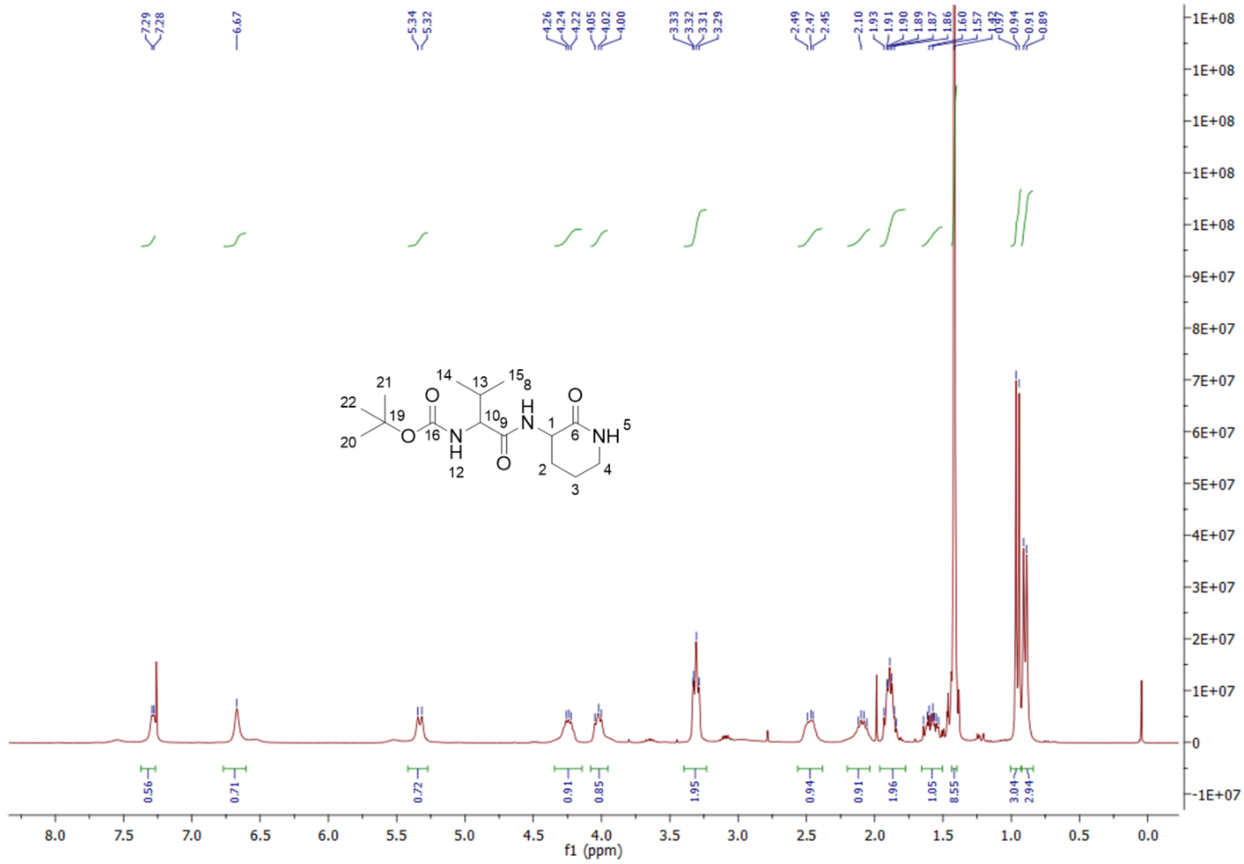

$^{1}H$ NMR of compound 10 measured in $CDCl_3$

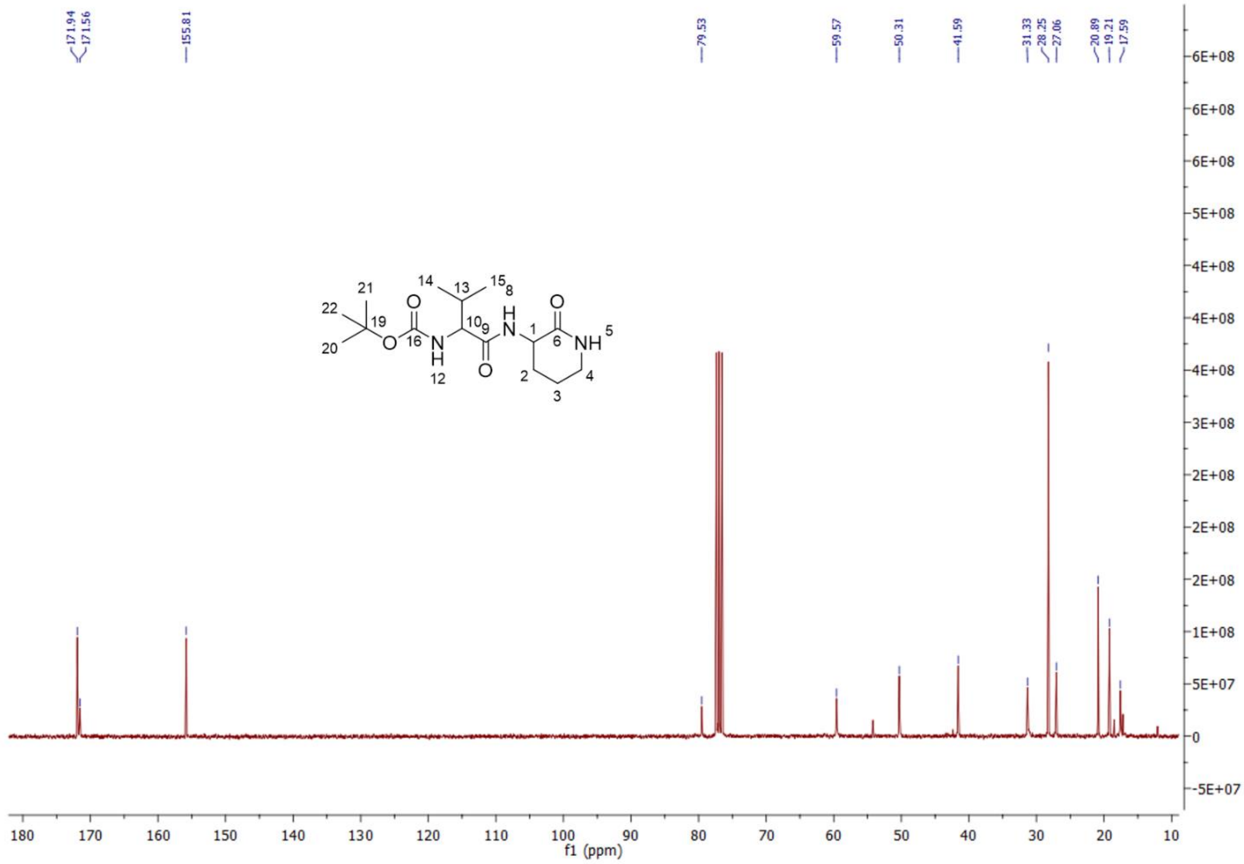

$^{13}C$ NMR of compound 10 measured in $CDCl_3$

| **Position** | **$\delta_C$** | **$\delta_H$, mult. (*J* in Hz)** |
|---|---|---|
| 1 | 50.3, CH | 4.10 – 3.96, m |
| 2a<br>2b | 27.1, $CH_2$ | 2.22 – 2.01, m<br>1.71 – 1.50, m |
| 3 | 20.9, $CH_2$ | 1.96 – 1.82, m |
| 4 | 41.6, CH | 3.40 – 3.23, m |
| 5 | | 6.67, s |
| 6 | 171.6, qC | |
| 8 | | 7.28, d (4.3 Hz) |
| 9 | 171.9, qC | |
| 10 | 59.6, CH | 4.34 – 4.18, m |
| 12 | | 5.33, d (8.6) |
| 13 | 31.3, CH | 2.58 – 2.39, m |
| 14 | 17.6, $CH_3$ | 0.90, d (6.8) |
| 15 | 19.2, $CH_3$ | 0.95, d (6.8) |
| 16 | 155.8, qC | |
| 19 | 79.5, qC | |
| 20 | 28.3, $CH_3$ | 1.42, s |
| 21 | 28.3, $CH_3$ | 1.42, s |
| 22 | 28.3, $CH_3$ | 1.42, s |

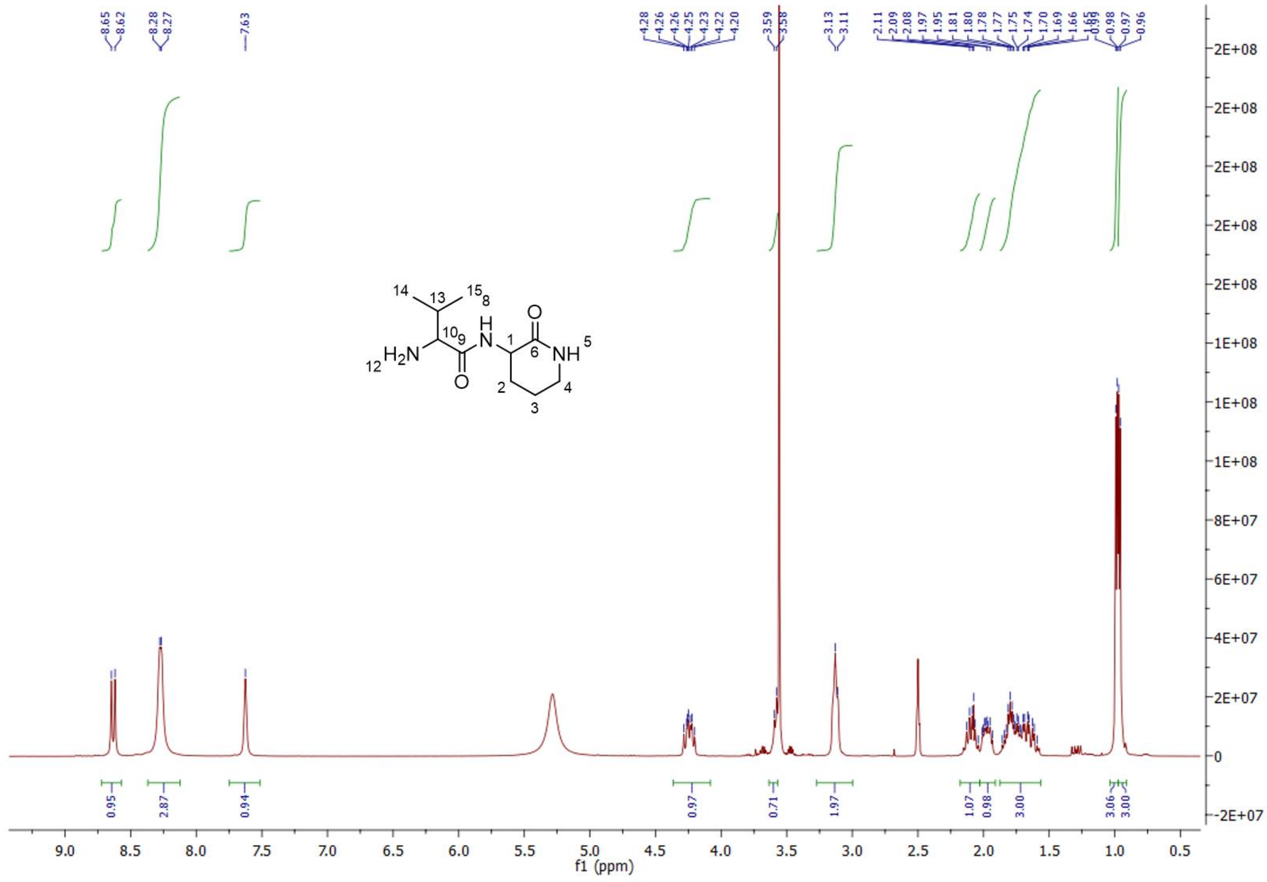

[1]H NMR of compound **11** measured in DMSO-d$_6$

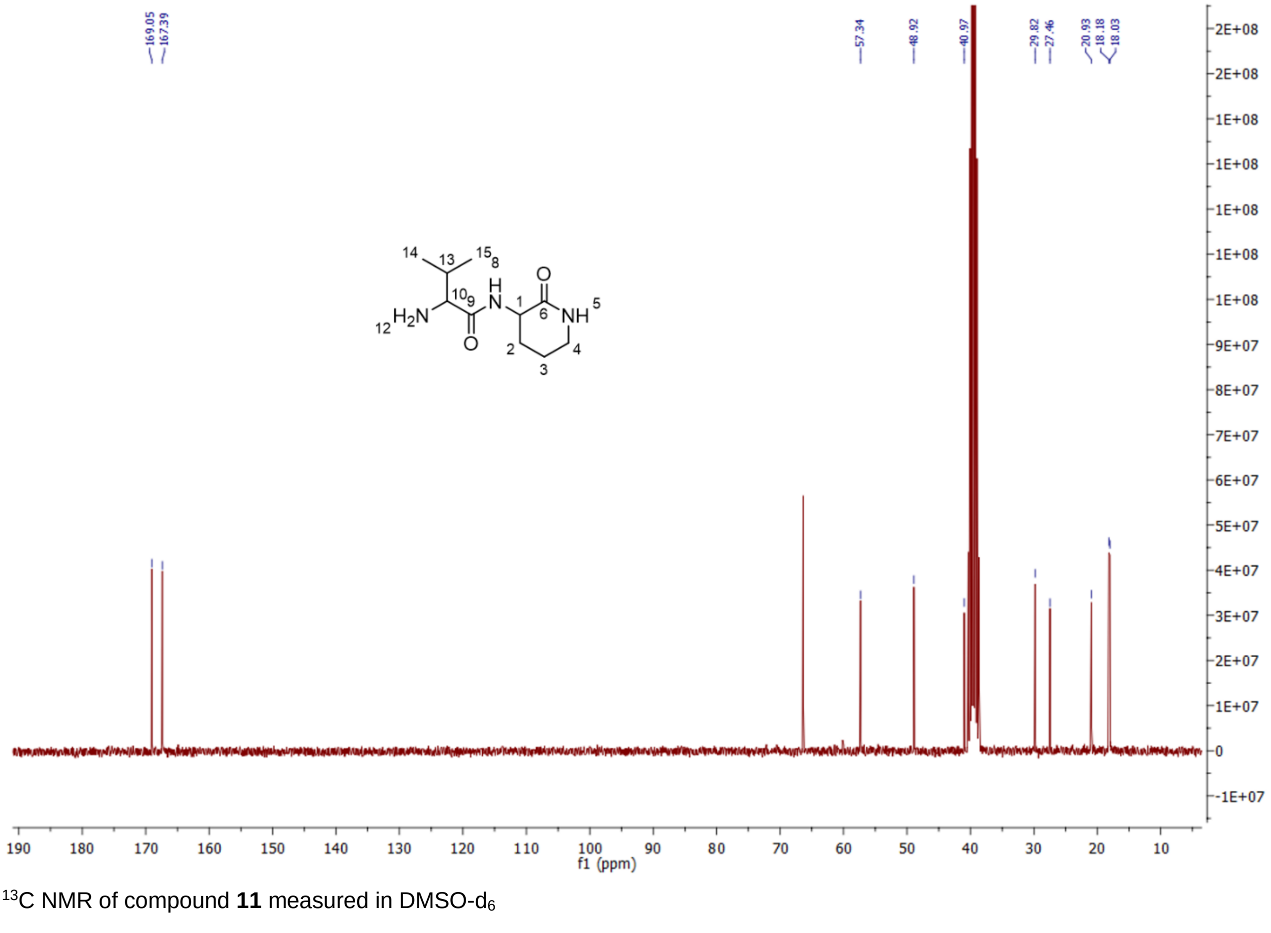

[13]C NMR of compound **11** measured in DMSO-d$_6$

| **Position** | **$\delta_C$** | **$\delta_H$, mult. (*J* in Hz)** |
|---|---|---|
| 1 | 48.9, CH | 3.64 – 3.57, m |
| 2a<br>2b | 27.5, $CH_2$ | 2.02 – 1.91, m<br>1.87 – 1.56, m |
| 3 | 20.9, $CH_2$ | 1.87 – 1.56, m |
| 4 | 41.0, $CH_2$ | 3.22 – 3.00, m |
| 5 | | 7.63, s |
| 6 | 169.1, qC | |
| 8 | | 8.63, d (8.4) |
| 9 | 167.4, qC | |
| 10 | 57.3, CH | 4.35 – 4.10, m |
| 12 | | 8.27, d (3.5) |
| 13 | 29.8, CH | 2.17 – 2.03, m |
| 14 | 18.2, $CH_3$ | 0.99, d (3.6) |
| 15 | 18.0, $CH_3$ | 0.96, d (3.6) |

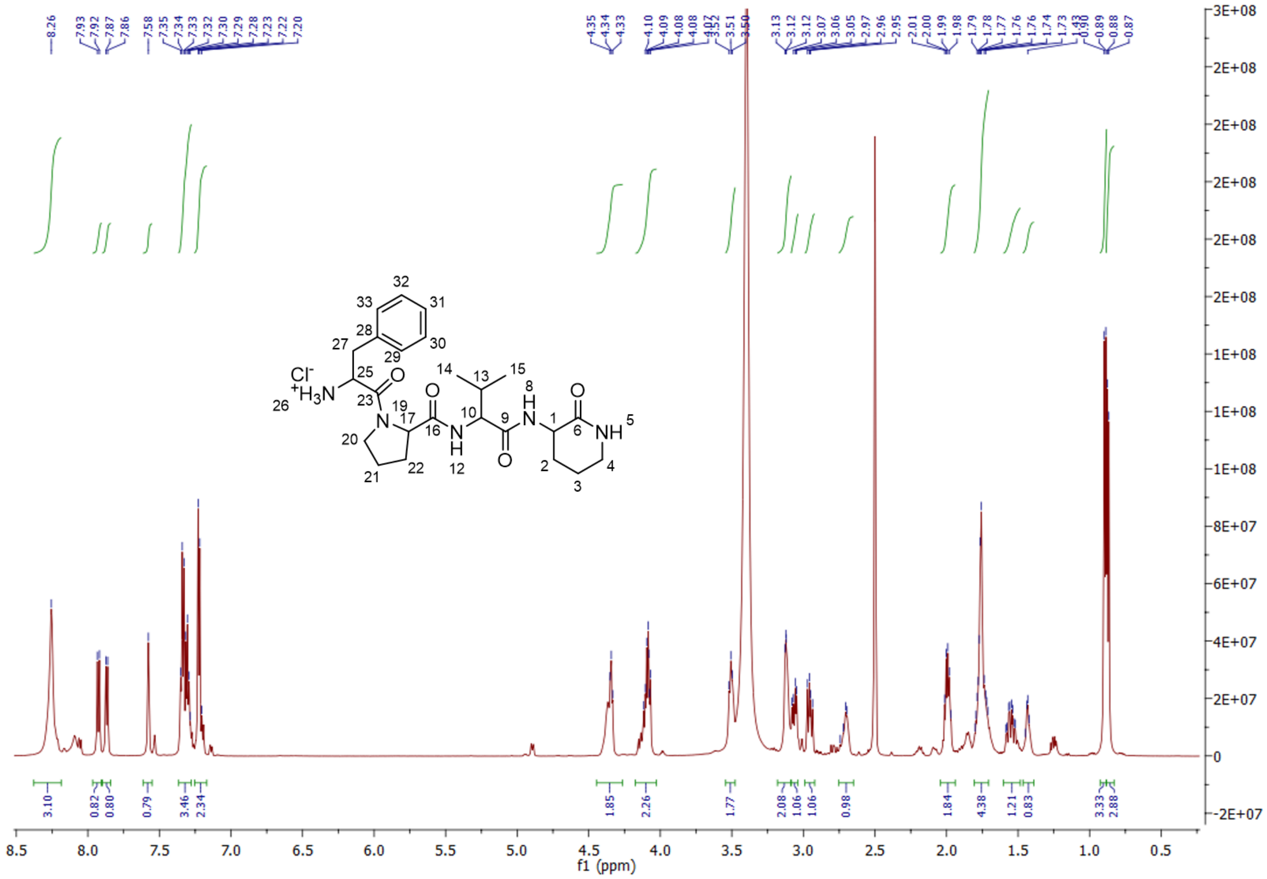

[1]H NMR of compound **13** measured in DMSO-$d_6$

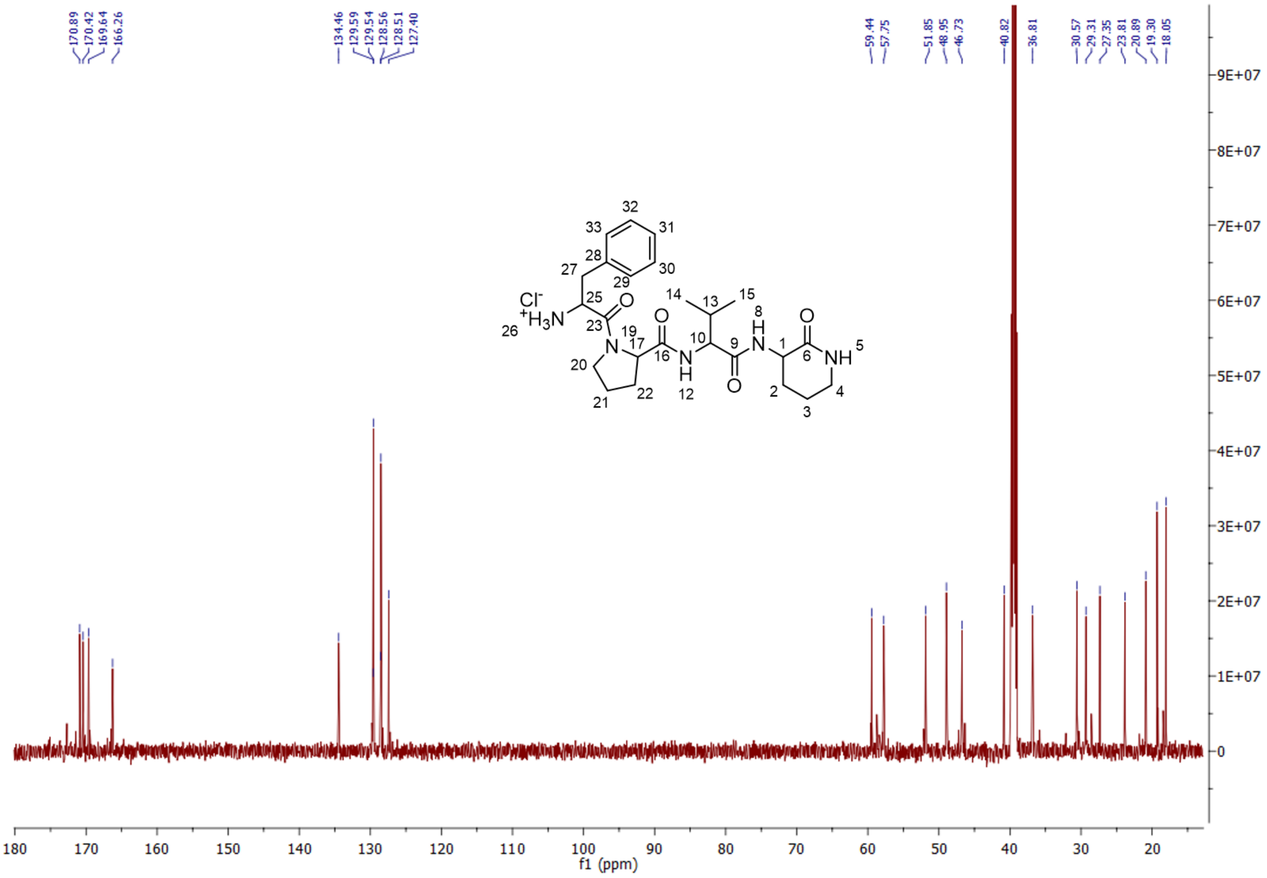

[13]C NMR of compound **13** measured in DMSO-$d_6$

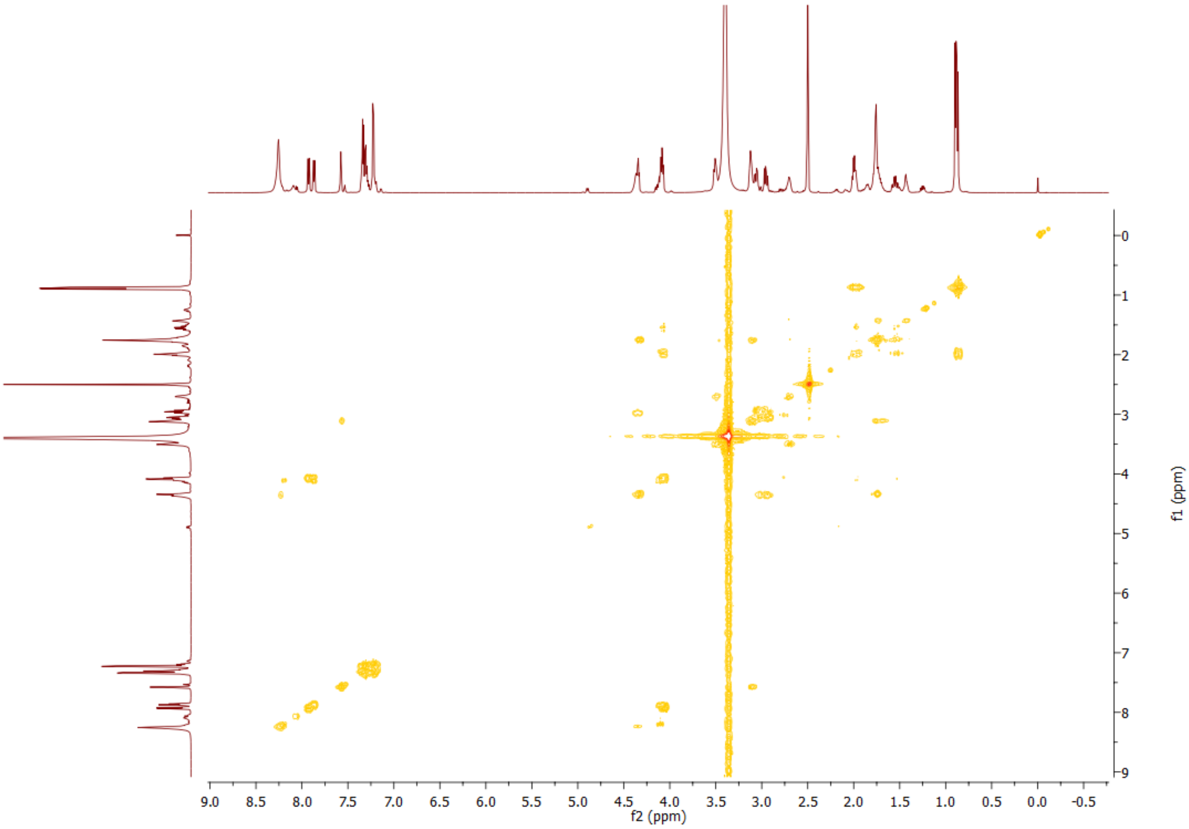

COSY spectra of compound **13** measured in DMSO-d$_6$

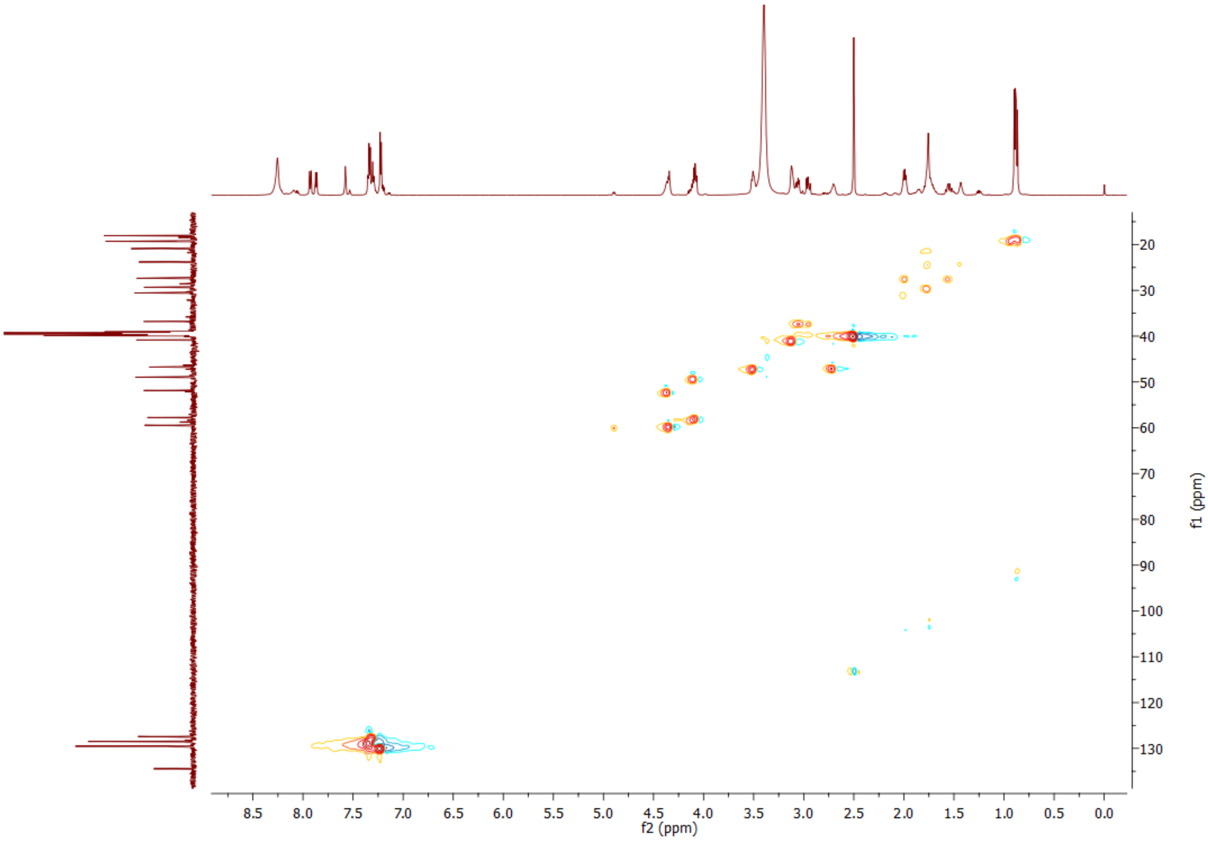

HSQC spectra of compound **13** measured in DMSO-d$_6$

| **Position** | **$\delta_C$** | **$\delta_H$, mult. (*J* in Hz)** |
|---|---|---|
| 1 | 49.0, CH | 4.17 – 4.04, m |
| 2a<br>2b | 27.4, $CH_2$ | 2.05 – 1.94, m<br>1.60 – 1.48, m |
| 3 | 20.9, $CH_2$ | 1.81 – 1.69, m |
| 4 | 40.8, $CH_2$ | 3.15 – 3.09, m |
| 5 | | 7.58, s |
| 6 | 169.6, qC | |
| 8 | | 7.87, d (7.6) |
| 9 | 170.4, qC | |
| 10 | 57.8, CH | 4.17 – 4.04, m |
| 12 | | 7.93, d (9.0) |
| 13 | 30.6, CH | 2.05 – 1.94, m |
| 14 | 19.3, $CH_3$ | 0.89, d (6.8) |
| 15 | 18.1, $CH_3$ | 0.87, d (6.8) |
| 16 | 170.9, qC | |
| 17 | 59.4, CH | 4.44 – 4.29, m |
| 20a<br>20b | 46.7, $CH_2$ | 3.58 – 3.47, m<br>2.76 – 2.64, m |
| 21a<br>21b | 23.8, $CH_2$ | 1.81 – 1.69, m<br>1.47 – 1.39, m |
| 22 | 29.3, $CH_2$ | 1.81 – 1.69, m |
| 23 | 166.3, qC | |
| 25 | 51.9, CH | 4.44 – 4.29, m |
| 26 | | 8.26, s |
| 27a<br>27b | 36.8, $CH_2$ | 3.06, dd (13.3, 6.0)<br>2.95, dd (13.3, 8.3) |
| 28 | 134.5, qC | |
| 29 | 128.5, CH | 7.25 – 7.18, m |
| 30 | 129.5, CH | 7.37 – 7.28, m |
| 31 | 127.4, CH | 7.37 – 7.28, m |
| 32 | 129.5, CH | 7.37 – 7.28, m |
| 33 | 128.5, CH | 7.25 – 7.18, m |

### References

[1] S. Gruenewald, H. D. Mootz, P. Stehmeier, T. Stachelhaus, *Appl. Environ. Microbiol.* **2004**, *70*, 3282–3291.

[2] H. Kries, R. Wachtel, A. Pabst, B. Wanner, D. Niquille, D. Hilvert, *Angew Chem Int Ed Engl* **2014**, *53*, 10105–10108.

[3] D. L. Niquille, D. A. Hansen, T. Mori, D. Fercher, H. Kries, D. Hilvert, *Nat. Chem.* **2018**, *10*, 282–287.

[4] H. A. Greisman, C. O. Pabo, *Science* **1997**, *275*, 657–661.

[5] X. Cai, S. Nowak, F. Wesche, I. Bischoff, M. Kaiser, R. Fürst, H. B. Bode, *Nat. Chem.* **2017**, *9*, 379–386.

[6] C. Fu, W. P. Donovan, O. Shikapwashya-Hasser, X. Ye, R. H. Cole, *PLoS One* **2014**, *9*, e115318.

[7] C. T. Chung, S. L. Niemela, R. H. Miller, *Proc Natl Acad Sci U S A* **1989**, *86*, 2172–2175.

[8] D. Jantz, J. M. Berg, *Biophys. J.* **2010**, *98*, 852–860.

[9] E. Krieger, G. Vriend, *Bioinformatics* **2014**, *30*, 2981–2982.

[10] M. Biasini, S. Bienert, A. Waterhouse, K. Arnold, G. Studer, T. Schmidt, F. Kiefer, T. G. Cassarino, M. Bertoni, L. Bordoli, et al., *Nucleic Acids Res.* **2014**, *42*, W252-8.

[11] Schrodinger LLC, **2010**.

[12] C. Hacker, X. Cai, C. Kegler, L. Zhao, A. K. Weickhmann, J. P. Wurm, H. B. Bode, J. Wöhnert, *Nat. Commun.* **2018**, *9*, 4366.

[13] M. Hahn, T. Stachelhaus, *Proc. Natl. Acad. Sci. USA* **2006**, *103*, 275–280.
